# Multiomic Characterization of Stage I Lung Adenocarcinoma Reveals Distinct Genetic and Immunologic Features of Recurrent Disease

**DOI:** 10.1101/2021.05.20.444915

**Authors:** Johannes R. Kratz, Jack Z. Li, Jessica Tsui, Jen C. Lee, Vivianne W. Ding, Arjun A. Rao, Michael J. Mann, Vincent Chan, Alexis J. Combes, Matthew F. Krummel, David M. Jablons

**Affiliations:** Department of Surgery, University of California San Francisco, San Francisco, CA 94143, USA; Thoracic Oncology Laboratory, University of California San Francisco, San Francisco, CA 94143, USA; Helen Diller Family Comprehensive Cancer Center, University of California San Francisco, San Francisco, CA 94143, USA; ImmunoX Initiative, University of California San Francisco, San Francisco, CA 94143, USA; Department of Pathology, University of California San Francisco, San Francisco, CA 94143, USA; UCSF CoLabs, University of California San Francisco, San Francisco, CA 94143, USA; Department of Microbiology and Immunology, University of California San Francisco, San Francisco, CA 94143, USA

## Abstract

**Background:** Recurrence after surgery for early-stage lung cancer is common, occurring between 30-50% of the time. Despite the popularization of prognostic gene signatures in early-stage lung cancer that allow us to better predict which patients may recur, why patients recur after surgery remains unclear.

**Methods:** Using a large cohort of lung adenocarcinoma patients with complete genetic, genomic, epigenetic and clinical profiling, a recurrence classifier was developed which identifies patients at highest risk of recurrence. The genetic, genomic, and epigenetic profiles of stage I patients with low-vs. high-risk of recurrence were compared. To characterize the tumor immune microenvironment of recurrent stage I tumors, single cell RNA-seq was performed on fresh tissue samples undergoing lung adenocarcinoma resection at UCSF to identify unique immune population markers and applied to the large stage I lung adenocarcinoma cohort using digital cytometry.

**Results:** Recurrence high-risk stage I lung adenocarcinomas demonstrated a higher mutation burden than low-risk tumors, however, none of the known canonical lung cancer driver mutations were more prevalent in high-risk tumors. Transcriptomic analysis revealed widespread activation of known cancer and cell cycle pathways with simultaneous downregulation of immune response pathways including antigen presentation and Th1/Th2 activation. Tumors at high-risk of recurrence displayed depleted adaptive immune populations, and depletion of adaptive immune populations was independently prognostic of recurrence in stage I lung adenocarcinomas.

**Conclusion:** Recurrent stage I lung adenocarcinomas display distinct features of genomic and genetic instability including increased tumor mutation burden, neoantigen load, activation of numerous mitotic and cell cycle genes, and decreased genome-wide methylation burden. Relative depletion of infiltrating adaptive immune populations may allow these tumors to escape immunosurveillance and recur after surgery.

## Background

Over 1.8 million people are diagnosed with lung cancer worldwide annually, resulting in 1.6 million deaths each year^1^. Although early detection has been shown to improve survival in other cancers, early detection of lung cancer is not enough. Even after complete surgical resection, early-stage lung cancers recur up to 30-50% of the time within 5 years^2^, causing a staggering number of deaths worldwide. Despite these overwhelming recurrence rates, the current standard of care for most patients who undergo surgery for early-stage lung cancer is “observation”. Chemotherapy after surgery does not improve survival for the majority of these patients^3–7^.

In order to predict which patients recur after surgery, multiple groups have identified molecular signatures that more accurately predict recurrence and mortality beyond conventional TNM staging^2, 8–29^. These signatures are based on a variety of genetic, genomic, and epigenetic biomarkers prognostic of recurrence^2, 8–29^. Despite an improved ability to predict who may recur based on these biomarkers, however, the biology of why these patients recur after surgery is less well understood. While multiple studies have described the genetic, genomic, and epigenetic alterations in tumor vs. normal tissue^30, 31^, only a handful of groups have investigated the molecular biology of early-stage lung cancer recurrence^32, 33^. These studies have primarily focused on mutation profiles associated with recurrence by utilizing targeted NGS panels of known cancer mutations^32, 33^. A more comprehensive account of the genomic and epigenetic molecular features of recurrence, however, has not been described.

The relationship between the tumor immune microenvironment and early-stage lung cancer recurrence after surgery also remains poorly understood. Advances in cancer immunotherapy have highlighted the importance of the immune system in keeping tumor progression and metastasis in check. Several cancer-immune landscapes have been described: 1. inflamed or “hot” tumors, infiltrated by exhausted and ineffective immune cells, 2. immune-desert or “cold” tumors, which lack appropriate T-cell activation and priming, and 3. immune-excluded tumors, in which vascular or stromal barriers prevent immune infiltration^34^. Precisely defining the cancer-immune landscape of recurrent early-stage lung cancer has important clinical implications. Multiple studies, for example, have demonstrated that checkpoint inhibitors are primarily effective in “hot” tumor landscapes with a large number of infiltrating but dormant T-cells^34^. In fact, the presence of alternative “cold” and “excluded” tumor landscapes may explain why a durable clinical response to checkpoint inhibitors is seen in only approximately 20% of patients^34^. To address this, several groups have assessed the prognostic significance of tumor infiltrating leukocytes in lung adenocarcinomas using immunohistochemistry with a pre-defined set of immune cell surface markers and cytokines^35, 36^ or by using computational immune profiling techniques^37, 38^. These groups, however, have only assessed small focused panels of immune cell types^35, 36^ or used pre-defined sets of immune cell markers not derived from tumor infiltrating leukocytes to perform computational profiling^37, 38^. More comprehensive and specific descriptions of the tumor immune microenvironment characteristic of early recurrence after surgery are still needed to focus our current immunotherapy treatments and reveal novel immunotherapy targets.

This study investigates the molecular and immunologic features of early recurrence in stage I lung adenocarcinoma using a large cohort of patients with complete genomic, genetic, epigenetic, and clinical profiles. scRNA-seq analysis is used to develop unique early-stage lung adenocarcinoma immune population profiles that allow for comprehensive characterization of immune populations in this cohort. Our analysis reveals distinct genetic, genomic, epigenetic, and immunologic differences between recurrent and non-recurrent early-stage lung adenocarcinomas and identifies several novel biomarkers and therapeutic targets associated with early recurrence.

## Methods

### Patient Cohorts

Tumor and normal samples from 522 patients in the TCGA lung adenocarcinoma cohort^39^ were included in this study. Indexed clinical data were downloaded from the Genomic Data Commons (GDC) portal (TCGAbiolinks v2.18^40, 41^). Clinical outcomes including survival and recurrence endpoints were obtained from the TCGA Clinical Data Resource^42^.

To perform tumor immunoprofiling, additional tumor and adjacent normal tissue samples from six patients who underwent complete surgical resection of pathologic stage I lung adenocarcinoma via lobectomy and mediastinal lymph node dissection at UCSF between 2019 and 2020 were characterized. None of these patients underwent neoadjuvant therapy prior to resection and sample collection. All patients gave informed consent for sample collection. The study was approved by the UCSF Institutional Review Board (IRB# 11-06107).

### Recurrence Score

Tumor and normal RNA-seq TOIL recompute counts^43^ from the TCGA Pan-Cancer (PANCAN) cohort were downloaded from the UCSC Xena browser^44^. Gene counts-per-million (CPM) for each tumor and normal sample were generated using edgeR^45^. To create a risk score associated with recurrence, a combination of *L1*- and *L2*-penalized Cox proportional hazards modeling (R package glmnet v4.1.1) was used with progression-free interval (PFI.1 in the TCGA Clinical Data Resource^42^) as an endpoint. Using 10-fold cross-validation, the amount of *L1*-penalization was increased until a parsimonious set of genes that maximized the concordance index was obtained (Supplemental Table S1). *L2*-penalization was simultaneously applied in order to reduce model overfitting as much as possible. A continuous recurrence score was generated for each tumor sample by summing the product of CPM values for each model gene by cox proportional hazards model coefficients. Resultant predicted risk scores were divided at the 25^th^, 50^th^, and 75^th^ percentiles to generate low-, low-intermediate-, intermediate-high, and high-risk groups.

### Genomic Alteration Analysis

Mutation Annotation Format (MAF) files were downloaded from the Genomic Data Portal (R package TCGAbiolinks v2.18^40, 41^). The total number of mutations was summed across all surveyed genes for each sample (R package maftools v2.7.10^46^). Recurrence risk category and type of mutation were overlaid on the most frequently mutated genes and known lung cancer driver mutations to generate combined oncoplots. Differentially mutated genes were detected by performing Fisher exact tests on all genes between recurrence low- and recurrence high-risk cohorts. To analyze copy number alterations in each patient tumor sample, sample level copy number alteration data were downloaded from the Broad GDAC Firehose pipeline (R package RTCGAToolbox v2.20.0^47^). To analyzed genome-wide differences in copy number alterations across recurrence low-vs. high-risk cohorts, Affymetrix Genome-Wide Human SNP Array 6.0 data were downloaded from the GDC portal. Significant recurrent copy number alterations were identified using Genomic Analysis of Important Alterations (GAIA)^48^. Taking within-sample homogeneity into account, GAIA performs a conservative permutation test to extract the regions most likely involved in functional phenotype changes. Recurrent amplifications and deletions were identified for each recurrence risk cohort and annotated; genomic regions identified as significantly altered in copy number (adjusted *P-*value < 10^−4^) were represented on chromosome overview plots.

### Transcriptome and Methylation Analysis

Differential gene expression between recurrence low-vs. high-risk cohorts was performed with TOIL recompute counts (described above) using edgeR^45^, limma^49^, and voom^50^. A heatmap of differential gene expression was produced using the ComplexHeatmap package in R (v2.6.2). Partitioning Around Medoids (PAM) (R package cluster v2.1.0) was used to partition genes into distinct genomic clusters. Clinical covariates including recurrence risk category, age, stage, gender, and smoking history were used to annotate the heatmap. Conventional pathway analysis on the top differentially expressed genes in each cluster was performed using IPA (Qiagen, Germantown, PA). Topological pathway analysis was performed on the top differentially expressed genes using the R package SPIA (v2.42.0).

For methylation analysis, Illumina Human Methylation 450 array data aligned to homo sapiens (human) genome assembly GRCh37 (hg19) was downloaded from the Genomic Data Portal (R package TCGAbiolinks v2.18^40, 41^). Methylation data was annotated and filtered to remove unmapped probes, probes mapped to multiple places in the genome^51^, and duplicate probes as previously described^52^. Differential gene methylation between recurrence low-vs. high-risk cohorts was performed with M-values using edgeR^45^ and limma^49^. A heatmap of differential gene methylation was produced using the ComplexHeatmap package in R (v2.6.2). Partitioning Around Medoids (PAM) (R package cluster v2.1.0) was used to partition methylated genes into distinct genomic clusters. Clinical covariates including recurrence risk category, age, stage, gender, and smoking history were used to annotate the heatmap. Differentially methylated regions were defined and mapped to the chromosome as previously described^52^. Conventional pathway analysis on the top differentially methylated genes was performed using GO, KEGG, and GSEA Molecular Signatures Database Hallmark Pathways as previously described^52^. Upstream regulation of gene enhancers by transcription factors was analyzed by integration of annotated methylation and gene expression data using the ELMER R package (v3.12^53^).

### Tumor Immunoprofiling

Freshly resected tumor or adjacent normal lung tissue was thoroughly chopped with surgical scissors and transferred to GentleMACS C Tubes (Miltenyi Biotec) containing 20 uL/mL Liberase TL (5 mg/ml, Roche) and 50 U/ml DNAse I (Roche) in Leibovitz’s L-15 per 0.3 g tissue. GentleMACS C Tubes were then installed onto the GentleMACS Octo Dissociator (Miltenyi Biotec) and incubated for 45min according to the manufacturer’s instructions. Samples were quenched with 15 mL of sort buffer (PBS/2% FCS/2mM EDTA), filtered through 100 um filters and spun down. Red blood cell lysis was performed with 175 mM ammonium chloride if needed. Cells were subsequently incubated with with Zombie Aqua Fixable Viability Dye (Thermo), washed with sort buffer, and incubated with Human FcX (BioLegend) to prevent non-specific antibody binding. Following FcX incubation, cells were washed with sort buffer and incubated with cell surface antibodies mix diluted in the BV stain buffer (BD Biosciences) following manufacturer instructions for 30 minutes on ice in the dark. Live cells were sorted from these single cell suspensions on a BD FACSAria Fusion. After sorting, cells were pelleted and resuspended at 1×10^3^ cells/ml in 0.04% BSA/PBS and loaded onto the Chromium Controller (10X Genomics). Samples were processed for single-cell encapsulation and cDNA library generation using the Chromium Single Cell 3’ v3 Reagent Kits (10X Genomics). In order to duplex tumor and normal cells, cell hashing was performed using TotalSeq-A antibodies (BioLegend) according to manufacturer instructions. Libraries were subsequently sequenced on an Illumina HiSeq 4000 (Illumina).

Sequences from scRNA-seq were processed with Cellranger (10x Genomics) and read into the Seurat R package. The filtered count matrices were normalized and variance stabilized using negative binomial regression via the scTransform method^54^. Mitochondrial content, ribosomal content, and cell cycle state were regressed from the normalized data. Datasets from twelve samples (six tumor and six normal) were integrated using the anchoring method described by Stuart et al.^55^. The integrated matrix underwent dimensionality reduction using Principal Component Analyses (PCA); the first 40 principal coordinates were subjected to non-linear dimensionality reduction using Uniform Manifold Approximation and Projection (UMAP). Clusters of cells sharing similar transcriptomic profiles were identified using the Louvain algorithm with a clustering resolution of 0.8. Differentially expressed genes for each cluster were identified using the Wilcox test in Seurat and top cluster genes (log fold change >0.4) were used to identify cell clusters.

Measurements of immune cell population densities was performed using the digital cytometry method described by Newman et al.^56^. This method utilizes single-cell reference profiles to calculate cell type frequencies from bulk transcriptome data. 10,000 labeled cells from the scRNA-seq experiments above were used to create a single cell reference matrix. Cell fractions were imputed from RNA-seq data from the entire TCGA lung adenocarcinoma cohort in absolute mode with quantile normalization disabled. To cluster tumors by their immune profiles, a combination of *L1*- and *L2*-penalized Cox proportional hazards modeling (R package glmnet v4.1.1) was used as described above with immune cell population densities as predictors and recurrence as an endpoint. Using 10-fold cross-validation, unbiased increase in the amount of *L1*-penalization resulted in a parsimonious set of immune cell population densities predictive of recurrence. The immune cell populations predictive of recurrence using this method were enriched for adaptive immune populations. Adaptive immune cell scores were generated for each tumor sample by summing the product of density values for each immune cell population in the model by cox proportional hazards model coefficients, resulting in adaptive-depleted, adaptive-low, adaptive-neutral, and adaptive-rich clusters when separated at the 25^th^, 50^th^, and 75^th^ percentiles.

### Statistical Analysis

The primary endpoint in this study was recurrence (PFI.1 in the TCGA Clinical Data Resource^42^). The primary predictors evaluated were recurrence risk score, recurrence risk category, age, sex, smoking history, and stage. These predictors were correlated with recurrence using univariate and multivariate Cox proportional hazards modeling. Wald tests were performed to evaluate for statistical significance in Cox proportional hazards modeling. Stratified Kaplan-Meier analysis using a right-censored dataset and the log-rank test for trend were used to evaluate the correlation between primary predictors and the primary endpoint. Time-dependent Area Under the Receiver Operating Characteristic curve (AUROC) was calculated using the survcomp (v1.4.0) package in R. Differences in AUROCs were tested by multivariate Cox proportional hazards modeling, integrated AUROCs were compared by Wilcoxon rank sum test. For all statistical tests, a pre-specified two-sided α of 0.05 was considered significant. Adjusted *P*-values are reported for all analyses involving multiple comparisons. Analyses were conducted using the programming languages R^57^ (v4.0.3 for Macintosh) and Prism (v9.1.0, GraphPad Software, San Diego, CA).

## Results

### Recurrence Score

In order to incorporate time-to-recurrence data, a score that predicts the risk of recurrence for each patient in the TCGA lung adenocarcinoma was developed. This recurrence score relates each patient’s gene expression profile to both recurrence as well as time-to-recurrence. Elastic net penalization using cross-validation yielded a parsimonious set of genes associated with recurrence in 500 patients with stage I-IV lung adenocarcinoma in the TCGA cohort (Supplemental Table S1). Weighting and combining the expression of these risk classifier genes by coefficients assigned by elastic net penalization (Supplemental Table S1) resulted in a recurrence score for each patient in the TCGA lung adenocarcinoma cohort. Higher recurrence scores were directly proportional to the probability of recurrence within 5 years of diagnosis (Figure 1A). Grouping patients into cohorts by recurrence score quartile in the stage I-IV TCGA lung adenocarcinoma cohort resulted in significantly different recurrence rates by Kaplan-Meier analysis (5-year Freedom From Recurrence of 68.9%, 39.5%, 34.8%, 25.0% for low-, low-intermediate, intermediate-high, and high-risk groups respectively, *P* < 0.001). The TCGA lung adenocarcinoma cohort includes patients who presented with nodal and distant organ spread. In order to study the biology of recurrence in patients not known to have disease spread, we limited further analysis to stage I patients only. Grouping stage I lung adenocarcinoma patients into cohorts by recurrence score quartile identified a high-risk cohort with 57.6% incidence of recurrence within 2 years of surgical resection as opposed 3.1% in the low-risk cohort (Figure 1B, 2-year Freedom From Recurrence 96.9%, 84.5%, 69.6%, 42.4% for low-, low-intermediate, intermediate-high, and high-risk groups respectively, *P* < 0.001). By univariate analysis, recurrence risk category and stage were statistically significant predictors of recurrence (Table 1). Adjusting for age, smoking history, and stage, only recurrence risk category (HR 2.09, 95% CI 1.54-2.84, *P*<0.001) and female sex (HR 0.67, 95% CI 0.49-0.92, *P*=0.012) were significant predictors of recurrence by multivariate analysis (Table 1).

**Table 1.** Cox Proportional Hazards Models for 5-year Freedom From Recurrence.

|  | Univariate Analysis |  |  | Multivariate Analysis |  |  |
| --- | --- | --- | --- | --- | --- | --- |
|  | HR | 95% CI | Wald test *P* value | HR | 95% CI | Wald test *P* value |
| **Risk Category\*** | **2.01** | **1.59-2.52** | **<0.001** | **2.09** | **1.54-2.84** | **<0.001** |
| Age >65 | 0.92 | 0.58-1.46 | 0.731 | 1.01 | 0.58-1.75 | 0.976 |
| Smoking History§ | 1.20 | 0.86-1.68 | 0.292 | 0.99 | 0.70-1.38 | 0.937 |
| Female Sex | 0.97 | 0.61-1.55 | 0.914 | **0.67** | **0.49-0.92** | **0.012** |
| **Stage** | **1.61** | **1.01-2.56** | **0.045** | 1.28 | 0.72-2.29 | 0.402 |

\*Compared to Low Risk Category
§ <5, 6-20, 21-40, >40 Pack-Years

**Figure 1.**
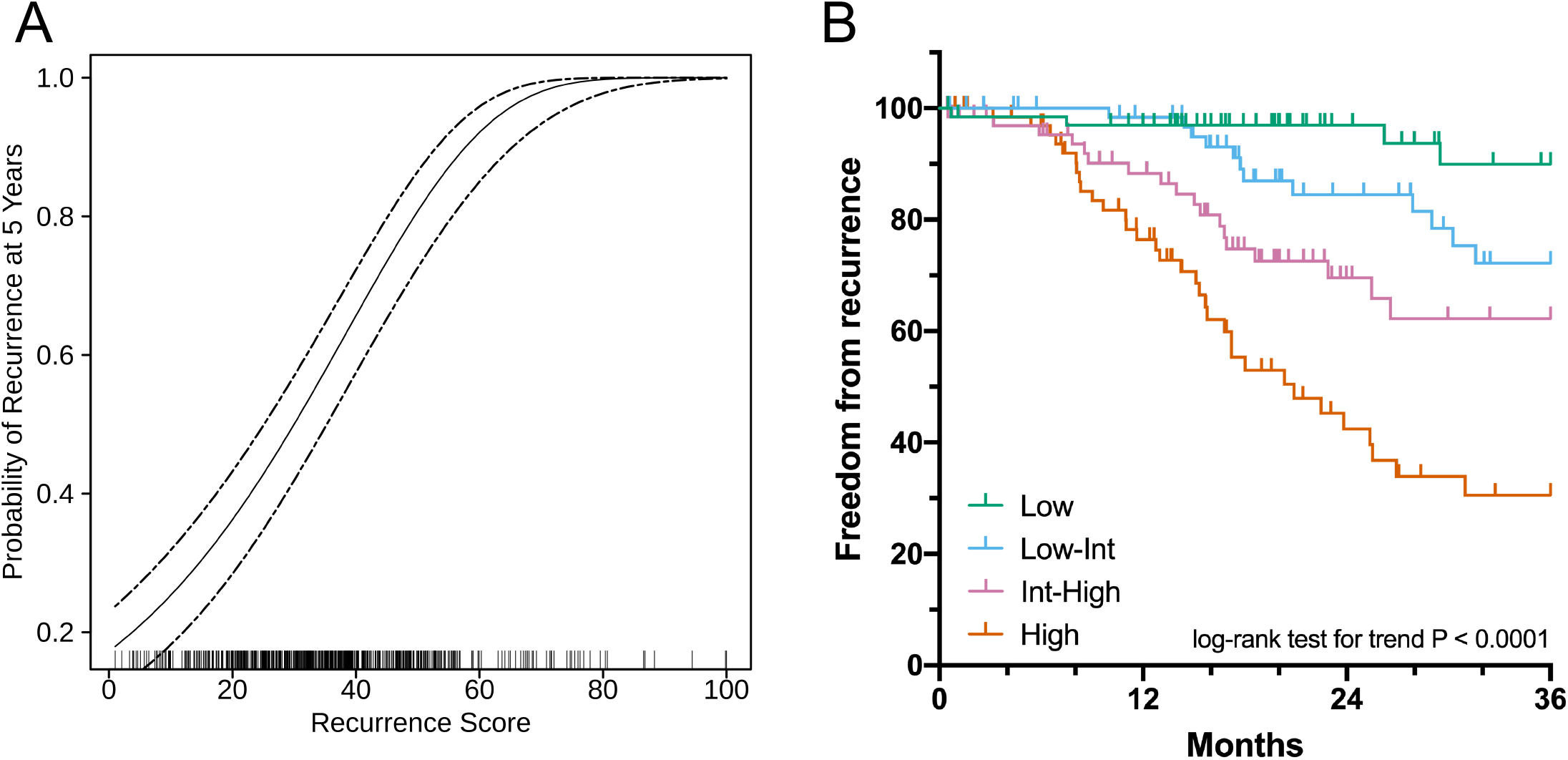
Recurrence score is prognostic of recurrence in the TCGA lung adenocarcinoma cohort. Recurrence score is directly proportional to probability of recurrence at 5 years within the entire TCGA lung adenocarcinoma cohort (A) and is prognostic of recurrence when stage I patients are divided into cohorts by recurrence score quartile (B).

Conventionally, TNM stage is used to predict risk of recurrence. In order to compare risk predictions using our risk categorization method vs. TNM stage in this cohort, we performed time-dependent area under the receiver operating characteristic curve (AUROC) analysis (Supplemental Figure 1A). The recurrence score better predicted risk of recurrence, increasing the time-dependent AUROC to 0.714 from 0.584 vs. stage alone (*P* < 0.001).

### Genetic Alterations

The median number of tumor mutations observed in the TCGA stage I lung adenocarcinoma cohort was 176. When assessing tumor mutation burden by recurrence risk category in this cohort, a positive association between tumor mutation burden and risk of recurrence was observed (median tumor mutations of 81, 158, 233, and 274 in recurrent low-, low-intermediate, intermediate-high, and high-risk tumor phenotypes respectively [*P* < 0.001], Figure 2A). *TP53* was most frequently mutated in the stage I cohort; 47% of samples had a *TP53* mutation (Figure 2C). Other frequently mutated genes included *TTN, MUC16, CSMD3, RYR2, LRP1B, ZFHX4, USH2A, KRAS*, and *FLG* (Figure 2C). 23 genes were mutated significantly more frequently in high-vs. low-risk stage I tumors, including *TP53, TTN, SLC8A1, AHNAK, KCNU1, COLA5A2, COL22A1, PKHD1L1, SMARCA4*, and *ZFHX4* (Table 2). Canonical lung cancer driver mutations were also assessed. Although *KRAS* mutations were observed more frequently in high-risk tumors (30% vs. 19% in low-risk tumors, adjusted *P* = 0.31) and *EGFR* mutations were observed more frequently in low-risk tumors (18% vs. 4% in high-risk tumors, adjusted *P* = 0.13), these differences were not statistically significant. In fact, no significant differences in canonical lung cancer driver mutations were observed between low- and high-risk tumors (Figure 2B). Similar to tumor mutation burden, copy number alterations were significantly increased in recurrence high-vs. low-risk tumors (median relative CNA of 1075 vs. 1462 in low-vs. high-risk categories respectively [*P* < 0.001], Figure 2D). Although copy number alterations and mutations did not show any predilection for a particular chromosome in high-risk tumors (Figure 2E), different patterns of copy number alterations were observed between low- and high-risk tumors across the genome (Figure 2F).

**Table 2.** Genes more frequenlty mutated in recurrence high-vs. low-risk stage I lung adenocarcinomas.

| Hugo_Symbol | # High Risk<br>Mutated Samples | # Low Risk<br>Mutated Samples | P-value | Odds<br>Ratio | 95%<br>Upper CI | 95%<br>Lower CI | Adj P-value |
| --- | --- | --- | --- | --- | --- | --- | --- |
| TP53 | 46 | 16 | 0 | 6.866 | 16.123 | 3.059 | 0 |
| TTN | 39 | 12 | 0 | 6.286 | 15.403 | 2.722 | 0.001 |
| SLC8A1 | 14 | 0 | 0 | Inf | Inf | 3.92 | 0.021 |
| AHNAK | 13 | 0 | 0 | Inf | Inf | 3.54 | 0.028 |
| KCNU1 | 13 | 0 | 0 | Inf | Inf | 3.54 | 0.028 |
| COL5A2 | 15 | 1 | 0 | 18.728 | 810.111 | 2.713 | 0.031 |
| COL22A1 | 12 | 0 | 0 | Inf | Inf | 3.174 | 0.031 |
| PKHD1L1 | 12 | 0 | 0 | Inf | Inf | 3.174 | 0.031 |
| SMARCA4 | 12 | 0 | 0 | Inf | Inf | 3.174 | 0.031 |
| ZFHX4 | 31 | 11 | 0 | 4.334 | 10.836 | 1.844 | 0.031 |
| CSMD3 | 37 | 16 | 0 | 3.889 | 8.86 | 1.769 | 0.031 |
| ADAMTS12 | 22 | 5 | 0 | 5.984 | 21.784 | 2.01 | 0.031 |
| NRXN1 | 22 | 5 | 0 | 5.984 | 21.784 | 2.01 | 0.031 |
| CACNA1E | 16 | 2 | 0.001 | 10.046 | 94.084 | 2.204 | 0.031 |
| FAT4 | 16 | 2 | 0.001 | 10.046 | 94.084 | 2.204 | 0.031 |
| XIRP2 | 26 | 8 | 0.001 | 4.623 | 13.044 | 1.81 | 0.031 |
| CDH18 | 11 | 0 | 0.001 | Inf | Inf | 2.82 | 0.031 |
| CTNND2 | 11 | 0 | 0.001 | Inf | Inf | 2.82 | 0.031 |
| DMXL1 | 11 | 0 | 0.001 | Inf | Inf | 2.82 | 0.031 |
| MAGI2 | 11 | 0 | 0.001 | Inf | Inf | 2.82 | 0.031 |
| SLC39A12 | 11 | 0 | 0.001 | Inf | Inf | 2.82 | 0.031 |
| MUC16 | 30 | 11 | 0.001 | 4.083 | 10.214 | 1.734 | 0.031 |
| TCHH | 13 | 1 | 0.001 | 15.644 | 682.961 | 2.218 | 0.049 |

**Figure 2.**
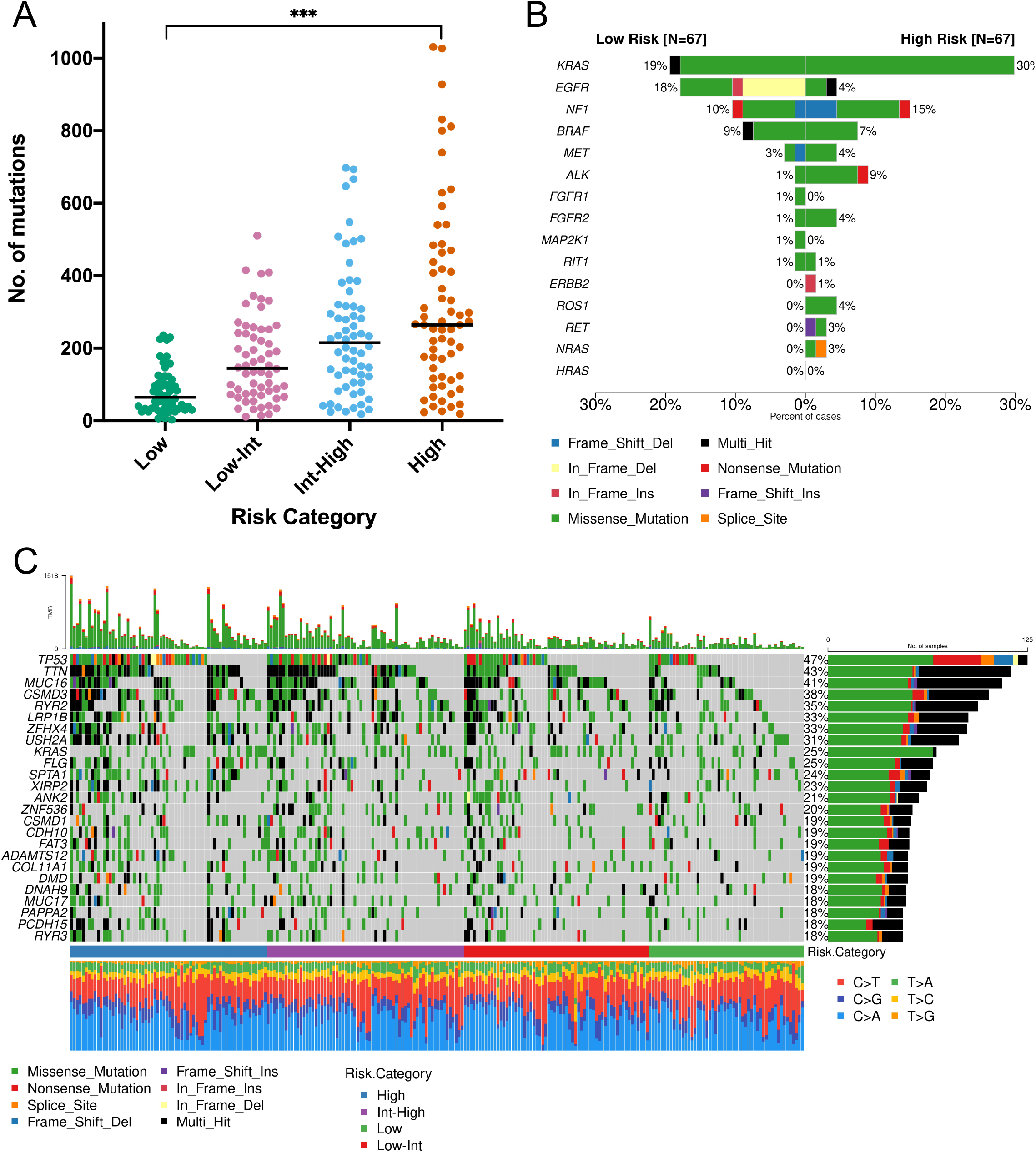
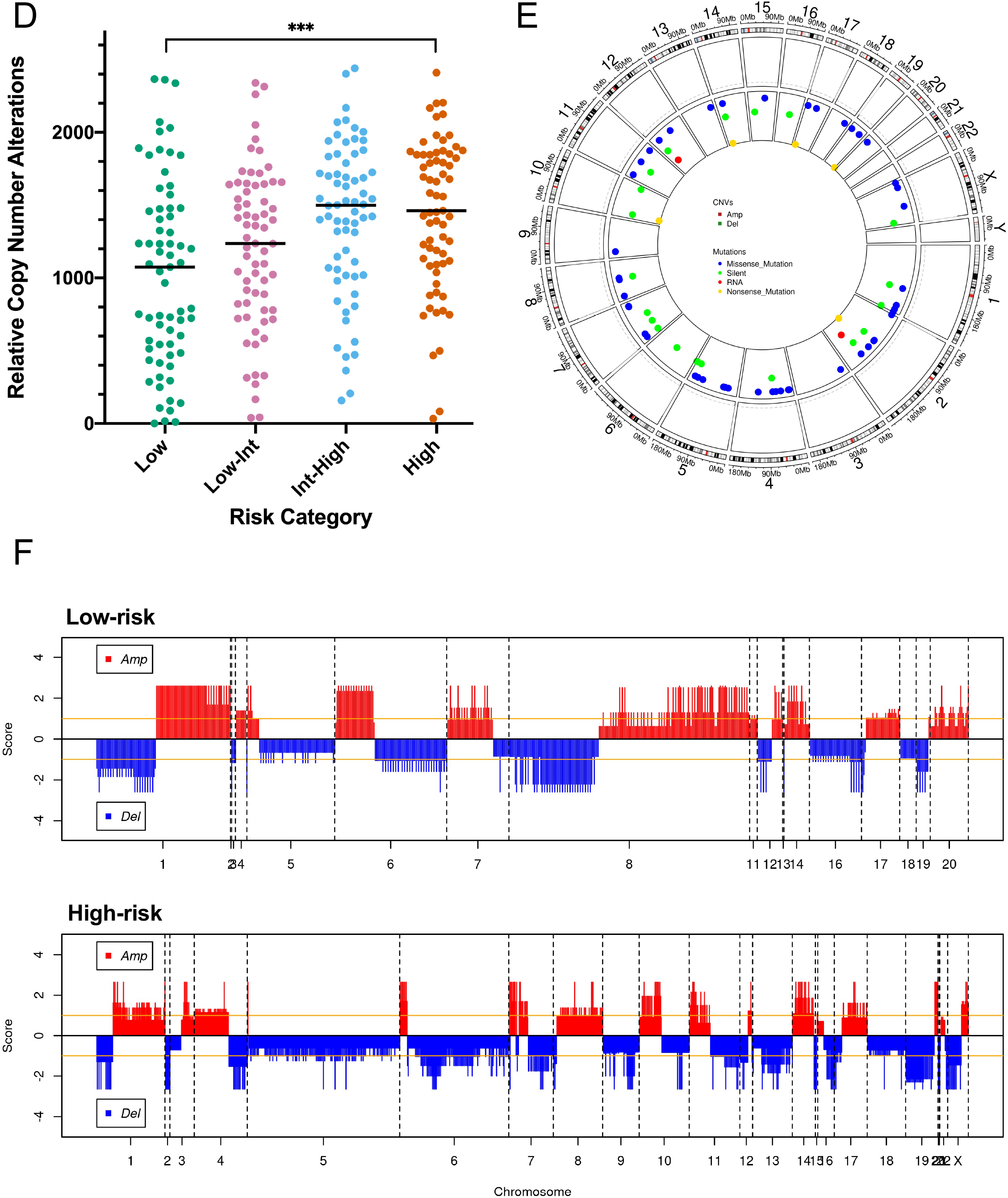
Genetic alterations in recurrent lung adenocarcinomas. Increasing risk of recurrence is associated with increasing tumor mutation burden (A). Canonical driver mutations are present in stage I tumors but do not occur more frequently in recurrent tumors (B). An oncoplot of the most frequently mutated genes in the stage I lung adenocarcinoma cohort is shown in (C). Copy number alterations occur more frequently in tumors with increasing risk of recurrence (D). Although copy number alterations and mutations are evenly distributed through the genome in recurrence high-risk tumors (E), high-vs. low-risk tumors show different genome-wide copy number alteration patterns (F).

### Transcriptome analysis

Comparing transcriptomes of recurrence high vs. low-risk tumors, 10,398 genes were differentially expressed (adjusted *P* value < 0.05). Filtering these genes by an absolute log fold-change > 0.6 and adjusted *P* value < 0.001, we identified a total of 4,150 top differentially regulated genes (2,544 downregulated and 1,606 upregulated genes, Supplemental Figure 2A). Heatmap clustering of these top differentially regulated genes resulted in clear separation between high- and low-risk tumors across four different gene clusters (Figure 3A). Stage also showed some separation by heatmap clustering. In contrast, other clinical co-variates including age, number of pack-years smoked, and sex were not significantly correlated with these gene clusters (Figure 3A).

**Figure 3.**
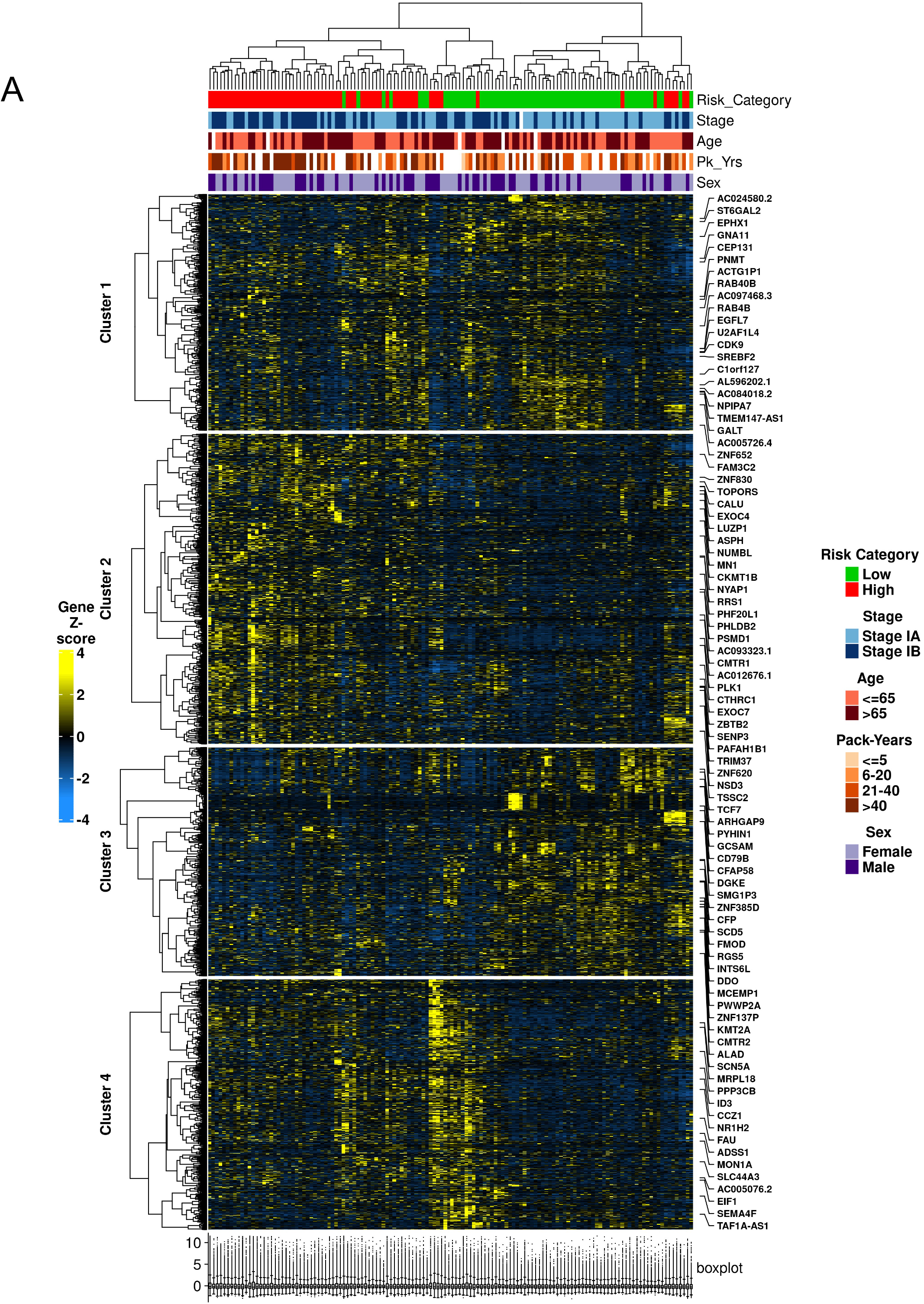
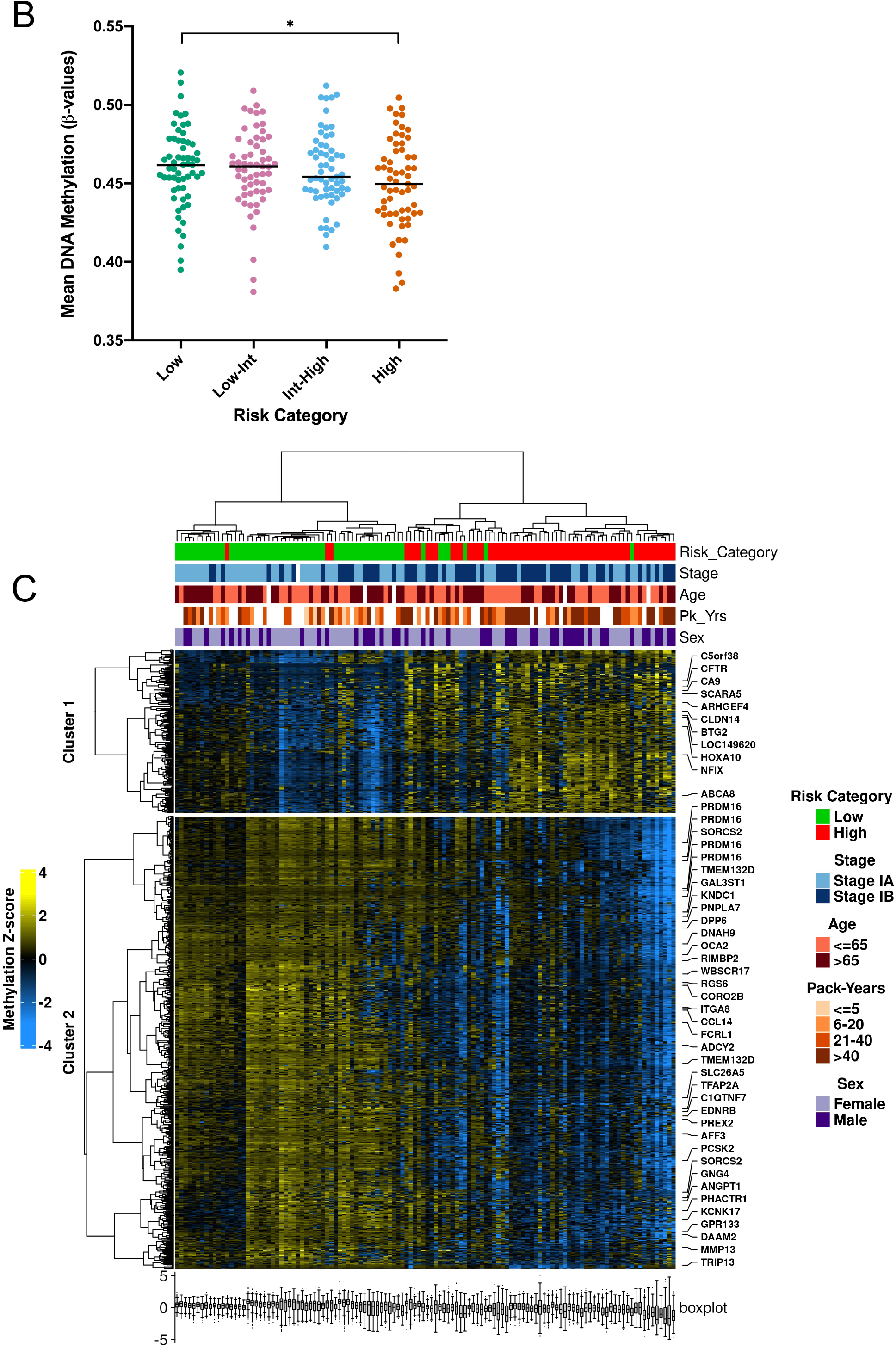
Transcriptomic analysis demonstrates distinct genomic and immunologic features of recurrent lung adenocarcinomas. Heatmap clustering of differentially regulated genes results in clear separation between high- and low-risk tumors across four different gene clusters (A). Differential methylation patterns are also associated with recurrent tumors; decreasing DNA methylation is observed in tumors with increasing risk of recurrence (B) while heatmap clustering of differentially methylated genes results in clear separation between high- and low-risk tumors across two different gene clusters (C).

Recurrence high-risk tumors were associated with overexpression of genes in cluster 2 (Figure 3A). Pathway analysis of genes in this cluster revealed activation of numerous cancer-related pathways including kinetochore metaphase signaling, cell cycle control of chromosomal replication, role of BRCA1 in DNA damage response, mitotic roles of polo-like kinase, hereditary breast cancer signaling, role of CHK proteins in cell cycle checkpoint control, and estrogen-mediated S-phase entry pathways (Supplemental Table S2).

Simultaneously, recurrence high-risk tumors were associated with downregulation of genes in cluster 3 (Figure 3A). Genes in cluster 3 were significantly associated with immune response pathways including antigen presentation, TH1 and Th2 activation, B cell development, neuroinflammation signaling, PD-1 and PD-L1 cancer immunotherapy, and IL-4 signaling pathways (Supplemental Table S3). 16 out of 39 (41%) genes involved in antigen presentation were significantly downregulated in high-risk tumors (*P* < 0.001) including *CIITA*, a master control regulator of class II MHC transcription (2.3 fold), numerous MHC class I and II genes including *HLA-E* (1.6 fold), *HLA-DMA* (2.7 fold), *HLA-DMB* (2.0 fold), *HLA-DOA* (3.0 fold), *HLA-DOB* (2.7 fold), *HLA-DPA1* (2.5 fold), *HLA-DPB1* (2.6 fold), *HLA-DQA1* (2.8 fold), *HLA-DQB1* (3.2 fold), *HLA-DQB2* (4.3 fold), *HLA-DRA* (2.2 fold), *HLA-DRB1* (3.0 fold), *HLA-DRB5* (4.0 fold), and *CD74*, a class II MHC chaperone molecule (2.5 fold). Simultaneously, 32 out of 172 (18.6%) genes involved in Th1 and Th2 activation were significantly downregulated in high-risk tumors including the class II MHC genes above, *CCR4* (2.7 fold), *CD40LG* (3.7 fold), *IL24* (1.8 fold), *IL33* (3.1 fold), *IL12B* (2.8 fold), *IL6R* (2.4 fold), *TGF-β* receptors 2 and 3 (1.5 and 2.3 fold respectively), *PTDGR2* (3.4 fold), *NFATC1, NFATC2*, and *NFATC3* (1.5, 1.6, and 1.4 fold respectively), *PIK3R1* and *PIK3R6* (1.4 and 1.8 fold respectively), and *PRKCQ* (2.4 fold).

Topology-based enrichment analysis is a third-generation pathway analysis technique that better relates gene expression patterns to biological pathways by taking the role, hierarchy, and expression direction of pathway genes into account^58^. Using topology-based enrichment analysis, 42 pathways are significantly perturbed in high-vs. low-risk tumors (pPERT < 0.05 and pGFdr < 0.05, Supplemental Table S4). Numerous cancer-related pathways are activated in recurrence high-risk tumors including central carbon metabolism in cancer, toll-like receptor signaling, and hedgehog signaling pathways. Simultaneously, cellular senescence, Ras signaling, and PPAR signaling pathways are inhibited in high-risk tumors. Multiple immune-related pathways are also perturbed in high-risk tumors. Human cytomegalovirus infection, bacterial invasion of epithelial cells, and complement and coagulation cascades pathways are activated. In contrast, cytokine-cytokine receptor interaction, human T-cell leukemia virus 1 infection, systemic lupus erythematosus, and intestinal immune network for IgA production pathways are inhibited in high-risk tumors (Supplemental Table S4).

### Epigenetic analysis

To investigate epigenetic features associated with recurrence, methylation differences between high- and low-risk tumors were analyzed. Increasing recurrence risk was associated with decreasing DNA methylation across risk categories (median β-value of 0.462, 0.461, 0.454, and 0.450 for low-, low-intermediate-, intermediate-high-, and high-risk categories respectively [*P* = 0.042 between low- and high-risk categories], Figure 3B). A total of 1,514 genes were differentially methylated between high- and low-risk tumors (absolute log fold-change > 0.4 and adjusted *P* value < 0.05). Genes were more than three times as likely to be hypo-rather than hypermethylated in the high-risk cohort (1,173 hypomethylated genes vs. 341 hypermethylated genes, Supplemental Figure 2B). The top 10 differentially methylated genes were *FOLR1, LEAP2, CRTAM, MIR1287, KIAA0513, GZMB, HTRA1, TMEM11, FAM49A*, and *SORCS3* (Supplemental Figure 2C). Heatmap clustering of differentially methylated genes resulted in clear separation between high- and low-risk tumors across two different gene clusters (Figure 3C). Stage also showed some separation by heatmap clustering. In contrast, other clinical co-variates including age, number of pack-years smoked, and sex were not significantly correlated with these gene clusters (Figure 3C). Pathway analysis of differentially methylated genes identified significant involvement of immune-related pathways including antigen binding, processing, and presentation pathways by Gene Ontology (GO) analysis and allograft rejection by Molecular Signatures Database analysis (Supplemental Table S5).

Differential methylation analysis was also performed on a chromosome level, revealing 3,461 differentially methylated regions (adjusted *P* < 0.001). As differential methylation pathway analysis identified antigen presentation pathways as significantly involved in high-risk tumors, chromosome 6 was examined for differentially methylated regions. High-risk tumors demonstrated heavy methylation in a specific region of chromosome 6 (bp 32,410,873 to 33,041,697) containing class II MHC genes (*HLA-DMB, HLA-DOA, HLA-DPA1, HLA-DQA2, HLA-DQB2, HLA-DRA*, Supplemental Figure 2D).

Integrated transcriptome and methylation analysis demonstrated a total of 295 genes that were both hypomethylated and upregulated in high-risk patients including *GPR115, KRT6A, KRT6B, KRT16, PPAPDC1A, PADI1, HMGA2, SLC2A1, CCL7* and *CCL11* (Supplemental Figure 2E). 771 genes were both hypermethylated and downregulated in high-risk patients including *FOLR1, NR0B2, AGER, GPR116, CLDN18, CYP4B1, PGC, KIAA0408*, and *PIGR* (Supplemental Figure 2E). Integrated analysis using enhancer linking also identified multiple differentially methylated transcription factor promoter binding sites. The transcription factors CENPA, MYBL2, FOXM1, ARNTL2, SHOX2, GBX2, E2F2, E2F7, HMGA1, and SP6 were found to be upregulated and bind hypomethylated promoter sites (Supplemental Figure 3A), whereas IRX5, NKX2-1, HLF, ZBTB18, ATOH8, PRDM16, IRX3, IRX2, ZNF491, and NR3C2 were found to be downregulated and bind hypermethylated promoter sites in high-vs. low-risk tumors (Supplemental Figure 3B).

### Tumor Immune Microenvironment

In order to obtain lung adenocarcinoma specific tumor infiltrating leukocyte profiles, we performed single-cell RNA-seq analysis on freshly resected tumor and adjacent normal lung tissues from six patients who underwent complete surgical resection of stage I lung adenocarcinoma at UCSF. Twenty distinct immune cell populations were identified by scRNA-seq analysis including 10 adaptive immune populations (helper T-cells, cytotoxic T-cells, exhausted cytotoxic T-cells, δγ T-cells, naïve T-cells, regulatory T-cells, B cells, plasma cells, NKT cells, and NK cells) and 10 innate immune populations (M1 macrophages, M2 macrophages, classical monocytes, intermediate monocytes, non-classical monocytes, mast cells, conventional dendritic cells 1, conventional dendritic cells 2, plasmacytoid dendritic cells, and neutrophils) (Figure 4A). Unique markers from these immune populations were used to calculate immune cell population densities from bulk RNA-seq data in the TCGA stage I lung adenocarcinoma population using digital cytometry. Despite a positive association between neoantigen burden and recurrence risk (73, 141, 173, and 234 predicted neoantigens in low-, low-intermediate, intermediate-high, and high-risk cohorts respectively [*P* < 0.001 between low- and high-risk cohorts], Figure 4B), a negative correlation between total immune cell infiltration and recurrence was observed (median relative immune cell densities of 54.1, 49.3, 49.0, and 46.1 in low-, low-intermediate, intermediate-high, and high-risk cohorts respectively [*P* < 0.001 between low- and high-risk cohorts], Figure 4C). Heatmap clustering of immune populations resulted in clear separation between recurrence high- and low-risk tumors across two distinct immune cell clusters (Figure 4D). Stage also showed some separation by heatmap clustering. In contrast, other clinical covariates including age, number of pack-years smoked, and sex were not significantly correlated with these immune cell clusters (Figure 4D). The cluster associated with recurrence high-risk tumors was depleted in adaptive immune populations including cytotoxic T-cells, NKT cells, helper T-cells, B-cells, δγ-T-cells, plasma cells, and T regulatory cells (Figure 4D). UMAP analysis also demonstrated clear tumor immune profile differences between recurrence high- and low-risk stage I lung adenocarcinoma tumors. When clustered by immune population densities only, separation of high- and low-risk tumors into distinct groups was observed (Figure 4E). Relative to low-risk tumors, high-risk tumors were significantly enriched in classical and non-classical monocytes, neutrophils, naïve T-cells, and exhausted cytotoxic T-cells (Figure 4F). In contrast, they were significantly depleted in adaptive immune populations including regulatory T-cells, helper T-cells, B-cells, cDC1s and cDC2s, and intermediate monocytes (Figure 4F).

**Figure 4.**
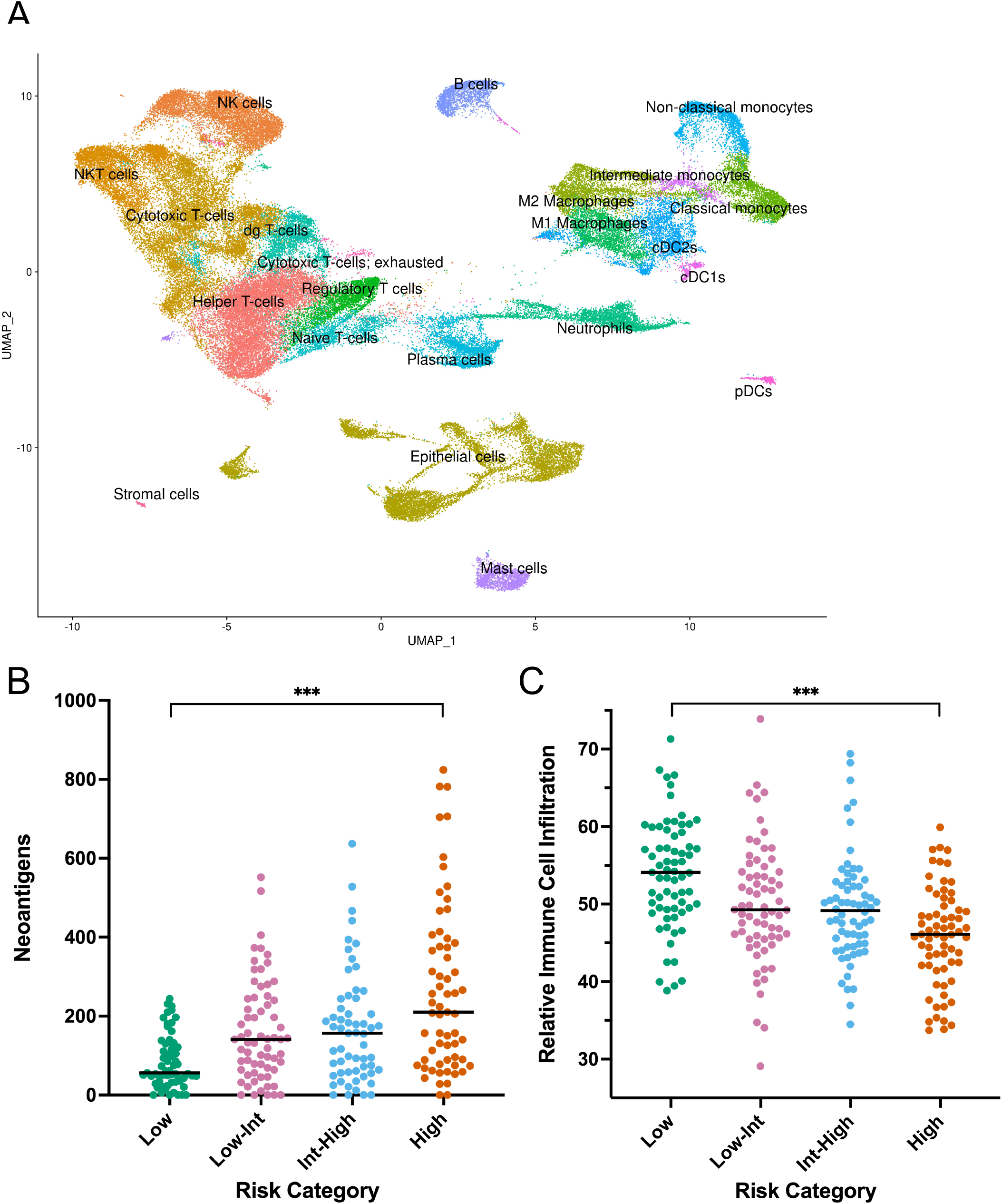
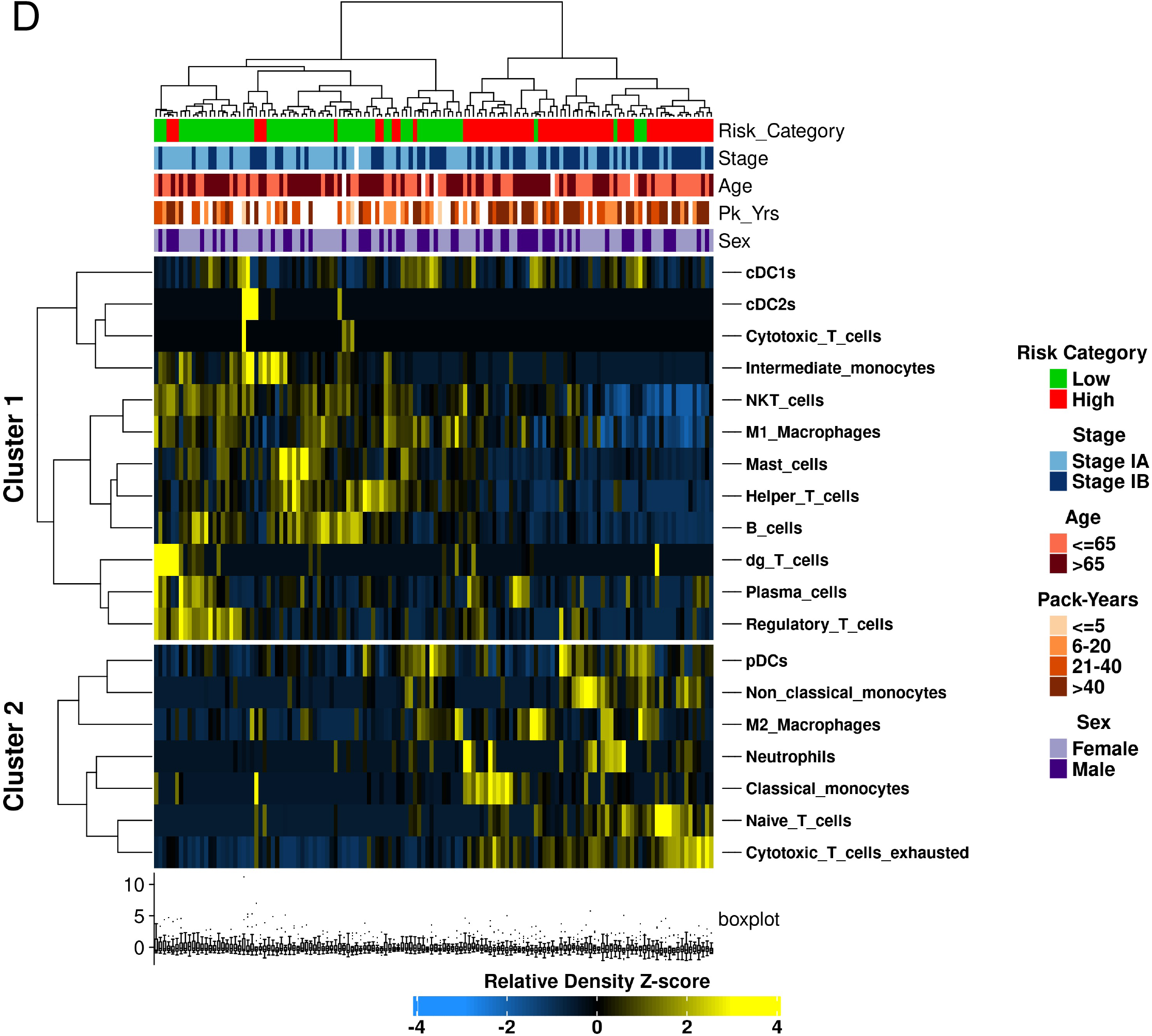
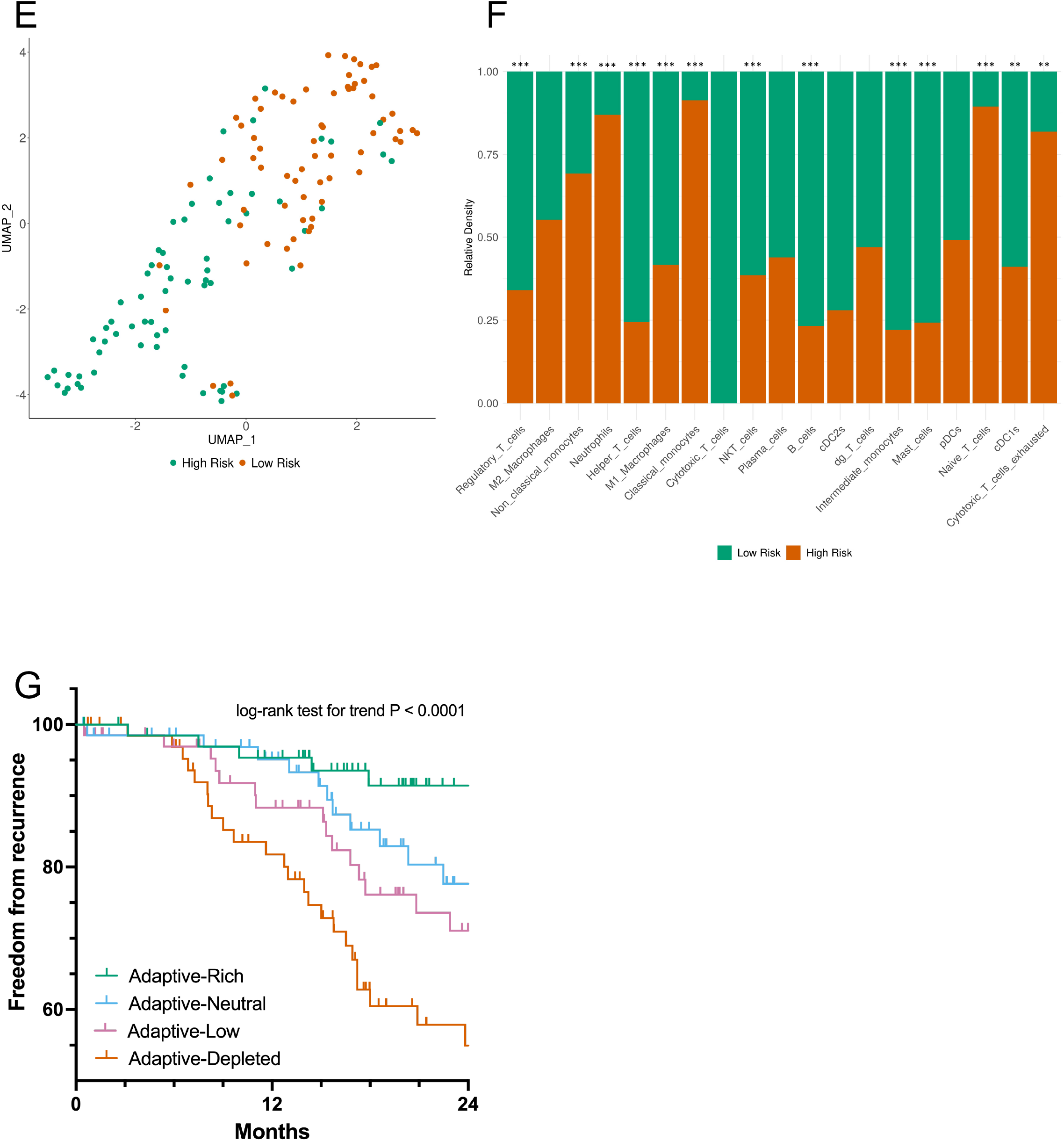
Recurrent stage I lung adenocarcinomas display adaptive immune cell depleted phenotypes. scRNAseq analysis reveals twenty distinct immune cell populations in stage I lung adenocarcinomas (A). Despite increasing antigenicity (B), tumors with increasing risk of recurrence demonstrate decreasing immune cell infiltration (C). Heatmap clustering (D) and UMAP analysis (E) demonstrate distinct immune profiles in tumors with low-vs. high-risk of recurrence; these differences are most prevalent in adaptive immune cell populations (F). Patients with adaptive-rich immune cell phenotypes have improved 24-month freedom from recurrence compared to tumors with adaptive-depleted phenotypes (G).

Adaptive immune population density was also prognostic of recurrence in our stage I lung adenocarcinoma cohort. Using cox regression analysis, an immune profile classifier was identified that clustered patients based on the relative abundance of adaptive tumor-infiltrating immune populations. Patients with adaptive-rich, adaptive-neutral, adaptive-low, and adaptive-depleted phenotypes had 24-month freedom from recurrence of 91.4%, 77.7%, 71.0%, and 54.9% respectively ([*P* < 0.001], Figure 4G).

## Discussion

Although recurrence in early-stage adenocarcinoma is common, the genetic and immunologic features of recurrence in lung cancer are not well described. One approach to describing these features is to treat recurrence as a binary event by grouping patients into cohorts of “recurred” vs “did not recur”. Recurrence, however, is a time-to-event datapoint. Early recurrence is not the same biological event as late recurrence. In addition, patients who have not recurred may only be labeled as such as they have not been followed long enough to experience recurrence. Lastly, recurrence is notoriously difficult to track accurately in the real world as many patients are lost to follow-up after surgery and other treatments for early-stage disease^59^. Using recurrence as a binary endpoint to group patients, therefore, is a suboptimal method to understand the biological features of recurrence. To address these concerns and incorporate time-to-recurrence data as an endpoint in this study, a recurrence score was developed which relates the patient’s probability of developing recurrence to the biologic features of the tumor. This strategy not only takes time-to-event data into account, but also lessens the impact of inaccurate clinical follow-up by simultaneously clustering the genomic profiles of these patients with their clinical endpoints. In the clinical setting, clustering patients by biological features related to time-to-event outcomes is a superior strategy that has been proven to add prognostic and predictive value in multiple studies^2, 23–25, 27–29, 60–62^. The recurrence score developed here is predictive of recurrence in the TCGA lung adenocarcinoma cohort and allows for further exploration of the biological features of early-stage recurrence.

Tumor mutation burden (TMB) has been shown to be predictive of immunotherapy response in prior studies but has not been reported to be associated with recurrence^63^. In this study, early-stage lung adenocarcinomas with higher TMB had significantly higher rates of recurrence. Copy number alterations also occurred more frequently in recurrent tumors, signifying genomic instability as a hallmark of recurrence. *TP53* was the most frequently mutated gene in the stage I cohort and was more frequently mutated in recurrent tumors in our study. *MUC16* (*CA-125*) was also observed to be frequently mutated in recurrent tumors. MUC16 is overexpressed in several cancers including ovarian, pancreatic, breast, and lung, and has been shown to be associated with cancer progression and poor prognosis^64^. *SMARCA4*, also frequently mutated in recurrent tumors in our study, has been previously shown to be associated with recurrence in lung adenocarcinoma^32, 33^. SMARCA4 is a DNA-dependent ATPase which is involved in transcription regulation via the SWI/SNF chromatin remodeling complex^65^. Mutations in *SMARCA4* can co-occur with *KRAS, STK11*, and *KEAP1* mutations^65^. *SMARCA4* mutations are prognostic of poor overall survival, especially in patients with *KRAS*-mutant tumors^65^. *MUC16* and *SMARCA4* are attractive therapeutic targets in patients at high-risk for recurrence after surgery and may warrant further investigation.

Other frequently mutated genes in our study included very large genes such as *TTN* (304,814 bases), *CSMD3* (1,214,572 bases), *RYR2* (791,805 bases), and *LRP1B* (1,901,041 bases). The presence of mutations in these large genes suggests that many mutations in recurrent tumors are random rather than targeted or selected events. With respect to canonical lung cancer driver mutations, previous studies have failed to demonstrate strong correlations between canonical lung cancer driver mutations and recurrence^32, 33^. This was also observed in our study where no canonical driver mutations were found to occur more frequently in recurrent tumors, suggesting that the biological processes of tumor initiation and propagation are distinct.

Transcriptomic analysis revealed discrete genomic and immunologic features associated with early-stage lung adenocarcinoma recurrence. As expected, a number of canonical cancer-related pathways involving cell cycle control, metaphase signaling, DNA damage response, mitosis, checkpoint control, and mismatch repair are strongly activated in recurrent tumors. These cancer pathways have previously been described as hallmarks of cancer progression and metastasis^66–69^. Hypomethylation has also been show in previous work to be associated with genome instability and cancer progression^70^. We also observed decreasing methylation in recurrent tumor phenotypes, which may partly explain the mutational instability and active cell cycle progression observed in the pathways above. Overall, the activation of these pathways and decreasing levels of methylation support the notion of an active, unstable genome in recurrent stage I lung adenocarcinomas.

In contrast, a number of key immune response pathways are strongly downregulated in recurrent tumors. This suggests that the presence of a permissive tumor microenvironment is also an important cause of early-stage lung adenocarcinoma recurrence. In support of this, recurrent tumors display less overall immune cell infiltration despite having higher mutation and neoantigen burden than non-recurrent tumors, consistent with a “cold” tumor landscape. In addition, the top genes present in the downregulated transcriptome cluster of recurrent tumors play vital roles in antigen presentation and Th1 and Th2 activation. Antigen presentation and Th1 and Th2 activation pathways are intimately associated with each other as antigen presentation by MHC molecules stimulates Th1 and Th2 differentiation and activation^71^. Notably, epigenetic analysis of recurrent tumors demonstrates heavy methylation of antigen presentation pathway genes including a specific region of chromosome 6 containing numerous class II MHC genes, suggesting that hypermethylation is one mechanism of antigen presentation suppression.

Other groups have used two main strategies to profile the tumor immune microenvironment in early-stage lung adenocarcinoma: 1. immunohistochemistry that allows for direct measurements of a limited number of specific immune cell populations and 2. computational techniques that allow for more comprehensive but indirect measurements of multiple immune cell populations. In the largest study of immune populations in stage I lung adenocarcinoma patients to date, Suzuki et al. investigated eight types of tumor-infiltrating leukocytes and five cytokines using immunohistochemistry^35^. This group observed that stromal FoxP3/CD3 ratio, tumor IL-12Rβ2, and tumor IL-7R were associated with recurrence, whereas the other immune cell populations including helper T cells, cytotoxic T cells, B cells, memory T cells, NK cells, and macrophages had no significant association^35^. Yan et al., in contrast, did not find a significant association between FoxP3-positive T regulatory cells and disease-free survival in their characterization of stage I-IV non-small cell lung cancer patients using immunohistochemistry^36^. Although our data also do not support this positive association between T regulatory cells and recurrence, neither our group nor Yan et al. distinguished between tumor nest or stromal T regulatory cell populations which may explain the lack of concordance. Varn et al. used computational techniques to characterize CD4+ T-cell, CD8+ T-cell, naïve B cell, memory B cell, NK cell, and myeloid cell populations in lung adenocarcinoma^38^. Murine immune cell gene expression profiles were used to develop consensus immune signatures for these six different immune populations. Consistent with our observations, they observed that low naïve B cell and CD8+ T-cell populations and high monocyte populations were associated with recurrence in stage I lung adenocarcinomas^38^. Other groups have used blood-based immune profiles to comprehensively characterize infiltrating immune populations in lung adenocarcinoma^37, 72^, however, the association between these immune populations and recurrence was not reported. To our knowledge, our study is the first study to combine direct measurements of tumor infiltrating immune cell populations using scRNA-seq with computational profiling in order to comprehensively characterize immune cell populations in a large cohort of patients with recurrent stage I adenocarcinoma.

Using this approach, we observed that the lack of sufficient antigen presentation and T-helper cell response in recurrent stage I tumors found in our transcriptomic analysis is supported by our analysis of specific infiltrating immune populations in recurrent vs. non-recurrent tumors. Although dendritic cell and macrophage populations are similar between recurrent and non-recurrent tumors, non-recurrent tumors are heavily infiltrated by B-cells while recurrent tumors are relatively lacking in these antigen-presenting cells. In addition, recurrent tumors exhibit relatively sparse populations of helper T-cells while displaying a high proportion of exhausted cytotoxic T-cells. These data suggest that recurrent tumors may evade immunosurveillance by active downregulation of antigen presentation to T-helper cells, resulting in an insufficient adaptive immune response. In support of this, we observed that adaptive-depleted tumors had significantly worse prognosis that adaptive-rich tumors, recurring five-times more frequently within two years of diagnosis.

There are several limitations to this study. First, recurrence is a time-to-event datapoint and the use of the TCGA lung adenocarcinoma cohort does not allow for independent verification of patient follow-up data such as recurrence and time-to-recurrence. Although the creation of a recurrence score helps mitigate inaccurate outcomes reporting and inadequate patient follow-up in this setting, the accuracy of the recurrence score used in this study could be improved with independent validation of clinical outcomes data. Second, although the TCGA cohort is the largest-ever assembled lung adenocarcinoma patient cohort with full genetic, genomic and epigenetic analysis, only 268 stage I patients were fully analyzed in this study. It is likely that stronger biological conclusions could be reached with a larger stage I lung adenocarcinoma patient cohort. Lastly, the use of computational techniques to quantify immune populations present in this large stage I lung adenocarcinoma cohort was necessary as direct measurements of immune cell densities were not available. Single-cell RNA-seq and spatial transcriptomics data will one day be available in large cohorts such as the one analyzed here and may allow for direct measurements of immune cell infiltration, integrated immune cell transcriptome analysis, and differentiation between tumor and stromal immune cell populations.

In conclusion, recurrence in early-stage lung adenocarcinoma is common despite complete surgical resection as the mainstay of therapy. Recurrent stage I lung adenocarcinomas display features of genomic and genetic instability including increased tumor mutation burden, neoantigen load, activation of numerous mitotic and cell cycle genes, and decreased genome-wide methylation burden. In parallel, impaired antigen presentation and Th1/Th2 responses appear to be central features of permissive tumor immune microenvironments in recurrent tumors and may be responsible for worse prognosis associated with adaptive-depleted tumors.

## Acknowledgements

The results described here are in whole or part based upon data generated by the TCGA Research Network: https://www.cancer.gov/tcga.

