## Supplemental Figure 1 for "Multiomic Characterization of Stage I Lung Adenocarcinoma Reveals Distinct Genetic and Immunologic Features of Recurrent Disease"

A

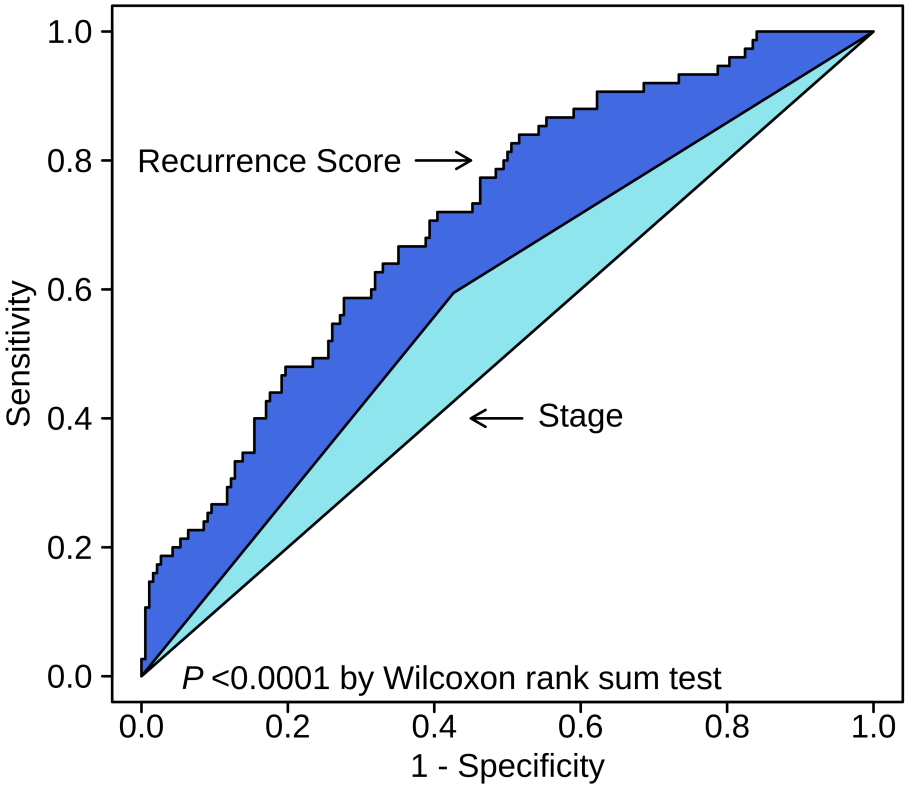


**Supplemental Figure 1. Recurrence Score improves recurrence predictions vs. stage alone.** The recurrence score increases the time-dependent AUROC to 0.714 from 0.584 vs. stage alone ($P < 0.001$).
