## Supplemental Figure 2 for "Multiomic Characterization of Stage I Lung Adenocarcinoma Reveals Distinct Genetic and Immunologic Features of Recurrent Disease"

A

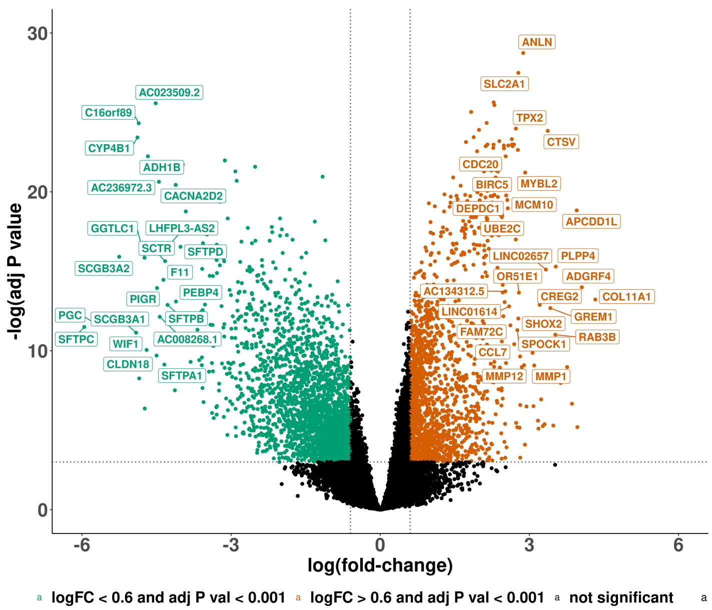


B

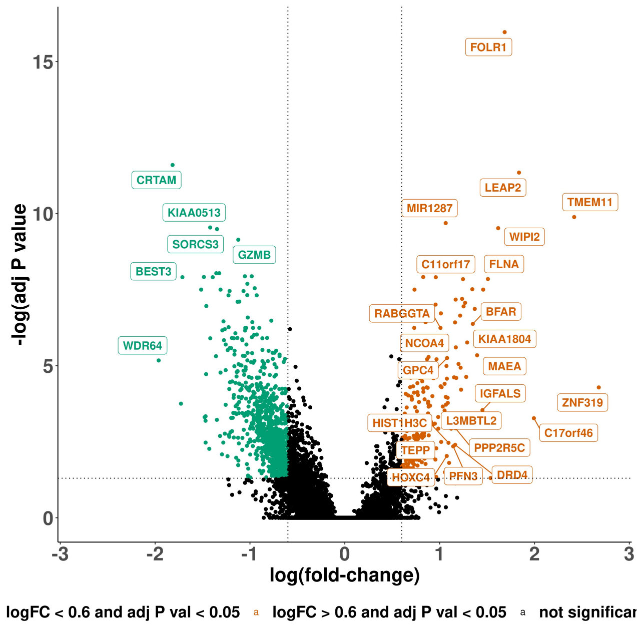


C

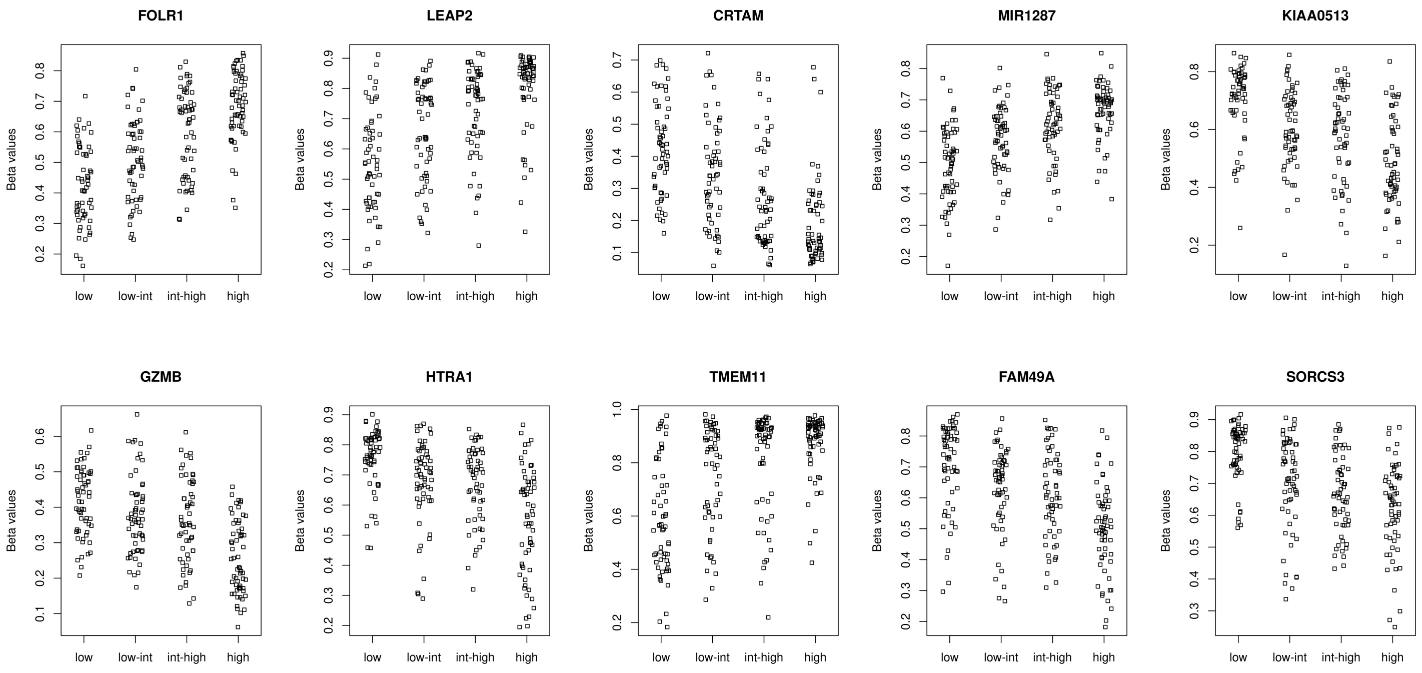

### Supplemental Figure 2

D

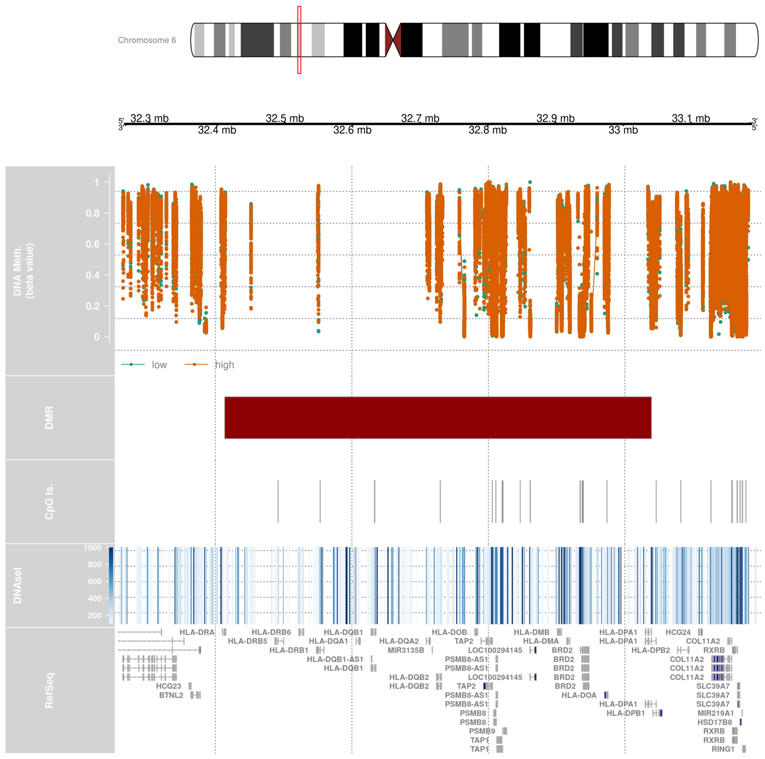

E

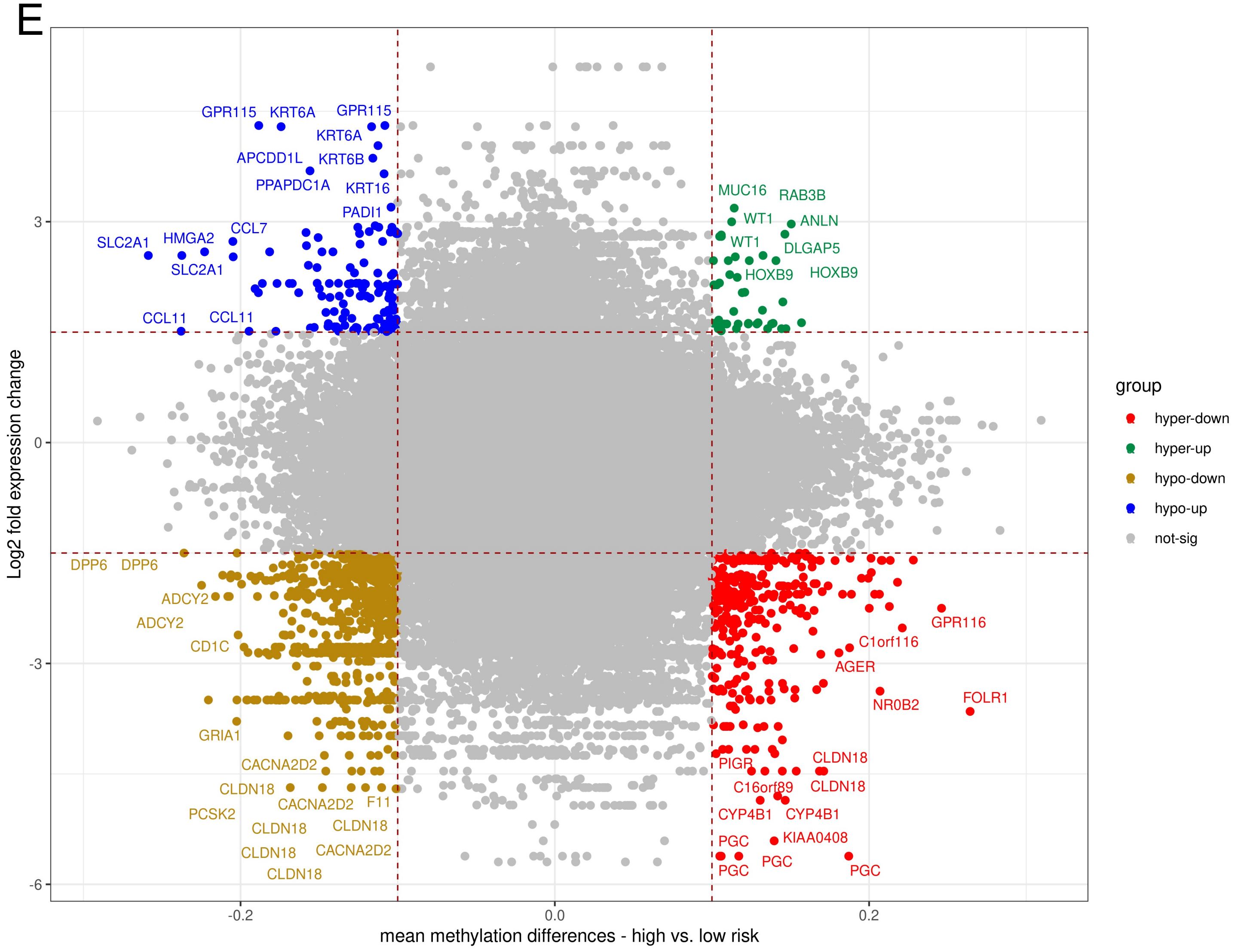

**Supplemental Figure 2. Genomic and epigenetic alterations in recurrent high- vs. low-risk stage I lung adenocarcinomas.** Volcano plots of differential gene expression (A) and differential methylation (B) in recurrent high- vs. low-risk tumors. The top 10 differentially methylated genes by recurrent risk category are shown in (C). Analysis of differentially methylated regions (DMRs) demonstrates heavy methylation of the chromosome 6 region containing class II MHC genes (D). Integrated transcriptome and methylation analysis demonstrates a total of 295 genes that are both hypomethylated and upregulated (blue quadrant) and 771 genes that are both hypermethylated and downregulated (red quadrant) in high- vs. low-risk patients (E).
