## Supplemental Figure 3 for "Multiomic Characterization of Stage I Lung Adenocarcinoma Reveals Distinct Genetic and Immunologic Features of Recurrent Disease"

## A

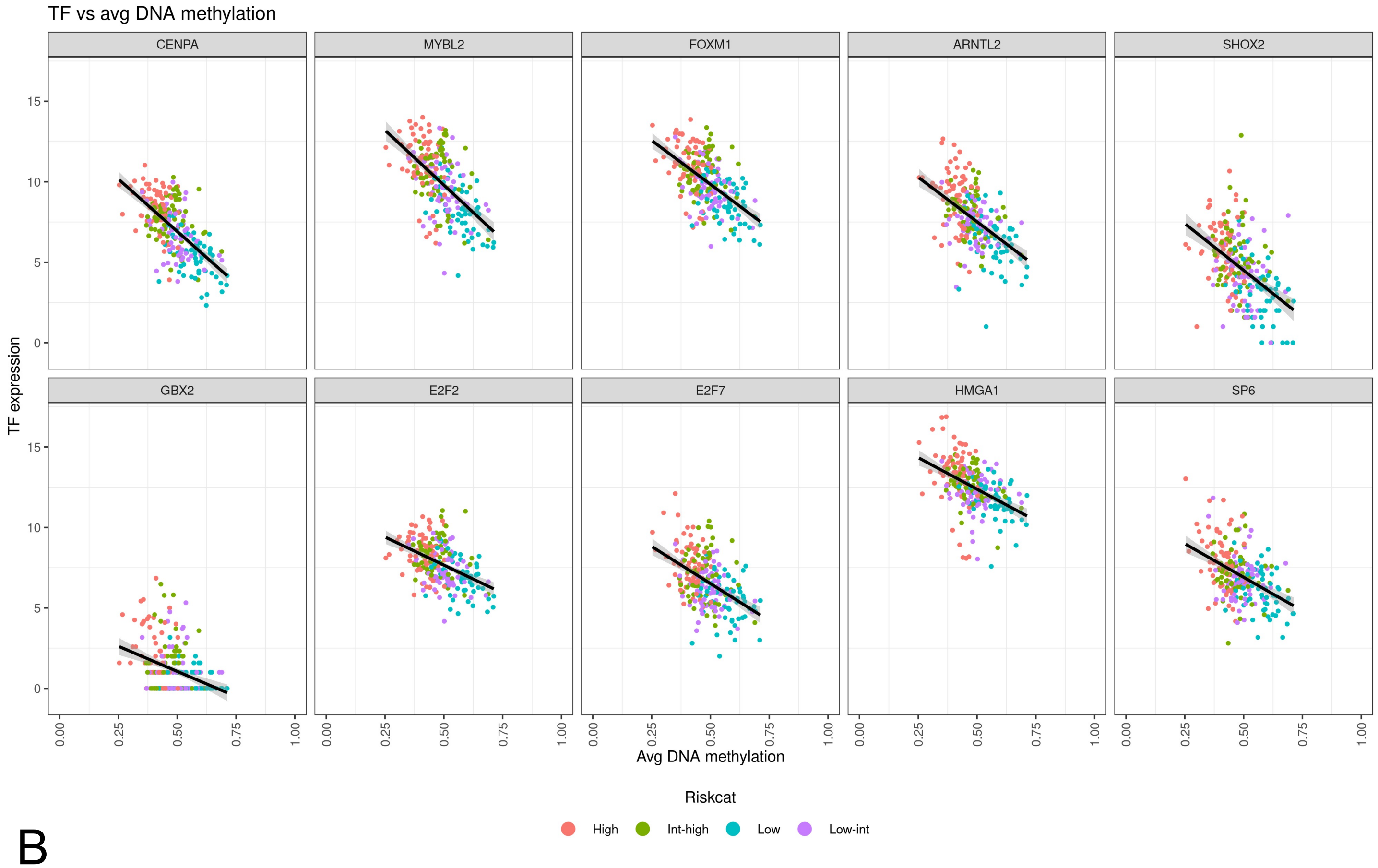


## B

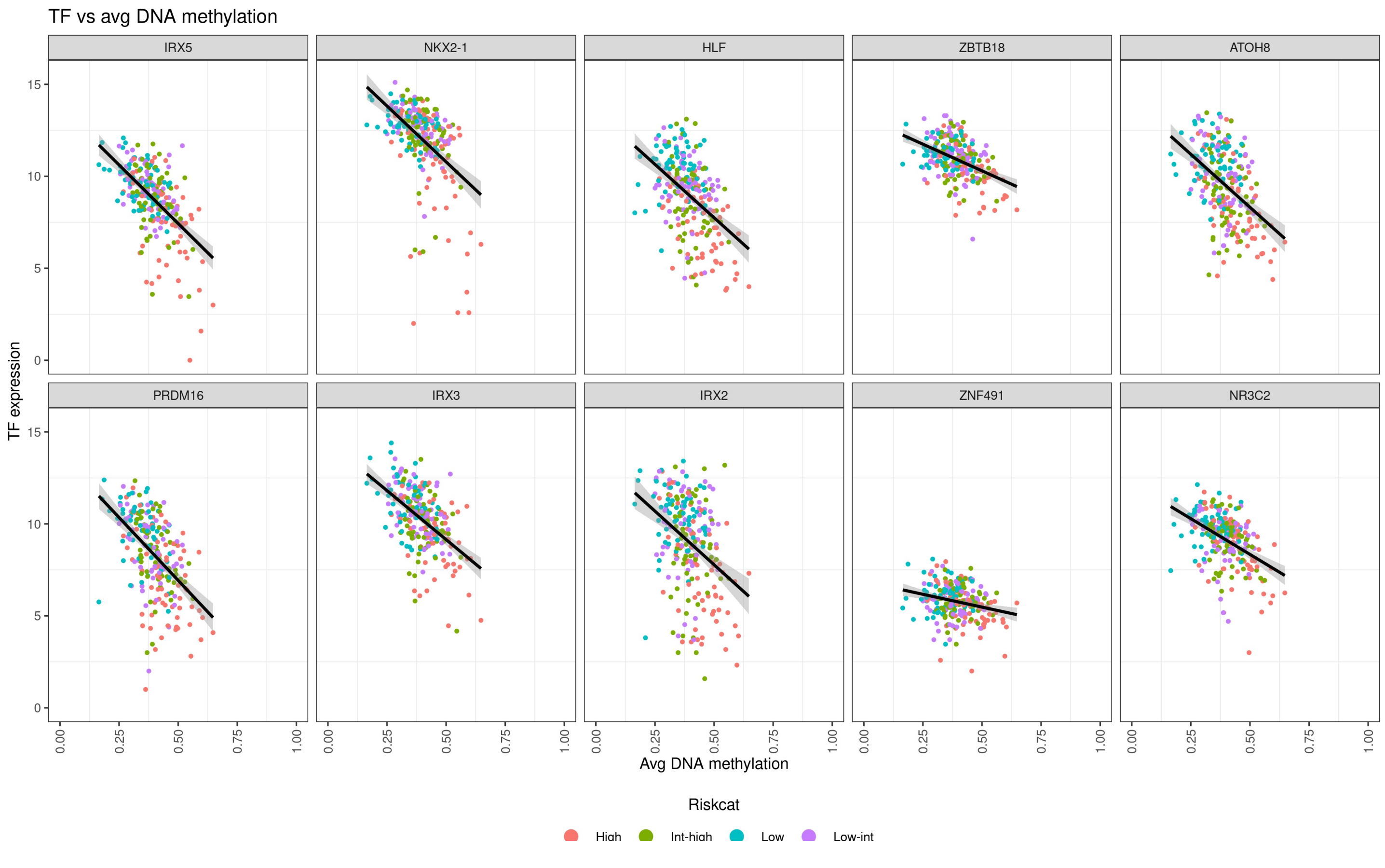

**Supplemental Figure 3. Integrated analysis using enhancer linking identifies multiple differentially methylated transcription factor promoter binding sites.** The top 10 upregulated transcription factors with hypomethylated promoter binding sites (A) and downregulated transcription factors with hypermethylated promoter binding sites (B) are shown; recurrence risk categories are labeled by color.
