## Supplemental Table S1 for "Multiomic Characterization of Stage I Lung Adenocarcinoma Reveals Distinct Genetic and Immunologic Features of Recurrent Disease"

**Table S1. Genes and Coefficients in Recurrence Risk Score Model**

| Gene | Coefficient |
|---|---|
| LDHA | 0.011638925 |
| PAX5 | -0.00068286 |
| AC046143.1 | 0.004911234 |
| TANGO2 | -0.00607408 |
| ZNF341-AS1 | 0.009427997 |
| CASP12 | -0.005758507 |
| ZNF763 | -0.002287844 |
| H3C13 | 0.004328153 |
| FAM117A | -0.034132088 |
| XRCC5 | 0.001544703 |
| CIDEC | 0.001305843 |
| INPP5J | -0.000471615 |
| ANLN | 0.005611476 |
| AL596223.1 | 0.002101452 |
| AC079601.2 | -0.001371535 |
| BEST3 | 0.000201193 |
| CCR6 | -0.002009526 |
| SKIL | 0.010454531 |
| KLRG2 | -0.000898403 |
| MPP6 | 0.003017565 |
| AC004947.2 | -0.004915431 |
| AP000695.2 | 0.002646879 |
| CREG2 | 0.002145222 |
| SEC14L4 | -0.001118089 |
| LDLRAD3 | 0.020860561 |
| KIR2DL3 | -0.002720483 |
| YWHABP2 | 0.022408439 |
| AKAP12 | 0.002065057 |
| LINC01117 | 0.009931264 |
| BTBD7P1 | -0.001467806 |
| RHOV | 0.001595022 |
| OPN3 | 0.008829789 |
| DKK1 | 0.004503076 |
| CHRNA6 | -0.012090302 |
| FAM76A | -0.01635178 |
| CYP4B1 | -5.48E-05 |
| CRHR2 | -0.003313881 |
| TLE1 | 0.022100011 |
| PLEKHB1 | -0.007319084 |
| B4GALT1 | 0.008964576 |
| TEX15 | 0.002233072 |
| GNG7 | -0.007211389 |
| KIAA0408 | -0.009124867 |
| SYNPR-AS1 | -0.011220243 |
| LINC00667 | -0.004310673 |
| AC068228.2 | 0.009142372 |
| AC127070.2 | -0.003982105 |
| AP000695.1 | 0.013513884 |
| TMEM213 | -0.005681923 |
| SLC47A1 | -0.001912309 |
| VIM-AS1 | -0.004906454 |
| EPHX1 | -0.003544239 |
| AC145676.1 | 0.002301641 |
| LINC01634 | -0.008772685 |
| NKILA | 0.004137425 |
| GOLM1 | 0.009448779 |
| LINC00862 | 0.008308114 |
| SLF1 | 0.004226585 |
| ZC3H12D | -0.012195406 |
| ZNF563 | -0.000248725 |
| ZCWPW1 | -0.000592242 |
| TRIM6 | 0.003242132 |
| AC105942.1 | -0.002783341 |
| LINC01806 | -0.00351529 |
