## Supplemental Table S2 for "Multiomic Characterization of Stage I Lung Adenocarcinoma Reveals Distinct Genetic and Immunologic Features of Recurrent Disease"

**Table S2. Pathway Analysis for Differential Gene Expression Heatmap Cluster 2**

| Ingenuity Canonical Pathways | -log(p-value) | Ratio | z-score | Molecules |
|---|---|---|---|---|
| Kinetochore Metaphase Signaling Pathway | 2.95E+01 | 4.95E-01 | 3.202 | ANAPC1,ANAPC7,AURKB,BIRC5,BUB1,BUB1B,BUB3,CCNB1,CDC20,CDC27,CDCA8,CDK1,CENPA,CENPE,CENPH,CENPK,CENPL,CENPN,CENPO,CENPP,CENPU,CENPW,DSN1,ESPL1,H2AX,H2AZ1,INCENP,KIF2C,KNL1,KNTC1,MAD2L1,MASTL,NDC80,NEK2,NUF2,PLK1,PPP1CC,PPP1R12A,PTTG1,RAD21,SKA1,SKA2,SKA3,SMC1B,SMC3,SPC24,SPC25,SPDL1,TTK,ZW10,ZWILCH,ZWINT |
| Cell Cycle Control of Chromosomal Replication | 1.85E+01 | 5.36E-01 | 5.477 | CDC45,CDC6,CDC7,CDK1,CDK12,CDK16,CDK2,CDK6,CDK8,CDT1,CHEK2,DBF4,DNA2,LIG1,MCM2,MCM3,MCM4,MCM5,MCM6,MCM7,MCM8,ORC1,ORC6,PCNA,POLA2,POLD1,POLE,PRIM1,PRIM2,TOP2A |
| Role of BRCA1 in DNA Damage Response | 1.78E+01 | 4.38E-01 | 1.961 | ACTB,BARD1,BLM,BRCA1,BRCA2,BRIP1,CHEK1,CHEK2,DPF1,E2F1,E2F2,E2F3,E2F7,E2F8,FANCA,FANCB,FANCC,FANCD2,FANCG,FANCL,FANCM,HLTF,MSH2,MSH6,NBN,PLK1,RAD51,RBBP8,RBL1,RFC2,RFC3,RFC4,RFC5,STAT1,TOPBP1 |
| Mitotic Roles of Polo-Like Kinase | 1.12E+01 | 3.79E-01 | 2.5 | ANAPC1,ANAPC7,CCNB1,CCNB2,CDC20,CDC25A,CDC25C,CDC27,CDC7,CDK1,CHEK2,ESPL1,FBXO5,HSP90AA1,HSP90B1,KIF11,KIF23,PKMYT1,PLK1,PLK4,PPP2R1B,PRC1,PTTG1,RAD21,SMC3 |
| Hereditary Breast Cancer Signaling | 1.12E+01 | 2.68E-01 | NaN | ACTB,BARD1,BLM,BRCA1,BRCA2,CCNB1,CDC25C,CDK1,CDK6,CHEK1,CHEK2,DPF1,E2F1,FANCA,FANCB,FANCC,FANCD2,FANCG,FANCL,FANCM,H2AX,HDAC2,HLTF,KRAS,MSH2,MSH6,NBN,NRAS,PALB2,RAD51,RALA,RAP2A,RAP2B,RFC2,RFC3,RFC4,RFC5,TUBG1 |
| Role of CHK Proteins in Cell Cycle Checkpoint Control | 9.28E+00 | 3.68E-01 | -0.535 | BRCA1,CDC25A,CDC25C,CDK1,CDK2,CHEK1,CHEK2,CLSPN,E2F1,E2F2,E2F3,E2F7,E2F8,NBN,PCNA,PLK1,PPP2R1B,RFC2,RFC3,RFC4,RFC5 |
| Estrogen-mediated S-phase Entry | 9.01E+00 | 5.38E-01 | 3.051 | CCNA2,CCNE1,CCNE2,CDC25A,CDK1,CDK2,E2F1,E2F2,E2F3,E2F7,E2F8,RBL1,SKP2,TFDP1 |
| RAN Signaling | 8.36E+00 | 6.47E-01 | 3.317 | CSE1L,IPO5,KPNA1,KPNA2,KPNA3,KPNA4,KPNB1,RANBP1,RANGAP1,RCC1,XPO1 |
| ATM Signaling | 7.92E+00 | 2.68E-01 | 1.043 | BLM,BRCA1,CBX1,CBX3,CCNB1,CCNB2,CDC25A,CDC25C,CDK1,CDK2,CHEK1,CHEK2,FANCD2,H2AX,NBN,PPP1CC,PPP2R1B,RAD51,RBBP8,SMC1B,SMC2,SMC3,SUV39H1,TDP1,TOPBP1,TRIM28 |
| Cell Cycle: G2/M DNA Damage Checkpoint Regulation | 7.86E+00 | 3.60E-01 | -1.069 | AURKA,BORA,BRCA1,CCNB1,CCNB2,CDC25C,CDC34,CDK1,CHEK1,CHEK2,PKMYT1,PLK1,PRKDC,SKP2,TOP2A,YWHAG,YWHAH,YWHAZ |
| Mismatch Repair in Eukaryotes | 7.43E+00 | 6.25E-01 | NaN | EXO1,FEN1,MSH2,MSH6,PCNA,POLD1,RFC2,RFC3,RFC4,RFC5 |
| Cyclins and Cell Cycle Regulation | 7.27E+00 | 2.74E-01 | 3.153 | CCNA2,CCNB1,CCNB2,CCNE1,CCNE2,CDC25A,CDC34,CDK1,CDK2,CDK6,CDKN2D,E2F1,E2F2,E2F3,E2F7,E2F8,HDAC2,PA2G4,PPP2R1B,RBL1,SKP2,SUV39H1,TFDP1 |
| Remodeling of Epithelial Adherens Junctions | 6.97E+00 | 2.94E-01 | 1 | ACTB,ACTG1,ACTN1,ACTN4,ACTR2,ACTR3,ARF6,CBLL1,CLIP1,DNM1L,MAPRE1,RALA,TUBA1B,TUBA1C,TUBB,TUBB3,TUBB6,TUBG1,VCL,ZYX |
| Cell Cycle: G1/S Checkpoint Regulation | 6.97E+00 | 2.94E-01 | -1.5 | CCNE1,CCNE2,CDC25A,CDC34,CDK2,CDK6,CDKN2D,E2F1,E2F2,E2F3,E2F7,E2F8,GNL3,HDAC2,PA2G4,PAK1IP1,RBL1,SKP2,SUV39H1,TFDP1 |
| Epithelial Adherens Junction Signaling | 6.95E+00 | 2.09E-01 | NaN | ACTB,ACTG1,ACTN1,ACTN4,ACTR2,ACTR3,CDH2,CLIP1,JUP,KRAS,NECTIN2,NOTCH3,NRAS,PARD3,RAC1,RALA,RAP2A,RAP2B,SNAI1,SNAI2,SSX2IP,TCF3,TUBA1B,TUBA1C,TUBB,TUBB3,TUBB6,TUBG1,VAV2,VCL,WASF1,YES1,ZYX |
| Molecular Mechanisms of Cancer | 6.37E+00 | 1.46E-01 | NaN | ARHGEF4,ARHGEF5,AURKA,BCL2L11,BMP1,BMP8A,BRCA1,CCNE1,CCNE2,CDC25A,CDC25C,CDK1,CDK12,CDK16,CDK2,CDK6,CDK8,CDKN2D,CHEK1,CHEK2,E2F1,E2F2,E2F3,E2F7,E2F8,ELK1,FADD,FANCD2,GNA12,GNA13,GNAI3,GNG4,GSK3A,HAT1,HDAC2,HIF1A,IRS1,ITGA11,ITGA5,ITGAV,ITGB1,ITGB5,ITGB8,KRAS,NBN,NRAS,PA2G4,PAK1,PAK2,PLCB3,PRKDC,PTPN11,RAC1,RALA,RALBP1,RAP2A,RAP2B,RBL1,RHOF,RHOV,RND3,SHC1,SUV39H1,TCF3,TFDP1 |
| Protein Ubiquitination Pathway | 6.32E+00 | 1.67E-01 | NaN | AMFR,ANAPC1,BRCA1,CDC20,CDC34,CUL2,DNAJA1,DNAJB11,DNAJB6,DNAJC10,DNAJC13,DNAJC2,DNAJC22,DNAJC9,HSP90AA1,HSP90B1,HSPA13,HSPA1A/HSPA1B,HSPD1,HSPH1,PSMB2,PSMC1,PSMD1,PSMD11,PSMD12,PSMD2,PSMD3,PSMD7,SASS6,SKP2,THOP1,UBE2C,UBE2E3,UBE2H,UBE2K,UBE2R2,UBE2S,UBE2T,UBE2V1,UBE2V2,UCHL3,USP1,USP14,USP31,USP32,USP42 |
| Regulation of Cellular Mechanics by Calpain Protease | 6.14E+00 | 2.47E-01 | 1.897 | ACTN1,ACTN4,CCNA2,CCNE1,CCNE2,CDK1,CDK2,CDK6,CNGB1,ITGA11,ITGA5,ITGAV,ITGB1,ITGB5,ITGB8,KRAS,NRAS,PXN,RALA,RAP2A,RAP2B,VCL |
| Rac Signaling | 5.73E+00 | 2.03E-01 | 4.796 | ACTR2,ACTR3,CDK5R1,CFL1,ELK1,IQGAP3,ITGA11,ITGA5,ITGAV,ITGB1,ITGB5,ITGB8,KRAS,LIMK1,NCKAP1,NRAS,PAK1,PAK2,PARD3,PIP4K2A,PIP4K2C,PLD1,RAC1,RALA,RAP2A,RAP2B,RPS6KB1,WASF1 |
| Germ Cell-Sertoli Cell Junction Signaling | 5.65E+00 | 1.87E-01 | NaN | ACTB,ACTG1,ACTN1,ACTN4,CDH2,CFL1,ITGB1,JUP,KRAS,LIMK1,MAP3K10,MTMR2,NECTIN2,NRAS,PAK1,PAK2,PXN,RAC1,RALA,RAP2A,RAP2B,RHOF,RHOV,RND3,TUBA1B,TUBA1C,TUBB,TUBB3,TUBB6,TUBG1,VCL,ZYX |
| Ephrin Receptor Signaling | 5.37E+00 | 1.74E-01 | 4.796 | ACTR2,ACTR3,ADAM10,CFL1,EFNA3,EPHA1,EPHB2,EPHB4,GNA12,GNA13,GNAI3,GNG4,GRIN2D,ITGA11,ITGA5,ITGAV,ITGB1,ITGB5,ITGB8,KRAS,LIMK1,MAP4K4,NGEF,NRAS,PAK1,PAK2,PGF,PTPN11,PXN,RAC1,RALA,RAP2A,RAP2B,SHC1,VEGFC |
| Actin Nucleation by ARP-WASP Complex | 5.21E+00 | 2.26E-01 | 3.638 | ACTR2,ACTR3,GNA12,ITGA11,ITGA5,ITGAV,ITGB1,ITGB5,ITGB8,KRAS,NRAS,PPP1R12A,RAC1,RALA,RAP2A,RAP2B,RHOF,RHOV,RND3,VASP,WASF1 |
| Pancreatic Adenocarcinoma Signaling | 5.01E+00 | 1.98E-01 | 2.714 | BIRC5,BRCA2,CCNE1,CCNE2,CDK2,E2F1,E2F2,E2F3,E2F7,E2F8,ELK1,HDAC2,KRAS,PA2G4,PGF,PLD1,RAC1,RAD51,RALA,RALBP1,RBL1,STAT1,SUV39H1,TFDP1,VEGFC |
| Actin Cytoskeleton Signaling | 4.94E+00 | 1.59E-01 | 4.596 | ACTB,ACTG1,ACTN1,ACTN4,ACTR2,ACTR3,ARHGEF4,CFL1,DIAPH3,FGF11,FN1,GNA12,GNA13,IQGAP3,ITGA11,ITGA5,ITGAV,ITGB1,ITGB5,ITGB8,KRAS,LIMK1,MYLK2,NCKAP1,NRAS,PAK1,PAK2,PFN2,PPP1R12A,PXN,RAC1,RALA,RAP2A,RAP2B,SHC1,TRIO,VAV2,VCL,WASF1 |
| Oncostatin M Signaling | 4.89E+00 | 3.02E-01 | 3.606 | ELK1,KRAS,MMP1,MMP3,MT2A,NRAS,OSMR,PLAU,RALA,RAP2A,RAP2B,SHC1,STAT1 |
| UDP-N-acetyl-D-glucosamine Biosynthesis II | 4.82E+00 | 8.33E-01 | 2.236 | GFPT1,GFPT2,GNPNAT1,PGM3,UAP1 |
| Integrin Signaling | 4.80E+00 | 1.64E-01 | 4.95 | ACTB,ACTG1,ACTN1,ACTN4,ACTR2,ACTR3,ARF6,ASAP1,BCAR3,ITGA11,ITGA5,ITGAV,ITGB1,ITGB5,ITGB8,KRAS,MYLK2,NRAS,PAK1,PAK2,PFN2,PPP1R12A,PXN,RAC1,RALA,RAP2A,RAP2B,RHOF,RHOV,RND3,SHC1,TSPAN5,VASP,VCL,ZYX |

| | | | | |
|---|---|---|---|---|
| Axonal Guidance Signaling | 4.76E+00 | 1.30E-01 | NaN | ACTR2,ACTR3,ADAM10,ADAM12,ADAM17,ADAM32,ADAM9,ADAMTS12,ADAMTS2,ADAMTS4,ADAMTS5,ADAMTS6,BMP1,BMP8A,CFL1,EFNA3,EPHA1,EPHB2,EPHB4,GNA12,GNA13,GNAI3,GNG4,ITGA11,ITGA5,ITGAV,ITGB1,ITGB5,ITGB8,KRAS,LIMK1,MMP1,MMP11,MMP12,MMP14,MMP3,NGEF,NRAS,PAK1,PAK2,PFN2,PGF,PLCB3,PLCD1,PLCD3,PLXNA1,PTPN11,PXN,RAC1,RALA,RAP2A,RAP2B,SEMA3A,SEMA3C,SEMA4B,SEMA7A,SHC1,SRGAP1,TUBA1B,TUBA1C,TUBB,TUBB3,TUBB6,TUBG1,VASP,VEGFC |
| Paxillin Signaling | 4.67E+00 | 2.04E-01 | 3.771 | ACTB,ACTG1,ACTN1,ACTN4,ARF6,ITGA11,ITGA5,ITGAV,ITGB1,ITGB5,ITGB8,KRAS,NRAS,PAK1,PAK2,PTPN12,PXN,RAC1,RALA,RAP2A,RAP2B,VCL |
| RhoA Signaling | 4.65E+00 | 1.94E-01 | 3.13 | ACTB,ACTG1,ACTR2,ACTR3,ANLN,CDC42EP2,CFL1,EPHA1,GNA12,GNA13,LIMK1,MYLK2,NGEF,PFN2,PIP4K2A,PIP4K2C,PLD1,PLXNA1,PPP1R12A,RHPN2,RND3,RTKN,SEPTIN11,WASF1 |
| Ferroptosis Signaling Pathway | 4.53E+00 | 1.90E-01 | 0.408 | ANGPTL4,ARF6,CARS1,FANCD2,GCH1,GCLC,GSS,H2AX,H2AZ1,H2BC8,KRAS,NRAS,PRKAA2,PRKAB2,RALA,RAP2A,RAP2B,RBL1,SLC38A1,SLC39A14,SLC3A2,SLC7A11,TFRC,TXNRD1 |
| NER (Nucleotide Excision Repair, Enhanced Pathway) | 4.49E+00 | 2.04E-01 | 3 | CHAF1A,CHAF1B,DNA2,GTF2H3,H4C8,H4C9,LIG1,PCNA,POLA2,POLD1,POLD2,POLE,POLE2,PRIM1,PRIM2,RAD23B,RFC2,RFC3,RFC4,RFC5,TOP2A |
| Signaling by Rho Family GTPases | 4.38E+00 | 1.49E-01 | 4.642 | ACTB,ACTG1,ACTR2,ACTR3,ARHGEF4,ARHGEF5,CDC42EP2,CDH2,CDH24,CDH3,CFL1,CLIP1,DIAPH3,ELK1,GNA12,GNA13,GNAI3,GNG4,ITGA11,ITGA5,ITGAV,ITGB1,ITGB5,ITGB8,LIMK1,MAP3K10,PAK1,PAK2,PARD3,PIP4K2A,PIP4K2C,PLD1,PPP1R12A,RAC1,RHOF,RHOV,RND3,SEPTIN11,STMN1,WASF1 |
| Regulation of Actin-based Motility by Rho | 4.17E+00 | 1.90E-01 | 3.638 | ACTB,ACTR2,ACTR3,CFL1,ITGA11,ITGA5,ITGAV,ITGB1,ITGB5,ITGB8,LIMK1,PAK1,PAK2,PFN2,PIP4K2A,PIP4K2C,PPP1R12A,RAC1,RHOF,RHOV,RND3,WASF1 |
| Cell Cycle Regulation by BTG Family Proteins | 4.15E+00 | 2.97E-01 | 3 | CCNE1,CCNE2,CDK2,E2F1,E2F2,E2F3,E2F7,E2F8,HOXB9,NOCT,PPP2R1B |
| Sumoylation Pathway | 4.00E+00 | 1.94E-01 | -1.807 | DNMT3A,HDAC2,KDM1A,PCNA,RAC1,RANGAP1,RCC1,RFC2,RFC3,RFC4,RFC5,RHOF,RHOV,RND3,SAE1,SENP1,SENP2,SENP5,SERBP1,TDG |
| Sertoli Cell-Sertoli Cell Junction Signaling | 3.96E+00 | 1.55E-01 | NaN | ACTB,ACTG1,ACTN1,ACTN4,CLDN14,ELK1,GSK3A,ITGA11,ITGA5,ITGAV,ITGB1,ITGB5,ITGB8,JUP,KRAS,MAP3K10,MTMR2,NECTIN2,NOS1,NRAS,RAC1,RALA,RAP2A,RAP2B,SPTBN2,TUBA1B,TUBA1C,TUBB,TUBB3,TUBB6,TUBG1,VCL |
| Osteoarthritis Pathway | 3.94E+00 | 1.50E-01 | 2.785 | ACAN,ADAMTS4,ADAMTS5,CASP2,CCN4,DKK1,FADD,FN1,GREM1,H19,HIF1A,IL1R2,IL1RAP,ITGA11,ITGA5,ITGAV,ITGB1,ITGB5,ITGB8,MMP1,MMP12,MMP3,NAMPT,PGF,PPARD,PRKAA2,PRKAB2,PTHLH,RAC1,RUNX2,S100A8,S100A9,SPHK1,TCF3,VEGFC |
| tRNA Charging | 3.92E+00 | 2.82E-01 | 3.317 | CARS1,DARS2,FARSB,GARS1,IARS1,MARS1,NARS1,RARS1,TARS1,VARS1,YARS2 |
| Tetrahydrofolate Salvage from 5,10-methenyltetrahydrofolate | 3.79E+00 | 8.00E-01 | 2 | GART,MTHFD1,MTHFD1L,MTHFD2 |
| Bladder Cancer Signaling | 3.72E+00 | 1.81E-01 | 2.449 | E2F1,E2F2,E2F3,FGF11,HDAC2,KRAS,MMP1,MMP11,MMP12,MMP14,MMP3,NRAS,PA2G4,PGF,RALA,RAP2A,RAP2B,RBL1,SUV39H1,TFDP1,VEGFC |
| Neuregulin Signaling | 3.66E+00 | 1.79E-01 | 3.051 | ADAM17,CDK5R1,ELK1,EREG,ERRFI1,HSP90AA1,HSP90B1,ITGA11,ITGA5,ITGAV,ITGB1,ITGB5,ITGB8,KRAS,NRAS,PTPN11,RALA,RAP2A,RAP2B,RPS6KB1,SHC1 |
| DNA damage-induced 14-3-3œÉ Signaling | 3.46E+00 | 3.68E-01 | NaN | BRCA1,CCNB1,CCNB2,CCNE1,CCNE2,CDK1,CDK2 |
| DNA Double-Strand Break Repair by Homologous Recombination | 3.44E+00 | 4.29E-01 | NaN | BRCA1,BRCA2,GEN1,LIG1,NBN,RAD51 |
| Glutathione Biosynthesis | 3.34E+00 | 1.00E+00 | NaN | GCLC,GCLM,GSS |
| Coronavirus Replication Pathway | 3.32E+00 | 2.44E-01 | 3.317 | ARCN1,COPB2,COPG1,COPG2,CTSL,GSK3A,TUBA1B,TUBA1C,TUBB,TUBB3,TUBB6 |
| GADD45 Signaling | 3.30E+00 | 3.50E-01 | NaN | BRCA1,CCNB1,CCNE1,CCNE2,CDK1,CDK2,PCNA |
| VEGF Signaling | 3.29E+00 | 1.82E-01 | 3 | ACTB,ACTG1,ACTN1,ACTN4,EIF2S1,EIF2S2,HIF1A,KRAS,NRAS,PGF,PTPN11,PXN,RALA,RAP2A,RAP2B,SHC1,VCL,VEGFC |
| Role of Tissue Factor in Cancer | 3.28E+00 | 1.72E-01 | NaN | CFL1,GNA12,GNA13,ITGAV,ITGB1,KRAS,LIMK1,MMP1,NRAS,PAK1,PLAUR,PTPN11,RAC1,RALA,RAP2A,RAP2B,RPS6KA4,RPS6KB1,VEGFC,YES1 |
| RhoGDI Signaling | 3.28E+00 | 1.44E-01 | -3.962 | ACTB,ACTG1,ACTR2,ACTR3,ARHGEF4,ARHGEF5,CDH2,CDH24,CDH3,CFL1,GNA12,GNA13,GNAI3,GNG4,ITGA11,ITGA5,ITGAV,ITGB1,ITGB5,ITGB8,LIMK1,PAK1,PAK2,PIP4K2A,PIP4K2C,PPP1R12A,RAC1,RHOF,RHOV,RND3,WASF1 |
| FAK Signaling | 3.23E+00 | 1.71E-01 | NaN | ACTB,ACTG1,ASAP1,HMMR,ITGA11,ITGA5,ITGAV,ITGB1,ITGB5,ITGB8,KRAS,NRAS,PAK1,PAK2,PXN,RAC1,RALA,RAP2A,RAP2B,VCL |
| ILK Signaling | 3.22E+00 | 1.46E-01 | 3.674 | ACTB,ACTG1,ACTN1,ACTN4,CFL1,DSP,FBLIM1,FLNC,FN1,GSK3A,HIF1A,IRS1,ITGB1,ITGB5,ITGB8,KRT18,PGF,PPP1R12A,PPP2R1B,PXN,RAC1,RHOF,RHOV,RND3,RPS6KA4,SNAI1,SNAI2,VCL,VEGFC |
| Senescence Pathway | 3.13E+00 | 1.31E-01 | 0 | ANAPC1,ANAPC7,CCNB1,CCNB2,CCNE1,CCNE2,CDC25A,CDC25C,CDC27,CDK1,CDK2,CDK6,CGAS,CHEK1,CHEK2,DLD,E2F1,E2F2,E2F3,E2F7,E2F8,EED,EIF4EBP1,EZH2,ING1,KRAS,MAPK6,NBN,NRAS,PCGF6,PDK3,PHF19,PPP2R1B,RALA,RAP2A,RAP2B,RBL1,RPS6KA4,SERPINE1 |
| Polyamine Regulation in Colon Cancer | 3.13E+00 | 2.10E-01 | NaN | AZIN1,KRAS,MXD1,PSMB2,PSMC1,PSMD1,PSMD11,PSMD12,PSMD2,PSMD3,PSMD7,PSME3,PSME4 |
| Agrin Interactions at Neuromuscular Junction | 3.11E+00 | 2.00E-01 | 3.464 | ACTB,ACTG1,ITGB1,KRAS,LAMB1,LAMC1,NRAS,PAK1,PAK2,PXN,RAC1,RALA,RAP2A,RAP2B |
| Purine Nucleotides De Novo Biosynthesis II | 3.08E+00 | 4.55E-01 | 2.236 | ATIC,GART,GMPS,PAICS,PPAT |
| Prostate Cancer Signaling | 3.06E+00 | 1.70E-01 | NaN | CCNE1,CCNE2,CDK2,E2F1,E2F2,E2F3,HDAC2,HSP90AA1,HSP90B1,KRAS,NRAS,PA2G4,RALA,RAP2A,RAP2B,RBL1,SRD5A1,SUV39H1,TFDP1 |
| ERK5 Signaling | 2.98E+00 | 1.94E-01 | 3.742 | FOSL1,GNA12,GNA13,KRAS,NRAS,PTPN11,RALA,RAP2A,RAP2B,RPS6KA4,RPS6KB1,YWHAG,YWHAH,YWHAZ |
| p53 Signaling | 2.91E+00 | 1.73E-01 | 0 | BIRC5,BRCA1,CCNK,CDK2,CHEK1,CHEK2,E2F1,GNL3,HIF1A,PCNA,PERP,PRKDC,SERPINB5,SNAI2,TIGAR,TOPBP1,WT1 |
| HIF1Œ± Signaling | 2.87E+00 | 1.39E-01 | 3.78 | ADM,CCNG2,CUL2,EGLN3,EIF4EBP1,GPI,HIF1A,HK2,HSP90AA1,HSPA1A/HSPA1B,KDM1A,KRAS,MMP1,MMP11,MMP12,MMP14,MMP3,NRAS,PGF,PKM,RAC1,RALA,RAP2A,RAP2B,RPS6KB1,SERPINE1,SLC2A1,SLC2A5,VEGFC |
| Pyridoxal 5'-phosphate Salvage Pathway | 2.86E+00 | 1.97E-01 | 3.606 | CDK1,CDK2,CDK6,CDK8,IRAK1,LIMK1,MAPK6,NEK2,PAK1,PAK2,PLK1,PRKAA2,TTK |
| PAK Signaling | 2.78E+00 | 1.61E-01 | 3.207 | CFL1,ITGA11,ITGA5,ITGAV,ITGB1,ITGB5,ITGB8,KRAS,LIMK1,NRAS,PAK1,PAK1IP1,PAK2,PXN,RAC1,RALA,RAP2A,RAP2B,SHC1 |
| 14-3-3-mediated Signaling | 2.78E+00 | 1.57E-01 | 2.496 | ELK1,GSK3A,KRAS,NRAS,PLCB3,PLCD1,PLCD3,RALA,RAP2A,RAP2B,SRPK2,TUBA1B,TUBA1C,TUBB,TUBB3,TUBB6,TUBG1,YWHAG,YWHAH,YWHAZ |

| | | | | |
|---|---|---|---|---|
| Clathrin-mediated Endocytosis Signaling | 2.74E+00 | 1.40E-01 | NaN | ACTB,ACTG1,ACTR2,ACTR3,AP2A1,AP2M1,ARF6,CD2AP,CLTCL1,CSNK2A1,DNM1L,EPHB2,FGF11,ITGA5,ITGB1,ITGB5,ITGB8,MYO1E,PGF,PICALM,RAC1,S100A8,SH3GL1,SNX9,STAM,TFRC,VEGFC |
| UDP-N-acetyl-D-galactosamine Biosynthesis II | 2.69E+00 | 3.85E-01 | 2.236 | GNPNAT1,GPI,HK2,PGM3,UAP1 |
| ERK/MAPK Signaling | 2.68E+00 | 1.36E-01 | 2.132 | DUSP4,EIF4EBP1,ELK1,H3C14,ITGA11,ITGA5,ITGAV,ITGB1,ITGB5,ITGB8,KRAS,NRAS,PAK1,PAK2,PPP1CC,PPP1R12A,PPP2R1B,PXN,RAC1,RALA,RAP2A,RAP2B,RPS6KA4,SHC1,STAT1,VRK2,YWHAG,YWHAH,YWHAZ |
| Inhibition of Matrix Metalloproteases | 2.63E+00 | 2.31E-01 | -2.333 | ADAM10,ADAM12,ADAM17,MMP1,MMP11,MMP12,MMP14,MMP3,THBS2 |
| Virus Entry via Endocytic Pathways | 2.62E+00 | 1.63E-01 | NaN | ACTB,ACTG1,AP2A1,AP2M1,CLTCL1,FLNC,ITGA5,ITGB1,ITGB5,ITGB8,KRAS,NRAS,RAC1,RALA,RAP2A,RAP2B,TFRC |
| IGF-1 Signaling | 2.62E+00 | 1.63E-01 | 3.051 | CSNK2A1,ELK1,IGFBP3,IRS1,KRAS,NRAS,PTPN11,PXN,RALA,RAP2A,RAP2B,RPS6KB1,SHC1,SOCS4,YWHAG,YWHAH,YWHAZ |
| Semaphorin Signaling in Neurons | 2.61E+00 | 1.94E-01 | NaN | CFL1,ITGB1,LIMK1,PAK1,PAK2,PLXNA1,RAC1,RHOF,RHOV,RND3,SEMA3A,SEMA7A |
| Ephrin A Signaling | 2.58E+00 | 2.13E-01 | NaN | ADAM10,CFL1,EFNA3,EPHA1,LIMK1,NGEF,PAK1,PTPN11,RAC1,VAV2 |
| Glycolysis I | 2.55E+00 | 2.69E-01 | 2.646 | ALDOA,ENO1,ENO2,GAPDH,GPI,PFKP,PKM |
| Glioblastoma Multiforme Signaling | 2.52E+00 | 1.40E-01 | 3 | CCNE1,CCNE2,CDK2,CDK6,E2F1,E2F2,E2F3,E2F7,E2F8,KRAS,NRAS,PLCB3,PLCD1,PLCD3,RAC1,RALA,RAP2A,RAP2B,RHOF,RHOV,RND3,RPS6KB1,SHC1,TCF3 |
| Folate Transformations I | 2.49E+00 | 4.44E-01 | 2 | MTHFD1,MTHFD1L,MTHFD2,SHMT2 |
| FAT10 Signaling Pathway | 2.49E+00 | 1.96E-01 | NaN | PSMB2,PSMC1,PSMD1,PSMD11,PSMD12,PSMD2,PSMD3,PSMD7,PSME3,PSME4,UBA6 |
| Salvage Pathways of Pyrimidine Ribonucleotides | 2.49E+00 | 1.63E-01 | 4 | AK4,APOBEC3B,CDK1,CDK2,CDK6,CDK8,IRAK1,LIMK1,MAPK6,NEK2,PAK1,PAK2,PLK1,PRKAA2,TTK,UCK2 |
| Chronic Myeloid Leukemia Signaling | 2.49E+00 | 1.59E-01 | NaN | CDK6,E2F1,E2F2,E2F3,E2F7,E2F8,HDAC2,KRAS,NRAS,PA2G4,PTPN11,RALA,RAP2A,RAP2B,RBL1,SUV39H1,TFDP1 |
| Glioma Invasiveness Signaling | 2.45E+00 | 1.78E-01 | 3.464 | HMMR,ITGAV,KRAS,NRAS,PLAU,PLAUR,RAC1,RALA,RAP2A,RAP2B,RHOF,RHOV,RND3 |
| Coronavirus Pathogenesis Pathway | 2.42E+00 | 1.33E-01 | -2.294 | ADAM17,ADAM9,BCL2L11,CCNE1,CCNE2,CDK2,CTSL,E2F1,E2F2,E2F3,E2F7,E2F8,ELK1,HDAC2,HIF1A,KPNB1,OAS1,OAS3,PA2G4,PTGES2,RBL1,SERPINE1,SIGMAR1,STAT1,SUV39H1,TFDP1,TOMM70 |
| Hypoxia Signaling in the Cardiovascular System | 2.40E+00 | 1.76E-01 | NaN | CDC34,HIF1A,HSP90AA1,HSP90B1,UBE2C,UBE2E3,UBE2H,UBE2K,UBE2R2,UBE2S,UBE2T,UBE2V1,UBE2V2 |
| dTMP De Novo Biosynthesis | 2.39E+00 | 6.00E-01 | NaN | DHFR,SHMT2,TYMS |
| Folate Polyglutamylation | 2.39E+00 | 6.00E-01 | NaN | MTHFD1,MTHFD1L,SHMT2 |
| DNA Methylation and Transcriptional Repression Signaling | 2.36E+00 | 2.29E-01 | NaN | DNMT1,DNMT3A,DNMT3B,H4C8,H4C9,HDAC2,MTA2,SAP30 |
| Caveolar-mediated Endocytosis Signaling | 2.35E+00 | 1.73E-01 | NaN | ACTB,ACTG1,ARCN1,COPB2,COPG1,COPG2,FLNC,ITGA11,ITGA5,ITGAV,ITGB1,ITGB5,ITGB8 |
| Ovarian Cancer Signaling | 2.31E+00 | 1.39E-01 | 2.828 | BRCA1,BRCA2,E2F1,E2F2,E2F3,HDAC2,KRAS,MSH2,MSH6,NRAS,PA2G4,PGF,RAD51,RALA,RAP2A,RAP2B,RBL1,RPS6KB1,SUV39H1,TCF3,TFDP1,VEGFC |
| Macropinocytosis Signaling | 2.30E+00 | 1.71E-01 | 3 | ACTN4,ARF6,ITGA5,ITGB1,ITGB5,ITGB8,KRAS,NRAS,PAK1,RAC1,RALA,RAP2A,RAP2B |
| Regulation of eIF4 and p70S6K Signaling | 2.26E+00 | 1.34E-01 | 2.714 | AGO2,EIF2S1,EIF2S2,EIF3B,EIF3J,EIF4A3,EIF4EBP1,EIF4G1,IRS1,ITGA11,ITGA5,ITGAV,ITGB1,ITGB5,ITGB8,KRAS,NRAS,PAIP1,PPP2R1B,RALA,RAP2A,RAP2B,RPS6KB1,SHC1 |
| Tumor Microenvironment Pathway | 2.26E+00 | 1.34E-01 | 4.796 | COL1A1,COL1A2,COL3A1,FGF11,FN1,HIF1A,ITGA5,KRAS,MMP1,MMP11,MMP12,MMP14,MMP3,NRAS,PGF,PLAU,RAC1,RALA,RAP2A,RAP2B,SLC16A1,SLC2A1,TNC,VEGFC |
| PI3K/AKT Signaling | 2.25E+00 | 1.31E-01 | 2.668 | EIF4EBP1,GSK3A,HSP90AA1,HSP90B1,IL12RB2,IL17RD,IL1R2,IL20RB,ITGA11,ITGA5,ITGAV,ITGB1,ITGB5,ITGB8,KRAS,NRAS,PPP2R1B,RALA,RAP2A,RAP2B,RPS6KB1,SHC1,SYNJ2,YWHAG,YWHAH,YWHAZ |
| Uridine-5'-phosphate Biosynthesis | 2.23E+00 | 1.00E+00 | NaN | CAD,UMPS |
| Glioma Signaling | 2.17E+00 | 1.45E-01 | 2.449 | CDK6,CDKN2D,E2F1,E2F2,E2F3,E2F7,E2F8,HDAC2,KRAS,NRAS,PA2G4,RALA,RAP2A,RAP2B,RBL1,SHC1,SUV39H1,TFDP1 |
| HOTAIR Regulatory Pathway | 2.15E+00 | 1.35E-01 | 4.025 | AGO2,COL1A1,COL1A2,COL3A1,DNMT3B,EED,EZH2,FOXM1,H3C14,HDAC2,JARID2,KDM1A,MMP1,MMP11,MMP12,MMP14,MMP3,MTF2,SNAI2,SUZ12,TCF3,TWIST1 |
| D-myo-inositol-5-phosphate Metabolism | 2.14E+00 | 1.30E-01 | 4.264 | CDC25A,CDC25C,DUSP14,DUSP4,HACD2,MTMR2,NOCT,NUDT1,NUDT15,PAWR,PGAM5,PIP4K2A,PIP4K2C,PLCB3,PLCD1,PLCD3,PPP1CC,PPP1R12A,PPP2R1B,PTPN11,PTPN12,PTPRH,PTPRN,SET,WBP11 |
| Aldosterone Signaling in Epithelial Cells | 2.12E+00 | 1.34E-01 | 1.633 | DNAJA1,DNAJB11,DNAJB6,DNAJC10,DNAJC13,DNAJC2,DNAJC22,DNAJC9,HSP90AA1,HSP90B1,HSPA13,HSPA1A/HSPA1B,HSPD1,HSPH1,KCNMB4,KRAS,PIP4K2A,PIP4K2C,PLCB3,PLCD1,PLCD3,SASS6 |
| Renal Cell Carcinoma Signaling | 2.11E+00 | 1.63E-01 | 3 | CUL2,EGLN3,HIF1A,KRAS,NRAS,PAK1,PAK2,PTPN11,RAC1,RALA,RAP2A,RAP2B,SLC2A1 |
| Reelin Signaling in Neurons | 2.10E+00 | 1.43E-01 | 3 | ACTR2,ACTR3,ARHGEF4,ARHGEF5,CDH2,CDK5R1,CFL1,GRIN2D,ITGA5,ITGB1,LIMK1,MAP3K10,PAFAH1B2,PAFAH1B3,PDK3,RAC1,WASF1,YES1 |
| Ephrin B Signaling | 2.07E+00 | 1.67E-01 | 3.162 | CFL1,EPHB2,EPHB4,GNA12,GNA13,GNAI3,GNG4,LIMK1,PAK1,PXN,RAC1,VAV2 |
| Natural Killer Cell Signaling | 1.98E+00 | 1.26E-01 | 2.2 | CFL1,COL1A1,COL1A2,COL3A1,COL5A3,HSPA1A/HSPA1B,IL12RB2,ITGB1,KIR2DL4,KRAS,LIMK1,MAP3K10,NECTIN2,NRAS,PAK1,PAK2,PTPN11,PVR,PXN,RAC1,RALA,RAP2A,RAP2B,ULBP2,VAV2 |
| Cleavage and Polyadenylation of Pre-mRNA | 1.98E+00 | 3.33E-01 | NaN | CPSF2,CPSF3,CSTF2,CSTF3 |
| BER pathway | 1.98E+00 | 3.33E-01 | NaN | FEN1,LIG1,PCNA,POLE |
| PTEN Signaling | 1.95E+00 | 1.33E-01 | -2.5 | BCL2L11,CSNK2A1,GSK3A,ITGA11,ITGA5,ITGAV,ITGB1,ITGB5,ITGB8,KRAS,MAST2,NRAS,RAC1,RALA,RAP2A,RAP2B,RPS6KB1,SHC1,SYNJ2,YWHAH |
| Spliceosomal Cycle | 1.94E+00 | 1.84E-01 | 3 | CTNNBL1,DHX15,EFTUD2,EIF4A3,PLRG1,PRPF19,SF3B4,U2AF1/U2AF1L5,U2AF2 |
| IL-8 Signaling | 1.93E+00 | 1.23E-01 | 4.899 | ANGPT2,EIF4EBP1,GNA12,GNA13,GNAI3,GNG4,IRAK1,ITGAV,KRAS,LASP1,LIMK1,MAP4K4,NRAS,PAK2,PGF,PLD1,RAC1,RALA,RAP2A,RAP2B,RHOF,RHOV,RND3,RPS6KB1,VASP,VEGFC |
| Mechanisms of Viral Exit from Host Cells | 1.93E+00 | 1.95E-01 | NaN | ACTB,ACTG1,LMNB1,LMNB2,SH3GL1,SNF8,VPS37C,XPO1 |
| Apelin Liver Signaling Pathway | 1.91E+00 | 2.31E-01 | 2.449 | COL1A1,COL1A2,COL3A1,COL5A3,GSK3A,IRS1 |
| HGF Signaling | 1.90E+00 | 1.36E-01 | 3.742 | CDK2,ELK1,ITGA11,ITGA5,ITGAV,ITGB1,ITGB5,ITGB8,KRAS,MAP3K10,NRAS,PAK1,PTPN11,PXN,RAC1,RALA,RAP2A,RAP2B |

| | | | | |
|---|---|---|---|---|
| Non-Small Cell Lung Cancer Signaling | 1.90E+00 | 1.49E-01 | 2.449 | CDK6,E2F1,E2F2,E2F3,HDAC2,KRAS,NRAS,PA2G4,RALA,RAP2A,RAP2B,RBL1,SUV39H1,TFDP1 |
| Superpathway of Serine and Glycine Biosynthesis I | 1.90E+00 | 4.29E-01 | NaN | PSAT1,PSPH,SHMT2 |
| Superpathway of Inositol Phosphate Compounds | 1.90E+00 | 1.20E-01 | 4.6 | CDC25A,CDC25C,DUSP14,DUSP4,HACD2,IPMK,IPPK,MTMR2,NOCT,NUDT1,NUDT15,PAWR,PGAM5,PIP4K2A,PIP4K2C,PLCB3,PLCD1,PLCD3,PPP1CC,PPP1R12A,PPP2R1B,PTPN11,PTPN12,PTPRH,PTPRN,SET,SYNJ2,WBP11 |
| HIPPO signaling | 1.89E+00 | 1.53E-01 | -2.121 | CSNK1E,FRMD6,ITCH,LLGL1,PARD3,PPP1CC,PPP1R12A,PPP2R1B,SKP2,TEAD4,YWHAG,YWHAH,YWHAZ |
| Role of JAK2 in Hormone-like Cytokine Signaling | 1.87E+00 | 2.06E-01 | NaN | HLTF,IRS1,PTPN11,SH2B2,SHC1,SOCS4,STAT1 |
| Small Cell Lung Cancer Signaling | 1.82E+00 | 1.46E-01 | 1 | CCNE1,CCNE2,CDK2,CDK6,E2F1,E2F2,E2F3,HDAC2,NOS1,PA2G4,RBL1,SKP2,SUV39H1,TFDP1 |
| Synaptogenesis Signaling Pathway | 1.80E+00 | 1.12E-01 | 4.564 | ACTR2,ACTR3,AP2A1,AP2M1,CDH2,CDH24,CDH3,CFL1,EFNA3,EIF4EBP1,EPHA1,EPHB2,EPHB4,GRIN2D,GRM8,KRAS,LIMK1,MARCKS,NLGN2,NLGN4X,NRAS,PAK1,RAC1,RALA,RAP2A,RAP2B,RPS6KB1,SHC1,STX1A,STXBP5,SYT5,THBS2,WASF1,YES1,YKT6 |
| 5-aminoimidazole Ribonucleotide Biosynthesis I | 1.77E+00 | 6.67E-01 | NaN | GART,PPAT |
| Inosine-5'-phosphate Biosynthesis II | 1.77E+00 | 6.67E-01 | NaN | ATIC,PAICS |
| 1D-myo-inositol Hexakisphosphate Biosynthesis V (from Ins(1,3,4)P3) | 1.77E+00 | 6.67E-01 | NaN | IPMK,IPPK |
| UVA-Induced MAPK Signaling | 1.75E+00 | 1.43E-01 | 2.53 | KRAS,NRAS,PARP12,PARP2,PLCB3,PLCD1,PLCD3,RALA,RAP2A,RAP2B,RPS6KA4,RPS6KB1,SMPD4,STAT1 |
| GP6 Signaling Pathway | 1.75E+00 | 1.34E-01 | 3 | ADAM10,COL11A1,COL12A1,COL15A1,COL1A1,COL1A2,COL3A1,COL5A1,COL5A2,COL5A3,COL7A1,GSK3A,LAMB1,LAMC1,LAMC2,RAC1,VAV2 |
| Pyrimidine Ribonucleotides De Novo Biosynthesis | 1.74E+00 | 1.94E-01 | 2.646 | AK4,CAD,CANT1,CTPS1,ENTPD7,NUDT15,UMPS |
| FLT3 Signaling in Hematopoietic Progenitor Cells | 1.72E+00 | 1.50E-01 | 2.887 | EIF4EBP1,ELK1,KRAS,NRAS,PTPN11,RALA,RAP2A,RAP2B,RPS6KA4,RPS6KB1,SHC1,STAT1 |
| Histidine Degradation III | 1.72E+00 | 3.75E-01 | NaN | MTHFD1,MTHFD1L,MTHFD2 |
| Purine Ribonucleosides Degradation to Ribose-1-phosphate | 1.72E+00 | 3.75E-01 | NaN | ADA,PGM2,PNP |
| p70S6K Signaling | 1.60E+00 | 1.29E-01 | 2.673 | GNAI3,IRS1,KRAS,NRAS,PLCB3,PLCD1,PLCD3,PLD1,PPP2R1B,RALA,RAP2A,RAP2B,RPS6KB1,SHC1,YWHAG,YWHAH,YWHAZ |
| NRF2-mediated Oxidative Stress Response | 1.58E+00 | 1.14E-01 | 3 | ACTB,ACTG1,BACH1,CDC34,DNAJA1,DNAJB11,DNAJB6,DNAJC10,DNAJC13,DNAJC9,FOSL1,GCLC,GCLM,HSP90AA1,HSP90B1,KRAS,NRAS,RALA,RAP2A,RAP2B,SCARB1,SLC35A2,STIP1,TXNRD1,UBE2E3,UBE2K,USP14 |
| BAG2 Signaling Pathway | 1.57E+00 | 1.43E-01 | 0 | HSP90AA1,HSPA1A/HSPA1B,PSMB2,PSMC1,PSMD1,PSMD11,PSMD12,PSMD2,PSMD3,PSMD7,PSME3,PSME4 |
| FcŒ≥ Receptor-mediated Phagocytosis in Macrophages and Monocytes | 1.56E+00 | 1.38E-01 | 3.606 | ACTB,ACTG1,ACTR2,ACTR3,ARF6,PAK1,PLD1,PXN,RAC1,RPS6KB1,VASP,VAV2,YES1 |
| Apelin Cardiac Fibroblast Signaling Pathway | 1.55E+00 | 2.17E-01 | -0.447 | ANGPT2,PRKAA2,PRKAB2,SERPINE1,SPHK1 |
| Estrogen Receptor Signaling | 1.54E+00 | 1.04E-01 | 3.244 | ATP5F1B,CARM1,CDK8,CFL1,EIF4EBP1,FBXO32,GNA12,GNA13,GNAI3,GNG4,GSK3A,HIF1A,HSP90AA1,HSP90B1,KRAS,LIMK1,MMP1,MMP11,MMP12,MMP14,MMP3,NRAS,NRIP1,PAK1,PCNA,PGF,PLCB3,PLCD1,PLCD3,PPP1R12A,PRKAA2,PRKAB2,PRKDC,RALA,RAP2A,RAP2B,RPS6KB1,RUNX2,SHC1,SNAI1,TBL1XR1,VEGFC |
| CNTF Signaling | 1.54E+00 | 1.58E-01 | 3 | KRAS,NRAS,PTPN11,RALA,RAP2A,RAP2B,RPS6KA4,RPS6KB1,STAT1 |
| Granzyme B Signaling | 1.52E+00 | 2.50E-01 | 2 | GZMB,LMNB1,LMNB2,PRKDC |
| CCR3 Signaling in Eosinophils | 1.52E+00 | 1.26E-01 | 3.162 | CCL11,CFL1,GNA12,GNA13,GNAI3,GNG4,KRAS,LIMK1,NRAS,PAK1,PAK2,PLCB3,PPP1R12A,RAC1,RALA,RAP2A,RAP2B |
| D-myo-inositol (1,4,5,6)-Tetrakisphosphate Biosynthesis | 1.50E+00 | 1.19E-01 | 4.243 | CDC25A,CDC25C,DUSP14,DUSP4,HACD2,IPMK,MTMR2,NOCT,NUDT1,NUDT15,PAWR,PGAM5,PPP1CC,PPP1R12A,PPP2R1B,PTPN11,PTPN12,PTPRH,PTPRN,SET,WBP11 |
| D-myo-inositol (3,4,5,6)-tetrakisphosphate Biosynthesis | 1.50E+00 | 1.19E-01 | 4.243 | CDC25A,CDC25C,DUSP14,DUSP4,HACD2,IPMK,MTMR2,NOCT,NUDT1,NUDT15,PAWR,PGAM5,PPP1CC,PPP1R12A,PPP2R1B,PTPN11,PTPN12,PTPRH,PTPRN,SET,WBP11 |
| Proline Biosynthesis I | 1.49E+00 | 5.00E-01 | NaN | ALDH18A1,PYCR1 |
| Oxidized GTP and dGTP Detoxification | 1.49E+00 | 5.00E-01 | NaN | NUDT1,RUVBL2 |
| Antiproliferative Role of Somatostatin Receptor 2 | 1.48E+00 | 1.43E-01 | 2.333 | ELK1,GAD1,GNG4,KRAS,NOS1,NRAS,PTPN11,RALA,RAP2A,RAP2B,SSTR2 |
| 3-phosphoinositide Degradation | 1.47E+00 | 1.16E-01 | 4.359 | CDC25A,CDC25C,DUSP14,DUSP4,HACD2,INPP4B,MTMR2,NOCT,NUDT1,NUDT15,PAWR,PGAM5,PPP1CC,PPP1R12A,PPP2R1B,PTPN11,PTPN12,PTPRH,PTPRN,SET,SYNJ2,WBP11 |
| Breast Cancer Regulation by Stathmin1 | 1.46E+00 | 9.80E-02 | 4.371 | ADGRE2,ADGRF4,ADGRG3,ARHGEF4,ARHGEF5,AURKA,AURKB,AVPR1A,BDKRB1,BDKRB2,CDK1,CDK2,CDK6,CELSR3,E2F1,E2F2,E2F3,E2F7,E2F8,FOXM1,GNA13,GNG4,GPR1,GPR157,GPR158,GPR176,GPR180,GPR19,GPR37,GPR37L1,GPR89A/GPR89B,GRM8,HIF1A,HTR1D,KRAS,NRAS,PAK1,PGF,PLCB3,PPP1CC,PPP1R12A,PPP2R1B,RAC1,RALA,RAP2A,RAP2B,RPS6KB1,SHC1,SLC52A2,SSTR2,STMN1,TUBA1B,TUBA1C,TUBB,TUBB3,TUBB6,TUBG1,VEGFC |
| Calcium Transport I | 1.44E+00 | 3.00E-01 | NaN | ATP2A2,ATP2B1,ATP2C1 |
| Aryl Hydrocarbon Receptor Signaling | 1.43E+00 | 1.19E-01 | 2.5 | ALDH18A1,ALDH1L2,ALDH3B2,CCNA2,CCNE1,CCNE2,CDK2,CDK6,CHEK1,CHEK2,DHFR,E2F1,HSP90AA1,HSP90B1,MCM7,NRIP1,RBL1,SLC35A2,TFDP1 |
| mTOR Signaling | 1.43E+00 | 1.13E-01 | 2.828 | EIF3B,EIF3J,EIF4A3,EIF4EBP1,EIF4G1,HIF1A,IRS1,KRAS,NRAS,PGF,PLD1,PPP2R1B,PRKAA2,PRKAB2,RAC1,RALA,RAP2A,RAP2B,RHOF,RHOV,RND3,RPS6KA4,RPS6KB1,VEGFC |
| Cholecystokinin/Gastrin-mediated Signaling | 1.40E+00 | 1.26E-01 | 3.742 | ELK1,GNA12,GNA13,KRAS,NRAS,PLCB3,PXN,RAC1,RALA,RAP2A,RAP2B,RHOF,RHOV,RND3,SHC1 |
| 3-phosphoinositide Biosynthesis | 1.40E+00 | 1.13E-01 | 4.359 | CDC25A,CDC25C,DUSP14,DUSP4,HACD2,IPMK,MTMR2,NOCT,NUDT1,NUDT15,PAWR,PGAM5,PIP4K2A,PIP4K2C,PPP1CC,PPP1R12A,PPP2R1B,PTPN11,PTPN12,PTPRH,PTPRN,SET,WBP11 |
| Inhibition of ARE-Mediated mRNA Degradation Pathway | 1.39E+00 | 1.18E-01 | 0.905 | AGO2,CNOT9,EXOSC2,MAPK6,PPP2R1B,PSMB2,PSMC1,PSMD1,PSMD11,PSMD12,PSMD2,PSMD3,PSMD7,PSME3,PSME4,TNFSF11,YWHAG,YWHAH,YWHAZ |
| PPARŒ±/RXRŒ± Activation | 1.38E+00 | 1.14E-01 | -2.668 | ADIPOR2,CAND1,CKAP5,GPD2,HSP90AA1,HSP90B1,IL1R2,IL1RAP,IRS1,ITGB5,KRAS,MAP4K4,NRAS,PLCB3,PLCD1,PLCD3,PRKAA2,PRKAB2,RALA,RAP2A,RAP2B,SHC1 |
| Chemokine Signaling | 1.37E+00 | 1.38E-01 | 2.111 | CCL11,CFL1,GNAI3,KRAS,LIMK1,NRAS,PLCB3,PPP1R12A,RALA,RAP2A,RAP2B |
| IL-2 Signaling | 1.37E+00 | 1.48E-01 | 2.828 | CSNK2A1,ELK1,KRAS,NRAS,PTPN11,RALA,RAP2A,RAP2B,SHC1 |
| Œ≥-linolenate Biosynthesis II (Animals) | 1.35E+00 | 2.22E-01 | 2 | ACSL3,FADS1,FADS2,SLC27A4 |
| D-myo-inositol (1,4,5)-Trisphosphate Biosynthesis | 1.34E+00 | 1.92E-01 | 1.342 | PIP4K2A,PIP4K2C,PLCB3,PLCD1,PLCD3 |

| | | | | |
|---|---|---|---|---|
| Gluconeogenesis I | 1.34E+00 | 1.92E-01 | 2.236 | ALDOA,ENO1,ENO2,GAPDH,GPI |
| Œ≥-glutamyl Cycle | 1.33E+00 | 2.73E-01 | NaN | GCLC,GCLM,GSS |
| GDP-glucose Biosynthesis | 1.33E+00 | 2.73E-01 | NaN | HK2,PGM2,PGM3 |
| HER-2 Signaling in Breast Cancer | 1.32E+00 | 1.12E-01 | 2.558 | ARF6,BCL2L11,BIRC5,CCNE1,CCNE2,COX6B2,ELK1,GSK3A,ITGB1,ITGB5,ITGB8,KPNB1,KRAS,NRAS,PARD3,PARD6G,RALA,RAP2A,RAP2B,RBL1,RPS6KB1,YES1 |
| Endocannabinoid Cancer Inhibition Pathway | 1.32E+00 | 1.19E-01 | -0.243 | CASP2,CCNE1,CCNE2,GNA13,GNAI3,HIF1A,NOS1,PGF,PRKAA2,PRKAB2,SMPD4,SNAI2,SPTLC1,TCF3,TRIB3,TWIST1,VEGFC |
| CDK5 Signaling | 1.31E+00 | 1.25E-01 | 1.732 | CDK5R1,ITGB1,KRAS,LAMB1,LAMC1,LAMC2,MAPK6,NRAS,PPP1CC,PPP1R12A,PPP2R1B,RALA,RAP2A,RAP2B |
