## Supplemental Table S3 for "Multiomic Characterization of Stage I Lung Adenocarcinoma Reveals Distinct Genetic and Immunologic Features of Recurrent Disease"

**Table S3. Pathway Analysis for Differential Gene Expression Heatmap Cluster 3**

| Ingenuity Canonical Pathways | -log(p-value) | Ratio | z-score | Molecules |
|---|---|---|---|---|
| Antigen Presentation Pathway | 1.03E+01 | 4.10E-01 | NaN | CD74,CIITA,HLA-DMA,HLA-DMB,HLA-DOA,HLA-DOB,HLA-DPA1,HLA-DPB1,HLA-DQA1,HLA-DQB1,HLA-DQB2,HLA-DRA,HLA-DRB1,HLA-DRB5,HLA-E,MR1 |
| Th1 and Th2 Activation Pathway | 9.15E+00 | 1.86E-01 | NaN | CCR4,CD40LG,HLA-DMA,HLA-DMB,HLA-DOA,HLA-DOB,HLA-DPA1,HLA-DPB1,HLA-DQA1,HLA-DQB1,HLA-DQB2,HLA-DRA,HLA-DRB1,HLA-DRB5,IKZF1,IL12B,IL24,IL33,IL6R,JUN,LTA,NFATC1,NFATC2,NFATC3,PIK3R1,PIK3R6,PRKCQ,PTGDR2,S1PR1,TGFBR2,TGFBR3,TSLP |
| CREB Signaling in Neurons | 8.94E+00 | 1.16E-01 | -7.878 | ACKR1,ADCY2,ADCY9,ADGRB3,ADGRF5,ADGRG2,ADRA2A,ADRB2,CACNA1D,CACNA2D2,CACNA2D3,CACNB4,CALCRL,CAMK2D,CAMK4,CCR2,CCR4,CCR6,CCR7,CNR2,CX3CR1,CYSLTR1,EDNRB,FGFR2,GHR,GNAL,GNAO1,GNG7,GPR146,GPR162,GPR20,GPR25,GPR55,GRIA1,GRID1,GRIK3,GRIK4,HGF,ITPR1,ITPR2,MCHR1,NGFR,NMUR1,NPY1R,NTRK2,NTRK3,P2RY12,P2RY13,P2RY14,PIK3R1,PIK3R6,PLCB2,PLCE1,PLCG2,PLCL1,PRKCB,PRKCE,PRKCQ,PTGDR2,PTGER4,PTGIR,PTH1R,RXFP1,S1PR1,S1PR4,SHC3,SSTR1,TGFBR2,TGFBR3,XCR1 |
| Cardiac Hypertrophy Signaling (Enhanced) | 8.61E+00 | 1.19E-01 | -7.211 | ADCY2,ADCY9,ADRA2A,ADRB2,AKAP13,CACNA1D,CACNA2D2,CACNA2D3,CACNB4,CAMK2D,CAMK4,CD40LG,EDNRB,ENPP6,FGF10,FGF14,FGFR2,GHR,GNG7,HSPB7,IL12B,IL13RA2,IL33,IL5RA,IL6R,IL6ST,ITGA10,ITGA8,ITGA9,ITGAL,ITPR1,ITPR2,JUN,LTA,LTB,MAP3K3,MAPK10,MEF2C,MYOCD,NFATC1,NFATC2,NFATC3,NGFR,PDE1B,PDE1C,PDE7B,PDE8B,PIK3R1,PIK3R6,PLCB2,PLCE1,PLCG2,PLCL1,PRKCB,PRKCE,PRKCQ,RPS6KA5,RYR2,TGFBR2,TGFBR3,TNFSF12,TNFSF13,WNT11,WNT2B |
| Th2 Pathway | 8.39E+00 | 1.97E-01 | -0.905 | CCR4,HLA-DMA,HLA-DMB,HLA-DOA,HLA-DOB,HLA-DPA1,HLA-DPB1,HLA-DQA1,HLA-DQB1,HLA-DQB2,HLA-DRA,HLA-DRB1,HLA-DRB5,IKZF1,IL12B,IL24,IL33,JUN,NFATC2,PIK3R1,PIK3R6,PRKCQ,PTGDR2,S1PR1,TGFBR2,TGFBR3,TSLP |
| B Cell Development | 7.53E+00 | 3.26E-01 | NaN | CD79B,HLA-DMA,HLA-DMB,HLA-DOA,HLA-DOB,HLA-DPA1,HLA-DPB1,HLA-DQA1,HLA-DQB1,HLA-DQB2,HLA-DRA,HLA-DRB1,HLA-DRB5,SPN |
| Neuroinflammation Signaling Pathway | 7.23E+00 | 1.33E-01 | -4 | AGER,BCL2,CD200R1,CX3CL1,CX3CR1,FOS,GRIA1,HLA-DMA,HLA-DMB,HLA-DOA,HLA-DOB,HLA-DPA1,HLA-DPB1,HLA-DQA1,HLA-DQB1,HLA-DQB2,HLA-DRA,HLA-DRB1,HLA-DRB5,HLA-E,IL12B,IL6R,JUN,KCNJ5,MAPK10,MMP9,MR1,NFATC1,NFATC2,NFATC3,PIK3R1,PIK3R6,PLCG2,S100B,SLC1A2,TGFBR2,TGFBR3,TLR10,TLR2,TLR3,TLR5,TLR7 |
| PD-1, PD-L1 cancer immunotherapy pathway | 6.71E+00 | 1.98E-01 | 3.638 | FBXO38,HLA-DMA,HLA-DMB,HLA-DOA,HLA-DOB,HLA-DPA1,HLA-DPB1,HLA-DQA1,HLA-DQB1,HLA-DQB2,HLA-DRA,HLA-DRB1,HLA-DRB5,HLA-E,IL12B,MR1,NGFR,PIK3R1,PIK3R6,PRKCQ,ZAP70 |
| IL-4 Signaling | 6.43E+00 | 2.07E-01 | NaN | FCER2,HLA-DMA,HLA-DMB,HLA-DOA,HLA-DOB,HLA-DPA1,HLA-DPB1,HLA-DQA1,HLA-DQB1,HLA-DQB2,HLA-DRA,HLA-DRB1,HLA-DRB5,NFATC1,NFATC2,NFATC3,NR3C2,PIK3R1,PIK3R6 |
| Protein Kinase A Signaling | 6.27E+00 | 1.17E-01 | -2.214 | ADCY2,ADCY9,AKAP13,AKAP14,AKAP6,AKAP7,CAMK2D,CAMK4,CNGA4,DUSP1,DUSP16,DUSP26,ENPP6,EPM2A,GNG7,H1-4,ITPR1,ITPR2,MYL3,NFATC1,NFATC2,NFATC3,NGFR,PDE1B,PDE1C,PDE7B,PDE8B,PHKB,PLCB2,PLCE1,PLCG2,PLCL1,PPP1R3C,PRKCB,PRKCE,PRKCQ,PTCH1,PTK2B,PTPN13,PTPN4,PTPRB,PTPRE,PTPRM,PTPRQ,PTPRT,RYR2,TGFBR2 |
| Th1 Pathway | 6.26E+00 | 1.80E-01 | -4 | CD40LG,HLA-DMA,HLA-DMB,HLA-DOA,HLA-DOB,HLA-DPA1,HLA-DPB1,HLA-DQA1,HLA-DQB1,HLA-DQB2,HLA-DRA,HLA-DRB1,HLA-DRB5,IL12B,IL6R,LTA,NFATC1,NFATC2,NFATC3,PIK3R1,PIK3R6,PRKCQ |
| Phagosome Formation | 5.30E+00 | 1.54E-01 | NaN | CR2,FCAMR,FCER1A,FCER2,ITGA10,ITGA8,ITGA9,ITGAL,PIK3R1,PIK3R6,PLCB2,PLCE1,PLCG2,PLCL1,PRKCB,PRKCE,PRKCQ,SCARA3,TLR10,TLR2,TLR3,TLR5,TLR7 |
| cAMP-mediated signaling | 5.19E+00 | 1.31E-01 | -4.315 | ADCY2,ADCY9,ADRA2A,ADRB2,AKAP13,AKAP14,AKAP6,AKAP7,CAMK2D,CAMK4,CCR4,CNGA4,CNR2,DUSP1,ENPP6,GNAL,GNAO1,NPY1R,P2RY12,P2RY13,P2RY14,PDE1B,PDE1C,PDE7B,PDE8B,PTGER4,PTGIR,PTH1R,S1PR1,XCR1 |
| Endocannabinoid Neuronal Synapse Pathway | 4.88E+00 | 1.50E-01 | -3.3 | ADCY2,ADCY9,CACNA1D,CACNA2D2,CACNA2D3,CACNB4,CNR2,DNAH10,DNALI1,GNAL,GNAO1,GNG7,GRIA1,ITPR1,ITPR2,KCNJ5,MAPK10,MGLL,PLCB2,PLCE1,PLCG2,PLCL1 |
| MSP-RON Signaling In Macrophages Pathway | 4.81E+00 | 1.62E-01 | 2.065 | CIITA,F11,FOS,HLA-DMA,HLA-DMB,HLA-DOA,HLA-DOB,HLA-DPA1,HLA-DPB1,HLA-DQA1,HLA-DQB1,HLA-DQB2,HLA-DRA,HLA-DRB1,HLA-DRB5,IL12B,JUN,PIK3R1,PIK3R6 |
| Systemic Lupus Erythematosus In B Cell Signaling Pathway | 4.76E+00 | 1.19E-01 | -5.048 | BCL2,BLNK,BTK,CAMK4,CD22,CD40LG,CD5,CD79B,FOS,IL12B,IL33,IL6R,IL6ST,JUN,LTA,LTB,NFATC1,NFATC2,NFATC3,PIK3R1,PIK3R6,PLCG2,PRKCB,PRKCE,PRKCQ,RASGRP2,SHC3,SHE,STING1,TLR3,TLR7,TNFSF12,TNFSF13 |
| Complement System | 4.74E+00 | 2.70E-01 | -0.816 | C3,C4A/C4B,C4BPA,C6,C7,C8B,CFD,CFI,CR2,MASP1 |
| Sperm Motility | 4.72E+00 | 1.22E-01 | -3.742 | BLK,BMX,BTK,CAMK4,CNGA4,EPHA3,ERBB4,FGFR2,FLT3LG,GNAL,GNAO1,GNG7,GUCY1A2,ITPR1,ITPR2,MUSK,NTRK2,NTRK3,PDE1B,PDE1C,PLCB2,PLCE1,PLCG2,PLCL1,PRKCB,PRKCE,PRKCQ,PTK2B,ROS1,TEK,ZAP70 |
| Crosstalk between Dendritic Cells and Natural Killer Cells | 4.58E+00 | 1.76E-01 | -3.873 | CAMK2D,CCR7,CD40LG,CD69,CD83,HLA-DRA,HLA-DRB1,HLA-DRB5,HLA-E,IL12B,ITGAL,LTA,LTB,NCR3,TLR3,TLR7 |
| Neuropathic Pain Signaling In Dorsal Horn Neurons | 4.57E+00 | 1.68E-01 | -4.123 | CAMK2D,CAMK4,FOS,GRIA1,ITPR1,ITPR2,KCNN3,NTRK2,PIK3R1,PIK3R6,PLCB2,PLCE1,PLCG2,PLCL1,PRKCB,PRKCE,PRKCQ |
| Adrenomedullin signaling pathway | 4.57E+00 | 1.31E-01 | -4.315 | ADCY2,ADCY9,BCL2,C3,CALCRL,CAMK4,FOS,GUCY1A2,IL33,ITPR1,ITPR2,KCNN3,LTA,MAPK10,PIK3R1,PIK3R6,PLCB2,PLCE1,PLCG2,PLCL1,PTK2B,RAMP2,RAMP3,RXRG,SHC3,SHE |
| PI3K Signaling in B Lymphocytes | 4.56E+00 | 1.47E-01 | -4.359 | BLK,BLNK,BTK,C3,CAMK2D,CAMK4,CD79B,CR2,FOS,ITPR1,ITPR2,JUN,NFATC1,NFATC2,NFATC3,PIK3R1,PLCB2,PLCE1,PLCG2,PLCL1,PRKCB |
| Endothelin-1 Signaling | 4.43E+00 | 1.31E-01 | -3.8 | ADCY2,ADCY9,CASQ1,EDNRB,FOS,GNAL,GNAO1,GPLD1,GUCY1A2,ITPR1,ITPR2,JUN,MAPK10,PIK3R1,PIK3R6,PLCB2,PLCE1,PLCG2,PLCL1,PLD4,PRKCB,PRKCE,PRKCQ,SHC3,SHE |
| Cellular Effects of Sildenafil (Viagra) | 4.29E+00 | 1.41E-01 | NaN | ADCY2,ADCY9,CACNA1D,CACNA2D2,CACNA2D3,CACNB4,CAMK4,GUCY1A2,ITPR1,ITPR2,KCNN3,MYH11,MYL3,PDE1B,PDE1C,PLCB2,PLCE1,PLCG2,PLCL1,PPP1R12B,SLC4A5 |

| | | | | |
|---|---|---|---|---|
| Corticotropin Releasing Hormone Signaling | 4.29E+00 | 1.41E-01 | -1.886 | ADCY2,ADCY9,CACNA1D,CACNA2D2,CACNA2D3,CACNB4,CAMK4,FOS,GNAO1,GUCY1A2,ITPR1,ITPR2,JUN,KRT1,MEF2C,PLCG2,PRKCB,PRKCE,PRKCQ,PTCH1,UCN3 |
| Breast Cancer Regulation by Stathmin1 | 4.20E+00 | 9.29E-02 | -6.26 | ACKR1,ADGRB3,ADGRF5,ADGRG2,ADRA2A,ADRB2,ANKHD1/ANKHD1-EIF4EBP3,ARHGEF15,ARHGEF18,ARHGEF6,ARHGEF9,CALCRL,CAMK2D,CAMK4,CCR2,CCR4,CCR6,CCR7,CNR2,CX3CR1,CYSLTR1,EDNRB,GNG7,GPR146,GPR162,GPR20,GPR25,GPR55,HGF,JUN,MCHR1,MMP9,NMUR1,NPY1R,P2RY12,P2RY13,P2RY14,PIK3R1,PIK3R6,PLCB2,PPP1R3C,PRKCB,PRKCE,PRKCQ,PTGDR2,PTGER4,PTGIR,PTH1R,RXFP1,S1PR1,S1PR4,SHC3,SSTR1,VEGFD,XCR1 |
| Role of Macrophages, Fibroblasts and Endothelial Cells in Rheumatoid Art | 4.09E+00 | 1.08E-01 | NaN | CAMK2D,CAMK4,CEBPA,FOS,FRZB,GNAO1,IL16,IL33,IL6R,IL6ST,JUN,LRP6,LTA,LTB,NFATC1,NFATC2,NFATC3,NGFR,PIK3R1,PIK3R6,PLCB2,PLCE1,PLCG2,PLCL1,PRKCB,PRKCE,PRKCQ,TLR10,TLR2,TLR3,TLR5,TLR7,VEGFD,WNT11,WNT2B |
| G-Protein Coupled Receptor Signaling | 4.03E+00 | 1.12E-01 | NaN | ADCY2,ADCY9,ADRA2A,ADRB2,CAMK2D,CAMK4,CCR4,CNR2,DUSP1,ENPP6,GNAL,GNAO1,NPY1R,P2RY12,P2RY13,P2RY14,PDE1B,PDE1C,PDE7B,PDE8B,PIK3R1,PIK3R6,PLCB2,PRKCB,PRKCE,PTGER4,PTGIR,PTH1R,PTK2B,S1PR1,XCR1 |
| Leukocyte Extravasation Signaling | 3.92E+00 | 1.24E-01 | -3.13 | ACTN2,ARHGAP6,BMX,BTK,CLDN18,CLDN2,DLC1,ITGAL,JAM2,MAPK10,MMP24,MMP28,MMP9,NCF4,PECAM1,PIK3R1,PIK3R6,PLCG2,PRKCB,PRKCE,PRKCQ,PTK2B,RASSF5,SPN |
| Role of NFAT in Cardiac Hypertrophy | 3.83E+00 | 1.18E-01 | -5 | ADCY2,ADCY9,CACNA1D,CACNA2D2,CACNA2D3,CACNB4,CAMK2D,CAMK4,GNG7,IL6ST,ITPR1,ITPR2,MAPK10,MEF2C,PIK3R1,PIK3R6,PLCB2,PLCE1,PLCG2,PLCL1,PRKCB,PRKCE,PRKCQ,SHC3,SLC8A3,TGFBR2 |
| Apelin Endothelial Signaling Pathway | 3.76E+00 | 1.37E-01 | -3.153 | ADCY2,ADCY9,ANGPT1,CAMK4,FOS,GNAL,GNAO1,GNG7,JUN,KLF2,MAPK10,MEF2C,PIK3R1,PIK3R6,PLCB2,PRKCB,PRKCE,PRKCQ,TEK |
| Œ±-Adrenergic Signaling | 3.76E+00 | 1.51E-01 | -3.162 | ADCY2,ADCY9,ADRA2A,CAMK4,EPM2A,GNAL,GNAO1,GNG7,ITPR1,ITPR2,PHKB,PLCG2,PRKCB,PRKCE,PRKCQ,SLC8A3 |
| Thrombin Signaling | 3.76E+00 | 1.19E-01 | -3.578 | ADCY2,ADCY9,ARHGEF15,ARHGEF6,ARHGEF9,CAMK2D,CAMK4,GATA5,GATA6,GNAL,GNAO1,GNG7,ITPR1,ITPR2,MYL3,PIK3R1,PIK3R6,PLCB2,PLCE1,PLCG2,PLCL1,PPP1R12B,PRKCB,PRKCE,PRKCQ |
| Opioid Signaling Pathway | 3.67E+00 | 1.09E-01 | -3.657 | ADCY2,ADCY9,ARRB2,BLK,CACNA1D,CACNA2D2,CACNA2D3,CACNB4,CAMK2D,CAMK4,FOS,FOSB,GNAL,GNAO1,GNG7,ITPR1,ITPR2,KCNJ5,PDE1B,PDE1C,PENK,PRKCB,PRKCE,PRKCQ,RGS13,RGS5,RGS9,RPS6KA5,RYR2,SCN7A |
| Nitric Oxide Signaling in the Cardiovascular System | 3.66E+00 | 1.43E-01 | -3.606 | CACNA1D,CACNA2D2,CACNA2D3,CACNB4,CAMK4,GUCY1A2,ITPR1,ITPR2,PDE1B,PDE1C,PIK3R1,PIK3R6,PRKCB,PRKCE,PRKCQ,RYR2,VEGFD |
| Granulocyte Adhesion and Diapedesis | 3.65E+00 | 1.22E-01 | NaN | CCL14,CCL17,CCL19,CCL22,CCL23,CCR2,CCR4,CCR6,CCR7,CLDN18,CLDN2,CSF3R,CX3CL1,CXCL14,CXCL16,IL33,MMP24,MMP28,MMP9,NGFR,PECAM1,SELL,SELP |
| Agranulocyte Adhesion and Diapedesis | 3.63E+00 | 1.17E-01 | NaN | AOC3,CCL14,CCL17,CCL19,CCL22,CCL23,CCR2,CCR4,CCR6,CCR7,CD34,CLDN18,CLDN2,CX3CL1,CXCL14,CXCL16,IL33,MMP24,MMP28,MMP9,MYH11,MYL3,PECAM1,SELL,SELP |
| Role of Pattern Recognition Receptors in Recognition of Bacteria and Viru | 3.55E+00 | 1.28E-01 | -3.464 | C3,CD40LG,IL12B,IL33,LTA,LTB,MAPK10,PIK3R1,PIK3R6,PLCG2,PRKCB,PRKCE,PRKCQ,RNASEL,TLR2,TLR3,TLR5,TLR7,TNFSF12,TNFSF13 |
| GŒ±q Signaling | 3.47E+00 | 1.24E-01 | -4.243 | BTK,CAMK4,EPM2A,GNAL,GNAO1,GNG7,GPLD1,ITPR1,ITPR2,NFATC1,NFATC2,NFATC3,PIK3R1,PIK3R6,PLCB2,PLCG2,PLD4,PRKCB,PRKCE,PRKCQ,PTK2B |
| Melatonin Signaling | 3.36E+00 | 1.67E-01 | -2.887 | CAMK2D,CAMK4,GNAO1,PLCB2,PLCE1,PLCG2,PLCL1,PRKCB,PRKCE,PRKCQ,RORA,RORB |
| Cardiac Œ≤-adrenergic Signaling | 3.33E+00 | 1.21E-01 | -2.333 | ADCY2,ADCY9,AKAP13,AKAP14,AKAP6,AKAP7,CACNA1D,CACNA2D2,CACNA2D3,CACNB4,ENPP6,GNAL,GNAO1,GNG7,PDE1B,PDE1C,PDE7B,PDE8B,PPP1R3C,RYR2,SLC8A3 |
| GPCR-Mediated Nutrient Sensing in Enteroendocrine Cells | 3.31E+00 | 1.38E-01 | -4 | ADCY2,ADCY9,CACNA1D,CACNA2D2,CACNA2D3,CACNB4,GNG7,ITPR1,ITPR2,PLCB2,PLCE1,PLCG2,PLCL1,PRKCB,PRKCE,PRKCQ |
| Factors Promoting Cardiogenesis in Vertebrates | 3.30E+00 | 1.26E-01 | -4.243 | BMP3,BMP5,CAMK2D,LRP6,MAPK10,MEF2C,MYOCD,PLCB2,PLCE1,PLCG2,PLCL1,PRKCB,PRKCE,PRKCQ,TBX5,TGFBR2,TGFBR3,WNT11,WNT2B |
| G Beta Gamma Signaling | 3.24E+00 | 1.32E-01 | -3.873 | ADCY2,ARHGEF6,BTK,CACNA1D,CACNA2D2,CACNA2D3,CACNB4,GNAL,GNAO1,GNG7,ITPR1,ITPR2,KCNJ5,PLCG2,PRKCB,PRKCE,PRKCQ |
| FcŒ≥RIIB Signaling in B Lymphocytes | 3.22E+00 | 1.53E-01 | -2.646 | BLNK,BTK,CACNA1D,CACNA2D2,CACNA2D3,CACNB4,CD79B,ITPR1,ITPR2,MAPK10,PIK3R1,PIK3R6,PLCG2 |
| Gustation Pathway | 3.16E+00 | 1.20E-01 | NaN | ADCY2,ADCY9,CACNA1D,CACNA2D2,CACNA2D3,CACNB4,ENPP6,GNG7,ITPR1,ITPR2,P2RX1,P2RY12,P2RY13,P2RY14,P2RY8,PDE1B,PDE1C,PDE7B,PDE8B,PLCB2 |
| Renin-Angiotensin Signaling | 3.14E+00 | 1.33E-01 | -3.742 | ADCY2,ADCY9,FOS,ITPR1,ITPR2,JUN,MAPK10,PIK3R1,PIK3R6,PLCG2,PRKCB,PRKCE,PRKCQ,PTK2B,SHC3,SHE |
| Dopamine-DARPP32 Feedback in cAMP Signaling | 3.11E+00 | 1.16E-01 | -3.357 | ADCY2,ADCY9,CACNA1D,CACNA2D2,CACNA2D3,CACNB4,CAMK4,GUCY1A2,ITPR1,ITPR2,KCNJ15,KCNJ16,KCNJ5,PLCB2,PLCE1,PLCG2,PLCL1,PPP1R3C,PRKCB,PRKCE,PRKCQ |
| Apelin Cardiomyocyte Signaling Pathway | 3.07E+00 | 1.41E-01 | -3.742 | CAT,ITPR1,MAPK10,MYL3,PIK3R1,PIK3R6,PLCB2,PLCE1,PLCG2,PLCL1,PRKCB,PRKCE,PRKCQ,SLC8A3 |
| Dilated Cardiomyopathy Signaling Pathway | 3.05E+00 | 1.23E-01 | -1.155 | ADCY2,ADCY9,BCL2,CACNA1D,CACNA2D2,CACNA2D3,CACNB4,CAMK2D,CAMK4,DES,DMD,ITPR1,ITPR2,MAP3K3,MYH11,MYL3,PRKCE,RYR2 |
| nNOS Signaling in Skeletal Muscle Cells | 3.03E+00 | 1.88E-01 | NaN | CACNA1D,CACNA2D2,CACNA2D3,CACNB4,CAMK4,DMD,ITPR1,ITPR2,RYR2 |
| IL-8 Signaling | 2.97E+00 | 1.09E-01 | -4.025 | ANGPT1,ARRB2,BCL2,CR2,FOS,GNAL,GNAO1,GNG7,GPLD1,JUN,MAPK10,MMP9,PIK3R1,PIK3R6,PLCB2,PLD4,PRKCB,PRKCE,PRKCQ,PTK2B,RAB11FIP2,TEK,VEGFD |
| Axonal Guidance Signaling | 2.89E+00 | 8.70E-02 | NaN | ADAMTS8,ARHGEF15,ARHGEF6,BMP3,BMP5,DPYSL2,EPHA3,GNAL,GNAO1,GNG7,HHIP,ITGA10,ITGA8,ITGA9,ITGAL,MMP24,MMP28,MMP9,MYL3,NFATC1,NFATC2,NFATC3,NGFR,NTNG1,NTRK2,NTRK3,PIK3R1,PIK3R6,PLCB2,PLCE1,PLCG2,PLCL1,PLXNA2,PRKCB,PRKCE,PRKCQ,PTCH1,RASSF5,ROBO2,SLIT2,SLIT3,VEGFD,WNT11,WNT2B |
| Amyotrophic Lateral Sclerosis Signaling | 2.88E+00 | 1.30E-01 | -1.508 | BCL2,CACNA1D,CACNA2D2,CACNA2D3,CACNB4,CAPN6,CAT,GRIA1,GRID1,GRIK3,GRIK4,PIK3R1,PIK3R6,SLC1A2,VEGFD |
| Aldosterone Signaling in Epithelial Cells | 2.86E+00 | 1.16E-01 | -3.606 | DNAJC27,DUSP1,HSPB6,HSPB7,HSPB8,ITPR1,ITPR2,KCNMB2,NR3C2,PIK3R1,PIK3R6,PIP5K1B,PLCB2,PLCE1,PLCG2,PLCL1,PRKCB,PRKCE,PRKCQ |
| Growth Hormone Signaling | 2.86E+00 | 1.55E-01 | -3.317 | A2M,CEBPA,FOS,GHR,PIK3R1,PIK3R6,PLCG2,PRKCB,PRKCE,PRKCQ,RPS6KA5 |
| April Mediated Signaling | 2.80E+00 | 1.90E-01 | -2.828 | FOS,JUN,MAPK10,NFATC1,NFATC2,NFATC3,TNFRSF13B,TNFSF13 |

| | | | | |
|---|---|---|---|---|
| UVB-Induced MAPK Signaling | 2.77E+00 | 1.73E-01 | -3 | FOS,JUN,MAPK10,PIK3R1,PIK3R6,PRKCB,PRKCE,PRKCQ,RPS6KA5 |
| Coagulation System | 2.63E+00 | 2.00E-01 | -0.378 | A2M,F10,F11,F8,PLG,SERPIND1,VWF |
| eNOS Signaling | 2.63E+00 | 1.13E-01 | -3.464 | ADCY2,ADCY9,AQP1,AQP4,AQP5,CAMK4,CHRNA6,CNGA4,GUCY1A2,ITPR1,ITPR2,PIK3R1,PIK3R6,PLCG2,PRKCB,PRKCE,PRKCQ,VEGFD |
| GDNF Family Ligand-Receptor Interactions | 2.61E+00 | 1.45E-01 | -3 | DOK6,FOS,GFRA1,GFRA2,ITPR1,ITPR2,JUN,MAPK10,PIK3R1,PIK3R6,PLCG2 |
| Signaling by Rho Family GTPases | 2.55E+00 | 9.70E-02 | -3.9 | ARHGEF15,ARHGEF18,ARHGEF6,ARHGEF9,CDH19,CDH23,DES,FOS,GNAL,GNAO1,GNG7,ITGA10,ITGA8,ITGA9,ITGAL,JUN,MAPK10,MYL3,PIK3R1,PIK3R6,PIP5K1B,PPP1R12B,PTK2B,SEPTIN1,SEPTIN4,WASF3 |
| GNRH Signaling | 2.53E+00 | 1.06E-01 | -3.873 | ADCY2,ADCY9,CACNA1D,CACNA2D2,CACNA2D3,CACNB4,CAMK2D,CAMK4,FOS,GNG7,ITPR1,ITPR2,JUN,MAP3K3,MAPK10,PLCB2,PRKCB,PRKCE,PRKCQ,PTK2B |
| Colorectal Cancer Metastasis Signaling | 2.53E+00 | 9.67E-02 | -4.264 | ADCY2,ADCY9,FOS,GNAL,GNAO1,GNG7,IL6R,IL6ST,JUN,LRP6,MAPK10,MMP24,MMP28,MMP9,PIK3R1,PIK3R6,PTGER4,TGFBR2,TLR10,TLR2,TLR3,TLR5,TLR7,VEGFD,WNT11,WNT2B |
| Cardiac Hypertrophy Signaling | 2.47E+00 | 9.69E-02 | -4.472 | ADCY2,ADCY9,ADRA2A,ADRB2,CACNA1D,CACNA2D2,CACNA2D3,CACNB4,CAMK4,GNAL,GNAO1,GNG7,IL6R,JUN,MAP3K3,MAPK10,MEF2C,MYL3,PIK3R1,PIK3R6,PLCB2,PLCE1,PLCG2,PLCL1,TGFBR2 |
| Type II Diabetes Mellitus Signaling | 2.46E+00 | 1.12E-01 | -2.828 | ACSBG1,ACSL5,ACSL6,CACNA1D,CACNA2D2,CACNA2D3,CACNB4,CD36,ITPR1,ITPR2,MAPK10,NGFR,PIK3R1,PIK3R6,PRKCB,PRKCE,PRKCQ |
| Relaxin Signaling | 2.43E+00 | 1.11E-01 | -2.53 | ADCY2,ADCY9,ENPP6,FOS,GNAL,GNAO1,GNG7,GUCY1A2,JUN,MMP9,PDE1B,PDE1C,PDE7B,PDE8B,PIK3R1,PIK3R6,RXFP1 |
| Glucocorticoid Receptor Signaling | 2.42E+00 | 8.09E-02 | NaN | A2M,ADRB2,AR,BCL2,CEBPA,DNAH10,DNALI1,DUSP1,FBP1,FOS,GHR,HLA-DMA,HLA-DMB,HLA-DOA,HLA-DOB,HLA-DPA1,HLA-DPB1,HLA-DQA1,HLA-DQB1,HLA-DQB2,HLA-DRA,HLA-DRB1,HLA-DRB5,HLA-E,HP,IL13RA2,IL5RA,IL6R,IL6ST,JUN,KAT2B,KRT1,MAPK10,MMP9,NFATC1,NFATC2,NFATC3,NR3C2,PGR,PIK3R1,PIK3R6,RPS6KA5,RXRG,SCGB1A1,SMARCA2,TGFBR2,TLR2 |
| CXCR4 Signaling | 2.40E+00 | 1.08E-01 | -2.673 | ADCY2,ADCY9,ELMO1,FOS,GNAL,GNAO1,GNG7,ITPR1,ITPR2,JUN,MAPK10,MYL3,PIK3R1,PIK3R6,PLCB2,PRKCB,PRKCE,PRKCQ |
| P2Y Purigenic Receptor Signaling Pathway | 2.39E+00 | 1.16E-01 | -3.357 | ADCY2,ADCY9,FOS,GNG7,JUN,P2RY12,PIK3R1,PIK3R6,PLCB2,PLCE1,PLCG2,PLCL1,PRKCB,PRKCE,PRKCQ |
| ErbB Signaling | 2.34E+00 | 1.28E-01 | -3.464 | ERBB4,FOS,JUN,MAPK10,NRG1,NRG2,PIK3R1,PIK3R6,PLCG2,PRKCB,PRKCE,PRKCQ |
| fMLP Signaling in Neutrophils | 2.33E+00 | 1.15E-01 | -3.606 | CAMK4,GNAL,GNAO1,GNG7,ITPR1,ITPR2,NFATC1,NFATC2,NFATC3,PIK3R1,PIK3R6,PLCB2,PRKCB,PRKCE,PRKCQ |
| HGF Signaling | 2.30E+00 | 1.14E-01 | -3.317 | FOS,HGF,ITGA10,ITGA8,ITGA9,ITGAL,JUN,MAP3K3,MAPK10,PIK3R1,PIK3R6,PLCG2,PRKCB,PRKCE,PRKCQ |
| Netrin Signaling | 2.29E+00 | 1.39E-01 | -2.828 | CACNA1D,CACNA2D2,CACNA2D3,CACNB4,ITPR1,ITPR2,NFATC1,NFATC2,NFATC3,RYR2 |
| UVA-Induced MAPK Signaling | 2.19E+00 | 1.22E-01 | -2.646 | FOS,JUN,MAPK10,PARP11,PARP15,PIK3R1,PIK3R6,PLCB2,PLCE1,PLCG2,PLCL1,RPS6KA5 |
| IL-15 Production | 2.19E+00 | 1.14E-01 | -3.742 | BLK,BMX,BTK,EPHA3,ERBB4,FGFR2,FLT3LG,MUSK,NTRK2,NTRK3,PTK2B,ROS1,TEK,ZAP70 |
| Calcium Signaling | 2.18E+00 | 9.72E-02 | -3.3 | CACNA1D,CACNA2D2,CACNA2D3,CACNB4,CAMK2D,CAMK4,CASQ1,CASQ2,CHRNA6,GRIA1,ITPR1,ITPR2,MEF2C,MYH11,MYL3,NFATC1,NFATC2,NFATC3,RYR2,SLC8A3,TRPC6 |
| GŒ±i Signaling | 2.12E+00 | 1.09E-01 | -2.309 | ADCY2,ADCY9,ADRA2A,CCR4,CNR2,GNAL,GNAO1,GNG7,NPY1R,P2RY12,P2RY13,P2RY14,S1PR1,SHC3,XCR1 |
| B Cell Activating Factor Signaling | 2.12E+00 | 1.63E-01 | -2.449 | FOS,JUN,MAPK10,NFATC1,NFATC2,NFATC3,TNFRSF13B |
| Regulation Of The Epithelial Mesenchymal Transition By Growth Factors P | 2.10E+00 | 9.90E-02 | -3.5 | CD40LG,FGF10,FGF14,FGFR2,FOS,HGF,IL6R,JUN,LTA,LTB,MAPK10,MMP9,NGFR,PIK3R1,PIK3R6,SHC3,TGFBR2,TNFSF12,TNFSF13 |
| Human Embryonic Stem Cell Pluripotency | 2.08E+00 | 1.02E-01 | NaN | BMP3,BMP5,FGFR2,GNAL,GNAO1,GNG7,LEFTY2,NTRK2,NTRK3,PIK3R1,PIK3R6,S1PR1,S1PR4,SMAD6,TGFBR2,WNT11,WNT2B |
| 4-aminobutyrate Degradation I | 2.08E+00 | 6.67E-01 | NaN | ABAT,ALDH5A1 |
| GP6 Signaling Pathway | 2.07E+00 | 1.10E-01 | -3.606 | BTK,CAMK4,COL6A5,COL6A6,GRAP2,ITPR1,LAMA2,PIK3R1,PIK3R6,PLCG2,PRKCB,PRKCE,PRKCQ,RASGRP2 |
| Synaptic Long Term Depression | 2.05E+00 | 9.79E-02 | -4.359 | CACNA1D,CACNA2D2,CACNA2D3,CACNB4,GNAL,GNAO1,GRIA1,GRID1,GUCY1A2,ITPR1,ITPR2,PLCB2,PLCE1,PLCG2,PLCL1,PRKCB,PRKCE,PRKCQ,RYR2 |
| B Cell Receptor Signaling | 2.05E+00 | 9.79E-02 | -4.359 | BLNK,BTK,CAMK2D,CAMK4,CD22,CD79B,JUN,MAP3K3,MEF2C,NFATC1,NFATC2,NFATC3,PIK3R1,PIK3R6,PLCG2,PRKCB,PRKCQ,PTK2B,RASSF5 |
| Inhibition of Angiogenesis by TSP1 | 2.05E+00 | 1.76E-01 | -2.236 | CD36,GUCY1A2,JUN,MAPK10,MMP9,TGFBR2 |
| Role of NFAT in Regulation of the Immune Response | 2.00E+00 | 7.76E-02 | -5.385 | BLNK,BTK,CAMK4,CD79B,FCER1A,FOS,GNAL,GNAO1,GNG7,HLA-DMA,HLA-DMB,HLA-DOA,HLA-DOB,HLA-DPA1,HLA-DPB1,HLA-DQA1,HLA-DQB1,HLA-DQB2,HLA-DRA,HLA-DRB1,HLA-DRB5,ITPR1,ITPR2,JUN,KPNA5,MEF2C,MS4A2,NFATC1,NFATC2,NFATC3,PIK3R1,PIK3R6,PLCB2,PLCG2,PRKCQ,TRAV8-2,TRAV8-4,TRAV8-6,TRAV9-2,TRBV19,TRBV29-1,TRBV5-1,TRBV6-1,ZAP70 |
| ErbB4 Signaling | 1.98E+00 | 1.32E-01 | -3 | ERBB4,NRG1,NRG2,PIK3R1,PIK3R6,PLCG2,PRKCB,PRKCE,PRKCQ |
| Airway Pathology in Chronic Obstructive Pulmonary Disease | 1.95E+00 | 1.10E-01 | NaN | CD40LG,CTSG,FGF10,FGF14,IL12B,IL33,LTA,LTB,MMP9,PTGDS,TNFSF12,TNFSF13,TSLP |
| GABA Receptor Signaling | 1.95E+00 | 1.07E-01 | NaN | ABAT,ADCY2,ADCY9,ALDH5A1,CACNA1D,CACNA2D2,CACNA2D3,CACNB4,GNAL,GNAO1,GNG7,ITPR1,ITPR2,KCNN3 |
| Acute Phase Response Signaling | 1.95E+00 | 9.73E-02 | -2.887 | A2M,C3,C4A/C4B,C4BPA,F8,FOS,HP,IL33,IL6R,IL6ST,ITIH3,JUN,NGFR,PIK3R1,PLG,RBP5,SERPIND1,VWF |
| Agrin Interactions at Neuromuscular Junction | 1.90E+00 | 1.29E-01 | -2.646 | ARHGEF6,ERBB4,JUN,LAMA2,MAPK10,MUSK,NRG1,NRG2,UTRN |
| RhoGDI Signaling | 1.90E+00 | 9.30E-02 | 2.309 | ARHGAP6,ARHGDIB,ARHGEF15,ARHGEF18,ARHGEF6,ARHGEF9,CDH19,CDH23,DLC1,GNAL,GNAO1,GNG7,ITGA10,ITGA8,ITGA9,ITGAL,MYH11,MYL3,PIP5K1B,PPP1R12B |
| Hepatic Cholestasis | 1.86E+00 | 9.52E-02 | NaN | ADCY2,ADCY9,CD40LG,CYP27A1,IL12B,IL33,JUN,LTA,LTB,MAPK10,NGFR,NR0B2,PRKCB,PRKCE,PRKCQ,SLCO3A1,TNFSF12,TNFSF13 |
| STAT3 Pathway | 1.84E+00 | 1.04E-01 | -3.162 | BCL2,FGFR2,GHR,HGF,IL13RA2,IL5RA,IL6R,IL6ST,MAPK10,NGFR,NTRK2,NTRK3,TGFBR2,TGFBR3 |
| White Adipose Tissue Browning Pathway | 1.82E+00 | 1.03E-01 | -2.887 | ADCY2,ADCY9,CACNA1D,CACNA2D2,CACNA2D3,CACNB4,FCER1A,FGFR2,GUCY1A2,ITPR1,ITPR2,MS4A2,NDN,RXRG |
| Erythropoietin Signaling Pathway | 1.82E+00 | 9.60E-02 | -0.5 | CD40LG,FOS,IL12B,IL33,ITPR1,ITPR2,JUN,LTA,LTB,PIK3R1,PIK3R6,PRKCB,PRKCE,PRKCQ,SHC3,TNFSF12,TNFSF13 |

| | | | | |
|---|---|---|---|---|
| Dendritic Cell Maturation | 1.81E+00 | 7.58E-02 | -5.916 | CCR7,CD1A,CD1B,CD1C,CD40LG,CD83,HLA-DMA,HLA-DMB,HLA-DOA,HLA-DOB,HLA-DPA1,HLA-DPB1,HLA-DQA1,HLA-DQB1,HLA-DQB2,HLA-DRA,HLA-DRB1,HLA-DRB5,HLA-E,IL12B,IL33,IRF8,LTA,LTB,MAPK10,MR1,NGFR,PIK3R1,PIK3R6,PLCB2,PLCE1,PLCG2,PLCL1,TLR2,TLR3,TRAV8-2,TRAV8-4,TRAV8-6,TRAV9-2,TRBV19,TRBV29-1,TRBV5-1,TRBV6-1 |
| PTEN Signaling | 1.81E+00 | 1.00E-01 | 3.162 | BCL2,FGFR2,GHR,ITGA10,ITGA8,ITGA9,ITGAL,MAGI3,NGFR,NTRK2,NTRK3,PIK3R1,PREX2,TGFBR2,TGFBR3 |
| GPCR-Mediated Integration of Enteroendocrine Signaling Exemplified by a | 1.79E+00 | 1.23E-01 | -0.333 | ADCY2,ADCY9,ADRB2,ITPR1,ITPR2,PLCB2,PLCE1,PLCG2,PLCL1 |
| Molecular Mechanisms of Cancer | 1.78E+00 | 7.87E-02 | NaN | ADCY2,ADCY9,ARHGEF15,ARHGEF18,ARHGEF6,ARHGEF9,BCL2,BMP3,BMP5,CAMK2D,FOS,GNAL,GNAO1,GNG7,ITGA10,ITGA8,ITGA9,ITGAL,JUN,LRP6,MAPK10,PIK3R1,PIK3R6,PLCB2,PRKCB,PRKCE,PRKCQ,PTCH1,RASGRF1,RBL2,SHC3,SMAD6,TGFBR2,WNT11,WNT2B |
| Leptin Signaling in Obesity | 1.75E+00 | 1.22E-01 | NaN | ADCY2,ADCY9,GHRL,PIK3R1,PIK3R6,PLCB2,PLCE1,PLCG2,PLCL1 |
| Role of Osteoblasts, Osteoclasts and Chondrocytes in Rheumatoid Arthritis | 1.72E+00 | 8.93E-02 | NaN | BCL2,BMP3,BMP5,CAMK4,FOS,FRZB,IL33,JUN,LRP6,MAPK10,NFATC1,NFATC2,NFATC3,NGFR,PIK3R1,PIK3R6,PTK2B,SMAD6,WNT11,WNT2B |
| Thrombopoietin Signaling | 1.71E+00 | 1.27E-01 | -2.828 | FOS,JUN,PIK3R1,PIK3R6,PLCG2,PRKCB,PRKCE,PRKCQ |
| Androgen Signaling | 1.69E+00 | 9.47E-02 | -3.317 | AR,CACNA1D,CACNA2D2,CACNA2D3,CACNB4,CAMK4,GNAL,GNAO1,GNG7,ITPR1,ITPR2,JUN,KAT2B,PRKCB,PRKCE,PRKCQ |
| Synaptic Long Term Potentiation | 1.66E+00 | 1.01E-01 | -2.887 | CAMK2D,CAMK4,GRIA1,ITPR1,ITPR2,PLCB2,PLCE1,PLCG2,PLCL1,PPP1R3C,PRKCB,PRKCE,PRKCQ |
| TREM1 Signaling | 1.65E+00 | 1.17E-01 | -3 | CD83,CIITA,NLRC3,PLCG2,TLR10,TLR2,TLR3,TLR5,TLR7 |
| Neuregulin Signaling | 1.62E+00 | 1.03E-01 | -2.828 | ERBB4,ITGA10,ITGA8,ITGA9,ITGAL,NRG1,NRG2,PIK3R1,PLCG2,PRKCB,PRKCE,PRKCQ |
| Toll-like Receptor Signaling | 1.62E+00 | 1.15E-01 | -2.449 | FOS,IL12B,IL33,JUN,TLR10,TLR2,TLR3,TLR5,TLR7 |
| RANK Signaling in Osteoclasts | 1.61E+00 | 1.10E-01 | -3 | CAMK4,FOS,JUN,MAP3K3,MAPK10,NFATC1,NFATC2,PIK3R1,PIK3R6,PTK2B |
| IL-17 Signaling | 1.61E+00 | 9.09E-02 | -3.638 | CCL17,CCL22,CD40LG,FOS,IL12B,IL33,JUN,LTA,LTB,MAPK10,MMP9,PIK3R1,PIK3R6,RGS13,TNFSF12,TNFSF13,VEGFD |
| Nicotine Degradation II | 1.60E+00 | 1.21E-01 | -2.828 | B4GAT1,CYP2U1,CYP4X1,FMO2,FMO3,FMO4,FMO5,INMT |
| Sphingosine-1-phosphate Signaling | 1.59E+00 | 1.02E-01 | -1.732 | ADCY2,ADCY9,CASQ1,PIK3R1,PIK3R6,PLCB2,PLCE1,PLCG2,PLCL1,PTK2B,S1PR1,S1PR4 |
| Glutamate Degradation III (via 4-aminobutyrate) | 1.59E+00 | 4.00E-01 | NaN | ABAT,ALDH5A1 |
| Cholecystokinin/Gastrin-mediated Signaling | 1.57E+00 | 1.01E-01 | -3.464 | FOS,IL33,ITPR1,ITPR2,JUN,MAPK10,MEF2C,PLCB2,PRKCB,PRKCE,PRKCQ,PTK2B |
| CD40 Signaling | 1.57E+00 | 1.19E-01 | -2.646 | CD40LG,FCER2,FOS,JUN,LTA,MAPK10,PIK3R1,PIK3R6 |
| Chemokine Signaling | 1.55E+00 | 1.13E-01 | -2.333 | CAMK2D,CAMK4,FOS,JUN,PLCB2,PLCG2,PPP1R12B,PRKCB,PTK2B |
| G Protein Signaling Mediated by Tubby | 1.53E+00 | 1.36E-01 | NaN | GNAL,GNAO1,GNG7,PLCB2,PLCG2,TUB |
| CCR3 Signaling in Eosinophils | 1.51E+00 | 9.63E-02 | -2.449 | CAMK4,GNAL,GNAO1,GNG7,ITPR1,ITPR2,PIK3R1,PIK3R6,PLCB2,PPP1R12B,PRKCB,PRKCE,PRKCQ |
| Estrogen Receptor Signaling | 1.51E+00 | 7.67E-02 | -4.536 | ADCY2,ADCY9,BCL2,CACNA1D,CACNA2D2,CACNA2D3,CACNB4,FOS,GNAL,GNAO1,GNG7,JUN,MMP24,MMP28,MMP9,MYL3,NR0B2,PGR,PIK3R1,PIK3R6,PLCB2,PLCE1,PLCG2,PLCL1,PPP1R12B,PRKCB,PRKCE,PRKCQ,SHC3,SHE,VEGFD |
| Senescence Pathway | 1.50E+00 | 8.08E-02 | -3.71 | CACNA1D,CACNA2D2,CACNA2D3,CACNB4,CAMK4,CAPN6,CAT,CBX7,ITPR2,JUN,KAT2B,NFATC1,NFATC2,NFATC3,PIK3R1,PIK3R6,RASSF5,RBL2,RPS6KA5,SMAD6,STING1,TGFBR2,TGFBR3,TLR2 |
| LPS/IL-1 Mediated Inhibition of RXR Function | 1.49E+00 | 8.33E-02 | -1.342 | ACSBG1,ACSL5,ACSL6,ALDH5A1,CAT,CES2,CHST7,CYP2U1,FABP4,FMO2,FMO3,FMO4,FMO5,GSTM5,IL33,JUN,MAOB,NGFR,NR0B2,SULT1C4,UST |
| IL-12 Signaling and Production in Macrophages | 1.49E+00 | 9.56E-02 | NaN | ALOX15,CD40LG,FOS,IL12B,IRF8,JUN,MAPK10,PIK3R1,PIK3R6,PRKCB,PRKCE,PRKCQ,TLR2 |
| Insulin Secretion Signaling Pathway | 1.48E+00 | 8.21E-02 | -3.9 | ADCY2,ADCY9,CACNA1D,CACNA2D2,CACNA2D3,CACNB4,CAMK2D,GHR,ITPR1,ITPR2,KCNB1,PIK3R1,PIK3R6,PLCB2,PLCE1,PLCG2,PLCL1,PRKCB,PRKCE,PRKCQ,RPS6KA5,RYR2 |
| PKCŒ∏ Signaling in T Lymphocytes | 1.44E+00 | 7.28E-02 | -4.69 | CACNA1D,CACNA2D2,CACNA2D3,CACNB4,CAMK2D,FOS,GRAP2,HLA-DMA,HLA-DMB,HLA-DOA,HLA-DOB,HLA-DPA1,HLA-DPB1,HLA-DQA1,HLA-DQB1,HLA-DQB2,HLA-DRA,HLA-DRB1,HLA-DRB5,ITPR1,ITPR2,JUN,MAP3K3,NFATC1,NFATC2,NFATC3,PIK3R1,PIK3R6,PLCG2,PRKCQ,TRAV8-2,TRAV8-4,TRAV8-6,TRAV9-2,TRBV19,TRBV29-1,TRBV5-1,TRBV6-1,ZAP70 |
| Fatty Acid Activation | 1.44E+00 | 2.14E-01 | NaN | ACSBG1,ACSL5,ACSL6 |
| RAR Activation | 1.44E+00 | 8.67E-02 | NaN | ADCY2,ADCY9,ADH1B,DUSP1,FOS,JUN,KAT2B,MAPK10,NRIP2,PIK3R1,PRKCB,PRKCE,PRKCQ,RBP5,RXRG,SMAD6,SMARCA2 |
| HMGB1 Signaling | 1.44E+00 | 8.98E-02 | -3.162 | AGER,CD40LG,FOS,IL12B,IL33,JUN,KAT2B,LTA,LTB,MAPK10,NGFR,PIK3R1,PIK3R6,TNFSF12,TNFSF13 |
| 14-3-3-mediated Signaling | 1.38E+00 | 9.45E-02 | -3.464 | FOS,JUN,MAPK10,PIK3R1,PIK3R6,PLCB2,PLCE1,PLCG2,PLCL1,PRKCB,PRKCE,PRKCQ |
| Prolactin Signaling | 1.38E+00 | 1.05E-01 | -2.828 | FOS,JUN,KCNMB2,PIK3R1,PIK3R6,PLCG2,PRKCB,PRKCE,PRKCQ |
| D-myo-inositol (1,4,5)-Trisphosphate Biosynthesis | 1.31E+00 | 1.54E-01 | -2 | PIP5K1B,PLCB2,PLCE1,PLCG2 |
