## Supplemental Table S4 for "Multiomic Characterization of Stage I Lung Adenocarcinoma Reveals Distinct Genetic and Immunologic Features of Recurrent Disease"

**Table S4 - Perturbed Biological Pathways in Recurrence High- vs. Low-Risk Stage I Lung Adenocarcinomas by Topology-Based Enrichment Analysis**

| Pathway | pSize | NDE | pNDE | tA | pPERT | pG | pGFdr | pGFWER | Status |
|---|---|---|---|---|---|---|---|---|---|
| Vascular smooth muscle contraction | 126 | 111 | 1.24E-31 | -58.38827702 | 5.00E-06 | 5.23E-35 | 1.94E-34 | 1.07E-32 | Inhibited |
| MicroRNAs in cancer | 230 | 146 | 3.25E-15 | -45.29966781 | 5.00E-06 | 7.58E-19 | 1.24E-18 | 1.55E-16 | Inhibited |
| Pathways in cancer | 470 | 422 | 2.14E-123 | 79.11631369 | 0.001 | 6.21E-124 | 1.27E-121 | 1.27E-121 | Activated |
| Human cytomegalovirus infection | 217 | 194 | 1.25E-56 | 88.78023538 | 0.001 | 1.71E-57 | 2.91E-56 | 3.49E-55 | Activated |
| Tight junction | 169 | 149 | 4.36E-42 | -42.60929075 | 0.001 | 4.50E-43 | 3.06E-42 | 9.18E-41 | Inhibited |
| Central carbon metabolism in cancer | 64 | 58 | 1.57E-18 | 43.23785372 | 0.001 | 7.66E-20 | 1.28E-19 | 1.56E-17 | Activated |
| Dilated cardiomyopathy (DCM) | 78 | 66 | 2.31E-17 | -28.23172681 | 0.001 | 1.07E-18 | 1.73E-18 | 2.18E-16 | Inhibited |
| Bile secretion | 30 | 22 | 8.64E-05 | -28.10858297 | 0.001 | 1.49E-06 | 1.65E-06 | 0.000304426 | Inhibited |
| Regulation of actin cytoskeleton | 214 | 188 | 3.36E-52 | 62.77732596 | 0.002 | 8.45E-53 | 1.08E-51 | 1.72E-50 | Activated |
| Cellular senescence | 147 | 139 | 2.02E-48 | -51.5404267 | 0.002 | 4.73E-49 | 4.82E-48 | 9.65E-47 | Inhibited |
| Cytokine-cytokine receptor interaction | 283 | 218 | 3.42E-41 | -21.53624554 | 0.002 | 6.88E-42 | 4.13E-41 | 1.40E-39 | Inhibited |
| Shigellosis | 223 | 206 | 3.39E-66 | 43.45722655 | 0.003 | 1.60E-66 | 3.64E-65 | 3.27E-64 | Activated |
| Ras signaling pathway | 228 | 205 | 8.99E-61 | -87.87430295 | 0.003 | 3.91E-61 | 7.26E-60 | 7.98E-59 | Inhibited |
| Ovarian steroidogenesis | 41 | 32 | 1.85E-07 | -29.36515716 | 0.003 | 1.24E-08 | 1.45E-08 | 2.53E-06 | Inhibited |
| Bacterial invasion of epithelial cells | 52 | 50 | 4.25E-19 | 39.19924354 | 0.004 | 8.30E-20 | 1.38E-19 | 1.69E-17 | Activated |
| GABAergic synapse | 66 | 51 | 7.96E-11 | 18.4672492 | 0.004 | 9.48E-12 | 1.28E-11 | 1.93E-09 | Activated |
| Small cell lung cancer | 91 | 91 | 3.89E-39 | 24.7215063 | 0.005 | 1.84E-39 | 9.89E-39 | 3.76E-37 | Activated |
| Amphetamine addiction | 63 | 53 | 5.44E-14 | -18.0023855 | 0.006 | 1.20E-14 | 1.73E-14 | 2.44E-12 | Inhibited |
| Toll-like receptor signaling pathway | 102 | 85 | 4.55E-21 | 59.34678208 | 0.007 | 1.68E-21 | 3.09E-21 | 3.43E-19 | Activated |
| Alzheimer disease | 240 | 220 | 5.57E-69 | -53.46304293 | 0.008 | 7.27E-69 | 2.12E-67 | 1.48E-66 | Inhibited |
| Estrogen signaling pathway | 126 | 100 | 1.99E-21 | 114.0503422 | 0.008 | 8.53E-22 | 1.60E-21 | 1.74E-19 | Activated |
| Kaposi sarcoma-associated herpesvirus infection | 162 | 141 | 1.83E-38 | 56.48316212 | 0.009 | 1.53E-38 | 7.61E-38 | 3.12E-36 | Activated |
| Neuroactive ligand-receptor interaction | 189 | 91 | 0.002513082 | -13.55541212 | 0.01 | 0.000291302 | 0.000306317 | 0.059425561 | Inhibited |
| Human T-cell leukemia virus 1 infection | 182 | 176 | 1.15E-65 | -33.6308446 | 0.014 | 2.49E-65 | 5.08E-64 | 5.08E-63 | Inhibited |
| Fanconi anemia pathway | 40 | 38 | 2.92E-14 | 6.930176609 | 0.017 | 1.80E-14 | 2.59E-14 | 3.67E-12 | Activated |
| Ferroptosis | 13 | 12 | 7.38E-05 | -5.831261782 | 0.017 | 1.83E-05 | 2.01E-05 | 0.003733879 | Inhibited |
| Legionellosis | 41 | 41 | 5.10E-18 | 17.86505779 | 0.02 | 4.56E-18 | 7.21E-18 | 9.30E-16 | Activated |
| Hepatitis C | 128 | 112 | 2.66E-31 | 36.65526896 | 0.022 | 4.40E-31 | 1.23E-30 | 8.97E-29 | Activated |
| Tuberculosis | 170 | 151 | 1.49E-43 | 41.14496513 | 0.024 | 3.69E-43 | 2.69E-42 | 7.54E-41 | Activated |
| Systemic lupus erythematosus | 18 | 18 | 2.57E-08 | -10.49341308 | 0.025 | 1.43E-08 | 1.66E-08 | 2.91E-06 | Inhibited |
| PPAR signaling pathway | 69 | 57 | 3.01E-14 | -12.59719624 | 0.026 | 2.80E-14 | 4.00E-14 | 5.72E-12 | Inhibited |
| FoxO signaling pathway | 123 | 114 | 1.42E-37 | 30.54527262 | 0.032 | 4.06E-37 | 1.88E-36 | 8.27E-35 | Activated |
| Sphingolipid signaling pathway | 96 | 94 | 3.75E-37 | 30.69862354 | 0.032 | 1.06E-36 | 4.70E-36 | 2.16E-34 | Activated |
| Hedgehog signaling pathway | 49 | 47 | 6.97E-18 | 4.954323919 | 0.032 | 9.81E-18 | 1.52E-17 | 2.00E-15 | Activated |
| Complement and coagulation cascades | 55 | 48 | 4.51E-14 | 7.661853258 | 0.035 | 5.53E-14 | 7.79E-14 | 1.13E-11 | Activated |
| Epstein-Barr virus infection | 162 | 145 | 8.22E-43 | 92.81298127 | 0.038 | 3.16E-42 | 1.95E-41 | 6.45E-40 | Activated |
| Intestinal immune network for IgA production | 27 | 25 | 4.08E-09 | -4.4799927 | 0.038 | 3.66E-09 | 4.36E-09 | 7.46E-07 | Inhibited |
| Cytosolic DNA-sensing pathway | 43 | 29 | 8.16E-05 | 11.48007474 | 0.04 | 4.45E-05 | 4.83E-05 | 0.009073866 | Activated |
| Salivary secretion | 48 | 41 | 1.49E-11 | -11.42482948 | 0.041 | 1.78E-11 | 2.35E-11 | 3.64E-09 | Inhibited |
| Salmonella infection | 201 | 193 | 3.46E-70 | 64.34851789 | 0.047 | 2.67E-69 | 9.07E-68 | 5.44E-67 | Activated |
| Adrenergic signaling in cardiomyocytes | 149 | 130 | 8.13E-36 | -23.48307348 | 0.048 | 3.31E-35 | 1.30E-34 | 6.75E-33 | Inhibited |
