## Supplemental Table S5 for "Multiomic Characterization of Stage I Lung Adenocarcinoma Reveals Distinct Genetic and Immunologic Features of Recurrent Disease"

**Table S5. Pathways Represented by Differentially Methylated Genes in Recurrence High- vs. Low-Risk Stage I Lung Adenocarcinomas**

| GO | ONTOLOGY | Description | N | DE | P.DE | FDR |
|---|---|---|---|---|---|---|
| GO:0030883 | MF | endogenous lipid antigen binding | 5 | 4 | 2.30E-06 | 1.74E-02 |
| GO:0030884 | MF | exogenous lipid antigen binding | 5 | 4 | 2.30E-06 | 1.74E-02 |
| GO:0048006 | BP | antigen processing and presentation, endogenous lipid antigen via MHC class Ib | 5 | 4 | 2.30E-06 | 1.74E-02 |
| GO:0030882 | MF | lipid antigen binding | 6 | 4 | 7.61E-06 | 4.32E-02 |
| GO:0030029 | BP | actin filament-based process | 795 | 87 | 4.12E-05 | 1.87E-01 |
| GO:0071723 | MF | lipopeptide binding | 10 | 4 | 7.61E-05 | 2.88E-01 |
| GO:0007154 | BP | cell communication | 6312 | 427 | 1.41E-04 | 3.59E-01 |
| GO:0050832 | BP | defense response to fungus | 36 | 7 | 1.51E-04 | 3.59E-01 |
| GO:0005886 | CC | plasma membrane | 5112 | 349 | 1.81E-04 | 3.59E-01 |
| GO:0098590 | CC | plasma membrane region | 1224 | 113 | 2.09E-04 | 3.59E-01 |

| KEGG | Description | N | DE | P.DE | FDR |
|---|---|---|---|---|---|
| path:hsa05146 | Amoebiasis | 100 | 18 | 2.56E-05 | 0.008719027 |
| path:hsa05131 | Shigellosis | 239 | 29 | 2.08E-04 | 0.035444957 |
| path:hsa04510 | Focal adhesion | 200 | 28 | 1.30E-03 | 0.114595143 |
| path:hsa04976 | Bile secretion | 81 | 12 | 0.001513946 | 0.114595143 |
| path:hsa05230 | Central carbon metabolism in cancer | 70 | 13 | 0.001680281 | 0.114595143 |
| path:hsa05135 | Yersinia infection | 134 | 17 | 0.004363642 | 0.185250164 |
| path:hsa04911 | Insulin secretion | 86 | 14 | 0.005282931 | 0.185250164 |
| path:hsa04918 | Thyroid hormone synthesis | 75 | 12 | 0.005392863 | 0.185250164 |
| path:hsa04930 | Type II diabetes mellitus | 46 | 10 | 0.005471343 | 0.185250164 |
| path:hsa04931 | Insulin resistance | 108 | 15 | 0.00558973 | 0.185250164 |

| MSigDB HALLMARK | Description | N | DE | P.DE | FDR |
|---|---|---|---|---|---|
| | HALLMARK_ALLOGRAFT_REJECTION | 199 | 39 | 0.001634497 | 0.039865048 |
| | HALLMARK_KRAS_SIGNALING_DN | 198 | 42 | 0.001937179 | 0.039865048 |
| | HALLMARK_MYOGENESIS | 200 | 47 | 0.002391903 | 0.039865048 |
| | HALLMARK_ESTROGEN_RESPONSE_LATE | 200 | 38 | 0.026081956 | 0.326024447 |
| | HALLMARK_INFLAMMATORY_RESPONSE | 199 | 31 | 0.03455946 | 0.345594598 |
| | HALLMARK_PANCREAS_BETA_CELLS | 40 | 9 | 0.066437634 | 0.553646949 |
| | HALLMARK_TNFA_SIGNALING_VIA_NFKB | 200 | 33 | 0.110477721 | 0.78912658 |
| | HALLMARK_ESTROGEN_RESPONSE_EARLY | 198 | 37 | 0.14697513 | 0.918594562 |
| | HALLMARK_SPERMATOGENESIS | 135 | 19 | 0.238967512 | 0.999976282 |
| | HALLMARK_REACTIVE_OXYGEN_SPECIES_PATHWAY | 49 | 8 | 0.267207436 | 0.999976282 |
